# A kernel integral method to remove biases in estimating trait turnover

**DOI:** 10.1101/2023.03.28.534538

**Authors:** Guillaume Latombe, Paul Boittiaux, Cang Hui, Melodie McGeoch

## Abstract

1. Trait diversity, including trait turnover, that differentiates the roles of species and communities according to their functions, is a fundamental component of biodiversity. Accurately capturing trait diversity is crucial to better understand and predict community assembly, as well as the consequences of global change on community resilience. Existing methods to compute trait turnover have limitations. Trait space approaches based on minimum convex polygons only consider species with extreme trait values. Tree-based approaches using dendrograms consider all species but distort trait distance between species. More recent trait space methods using complex polytopes try to harmonise the advantages of both methods, but their current implementation have mathematical flaws.
2. We propose a new kernel integral method (KIM) to compute trait turnover, based on the integration of kernel density estimators (KDEs) rather than using polytopes. We explore how this difference and the computational aspects of the KDE computation can influence the estimates of trait turnover. We compare our novel method to existing ones using justified theoretical expectations for a large number of simulations in which we control the number of species and the distribution of their traits. We illustrate the practical application of KIM using plant species introduced to the Pacific Islands of French Polynesia.
3. Analyses on simulated data show that KIM generates results better aligned with theoretical expectations than other methods and is less sensitive to the total number of species. Analyses for French Polynesia data also show that different methods can lead to different conclusions about trait turnover, and that the choice of method should be carefully considered based on the research question.
4. Mathematical aspects for computing trait turnover are crucial as they can have important effects on the results and therefore lead to different conclusions. Our novel kernel integral method generates values that better reflect the distribution of species in the trait space than other existing methods. We therefore recommend using KIM in future studies on trait turnover. In contrast, tree-based approaches should be kept for phylogenetic diversity, as phylogenetic trees will then reflect the constrained speciation process.

## 1. Introduction

Biodiversity is a complex concept and can most easily be quantified by distinguishing three complementary facets: taxonomic diversity based on a site-by-species matrix that captures the compositional properties of a community; phylogenetic diversity that captures the evolutionary relatedness among community members, using phylogenetic distance between species alongside the site-by-species matrix; and trait diversity that describes a community according to the traits of its residing species, using a species-by-trait matrix alongside the site-by-species matrix (Devictor et al., 2010). The study of functional traits has been advocated as fundamental to better understand and quantify community assembly (McGill et al., 2006), as well as the impact of global change on community resilience and on the ecosystem services that biodiversity provides (Gross et al., 2017). For example, through comparison with null models and by relating traits to environmental gradients and to each other, trait diversity can provide information about the assembly processes structuring an ecological community (Ackerly & Cornwell, 2007), including biotic interactions between species (Laureto et al., 2015). It also enables the estimation of components of ecosystem function, such as nutrient use and storage, or ecosystem productivity (Cadotte et al., 2011; Hillebrand & Matthiessen, 2009).

In addition to the decomposition of biodiversity into taxonomic, trait and phylogenetic components, unravelling how biodiversity is organised requires an understanding of how assemblages of species are more or less similar to one another at different places and times, i.e. turnover (Anderson et al., 2011). To do so, beta (β) diversity provides a direct link between biodiversity at the regional (gamma – γ – diversity) and local (alpha – α – diversity) scales (Anderson et al., 2011; Chao et al., 2005, 2019). In particular, taxonomic β diversity has been shown to be important for assessing the effects of conservation actions (Socolar et al., 2016), for example for estimating the effect of the spatial distribution of protected areas and their subdivision into multiple subareas on species diversity (Deane et al., 2022), or for extrapolating regional species richness from limited data (Kunin et al., 2018). Although having received less attention than taxonomic β diversity, trait turnover that describes change in trait diversity across communities or regions has also been measured using β diversity for similar applications (e.g. Carmona et al., 2012; Loiseau et al., 2017; Siefert et al., 2013; Swenson et al., 2012; Villéger et al., 2013).

As a valuable and increasingly measured biodiversity facet, there are multiple important steps to consider when estimating trait turnover over space or time. First, the precise choice of traits can substantially influence the outcome (Petchey & Gaston, 2006). Second, despite recent initiatives to collate large amounts of data for multiple traits across species (e.g. Kattge et al., 2020; Middleton-Welling et al., 2020; Tobias et al., 2022), trait data are still missing for many species and types of traits across taxonomic groups. Finally, and also the focus of this work, different mathematical methods exist to compute trait diversity and turnover that differ in outcome and therefore in the conclusions drawn in biodiversity studies (Loiseau et al., 2017; Sobral et al., 2016; Villéger et al., 2017). A systematic comparison can help identify an informative robust method and establish standards for quantifying trait turnover.

There are two main categories of methods for calculating trait β diversity: (i) methods based on the concept of trait space (referred to here as the ‘trait space approach’, and (ii) methods that use dendrograms (referred to here as the ‘tree-based approach’). The trait space approach is based on a multi-dimensional space whose axes are determined by the traits included in the analyses. Axes can correspond directly to the original traits or can be derived from these traits through ordinations to reduce dimensionality. A particular species typically represented as a single point in this trait space, and a polytope is computed as the trait envelope of a set of points representing the species of a community or assemblage. The minimum convex polytope (MCP), a convex hull, that encompasses all species of a community in the trait space (Figure 1), has traditionally been used in these analyses (Loiseau et al., 2017). As the MCP only captures information about the species with extreme trait values in a community, it is sensitive to outliers and ignores how species are distributed in the trait space, which can be crucial to delineate the functional roles of different species within an ecosystem (Mouillot et al., 2021). Although other hull methods can be used to compute the trait envelope (e.g. Irl et al., 2017), they are typically computationally intensive and have been seldomly applied to β diversity analyses.

**Figure 1.**
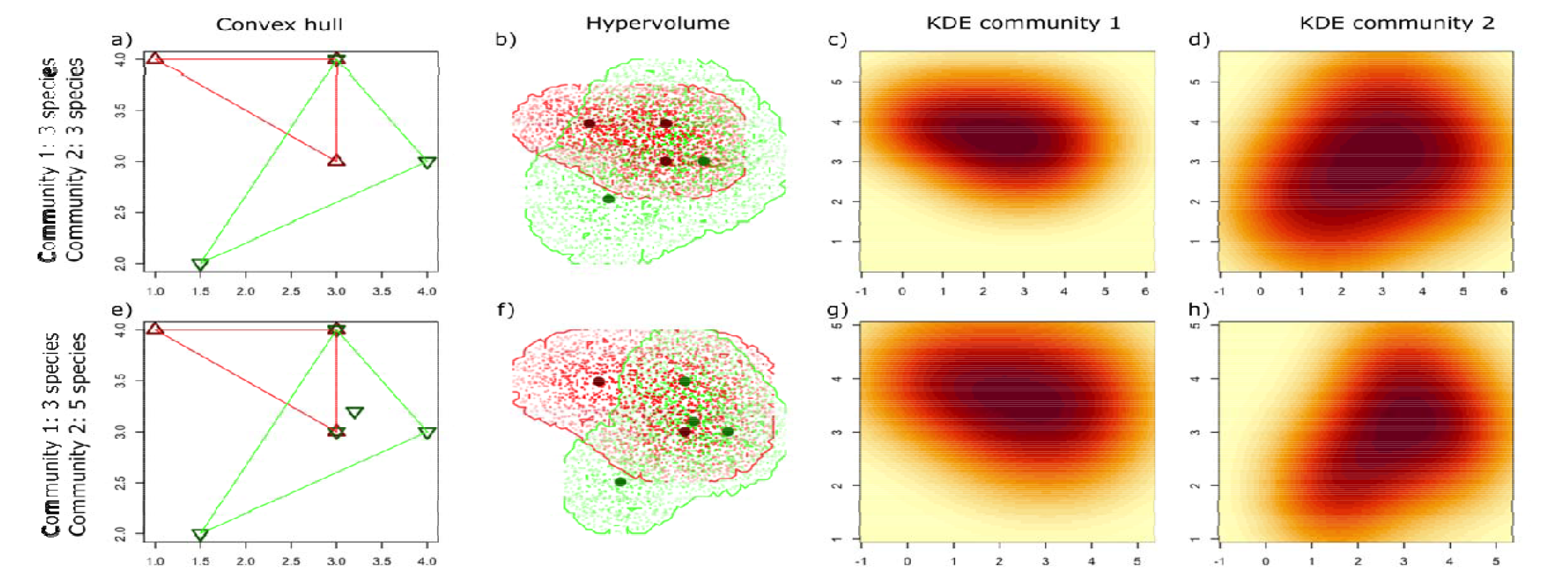
Summary of the trait space approaches for two communities with different species in a two-dimensional trait space. a,e) The convex hull remains the same irrespective of the additional species present in community 2, resulting in the same outcome when computing β diversity metrics. b,f) The KDH (kernel density hypervolume) method generates a polytope for each community, whose shape will vary with the absence or presence of the additional species in community 2 and is often non-convex. As a result, the outcome of the Jaccard dissimilarity or the Williams replacement formulas will differ. c,d) KDEs corresponding to the polytopes in (b). g,h) KDEs corresponding to the polytopes in (f).

The tree-based approach consists of computing all pairwise distance between species based on a set of traits, typically using the Gower distance to incorporate both continuous and discrete traits. A clustering algorithm is then applied to these distances to generate a dendrogram, from which measures of β diversity can be computed. Although the tree-based approach considers all species in the computation of trait turnover, the dendrogram splits into successive branches, and using the length of the branches connecting two species as a measure of distance distorts the original trait distance between them compared to the distance obtained through ordination in the trait space. In addition, the choice of the clustering algorithm for generating the dendrogram will inevitably influence the outcome (Loiseau et al., 2017).

The convex hull of trait space and the tree-based approach therefore make different computational trade-offs, and the appropriateness of the two approaches for measuring trait β diversity has been debated (Loiseau et al., 2017). In response to this debate and to incorporate information from all species, Mammola & Cardoso (2020) proposed another trait space approach where polytopes are defined by applying a threshold to the kernel density estimation (KDE; Figure 1; see details in Methods below). The resulting polytope is typically not convex, and its shape better reflects the distribution of species in the trait space. Although it has the potential to provide a more accurate estimate of trait diversity than the other two methods, this has not been formally assessed. The computational aspects when computing kernel densities have largely been overlooked. These aspects, as we plan to show here, are crucial so that all species contribute to β diversity in the communities.

Here we propose a new trait space method, which we term the kernel integral method (KIM), for computing trait β diversity based directly on the integration of the KDE rather than on the polytope. We explore how the computational aspects of the KDE computation can influence the estimates of trait β diversity with different methods. For comparison of the existing and new methods, we use a set of theoretical examples for which we can justify how trait β diversity metric should behave. We further apply the KIM method to compute non-native plant trait turnover across islands and archipelagos of the Pacific Islands of French Polynesia and compare results with the other methods.

## 2. Methods

### 2.1. Trait-space and tree-based approaches

#### 2.1.1 Convex Hull

Computing trait turnover between two communities using the convex hull methods simply consists in computing (i) the minimum convex polytopes (MCP) for each community, and (ii) the hypervolumes of the intersection and the union of these two MCPs (Figure 1a,e). It is then possible to compute a range of β diversity indices based on these four values. Here, following Mammola & Cardoso (2020), we used the Jaccard dissimilarity index *J* (Jaccard, 1908) and the Williams replacement index *W* (Williams, 1996), defined as:

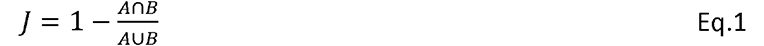

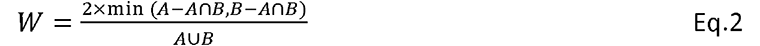

where A and B are the hypervolumes of the MCPs for two communities. In our analyses, we computed the MCPs and the indices using the hull.build() and hull.beta() functions from the BAT R package V.2.8.1 (Cardoso et al., 2015, 2022). The Williams replacement index evaluates the contribution of trait replacement to trait β diversity (Carvalho et al., 2012, 2013), and the difference between Jaccard and Williams indices quantifies how the trait richness difference between communities contributes to β diversity. Although there are other approaches and indices that can decompose β diversity into turnover replacement components, the relevance of these approaches is still debated (Baselga, 2010; Baselga & Leprieur, 2015; Cardoso et al., 2014; Carvalho et al., 2012, 2013). This debate is beyond the scope of this manuscript, and, to compare our methods, we only followed the decomposition used by Mammola & Cardoso’s (2020) (see sections on kernel density hypervolumes below), readily available from the BAT R package (Cardoso et al., 2015, 2022).

The main issue with the convex hull methods is that it is insensitive to the addition or removal of species within the MCP in the trait space (Figure 1). A corollary is that it is sensitive to outliers, as they will define the MCP.

#### 2.1.2 Tree-based method

The tree-based method consists in computing a dendrogram from the trait distance between all species in the species pool (i.e. the entire list of species over all included sites, not just those occurring in the pair of sites for each calculation of trait turnover; Figure 2a). Multiple clustering algorithms can be used to generate the dendrogram, but here we followed Loiseau et al. (2017) and used the unweighted pair group method with arithmetic mean (UPGMA) algorithm, using the hclust() function from the stats R package (R Core Team, 2022), as it has been shown to best conserve distances between species compared to the original distances in the trait space.

**Figure 2.**
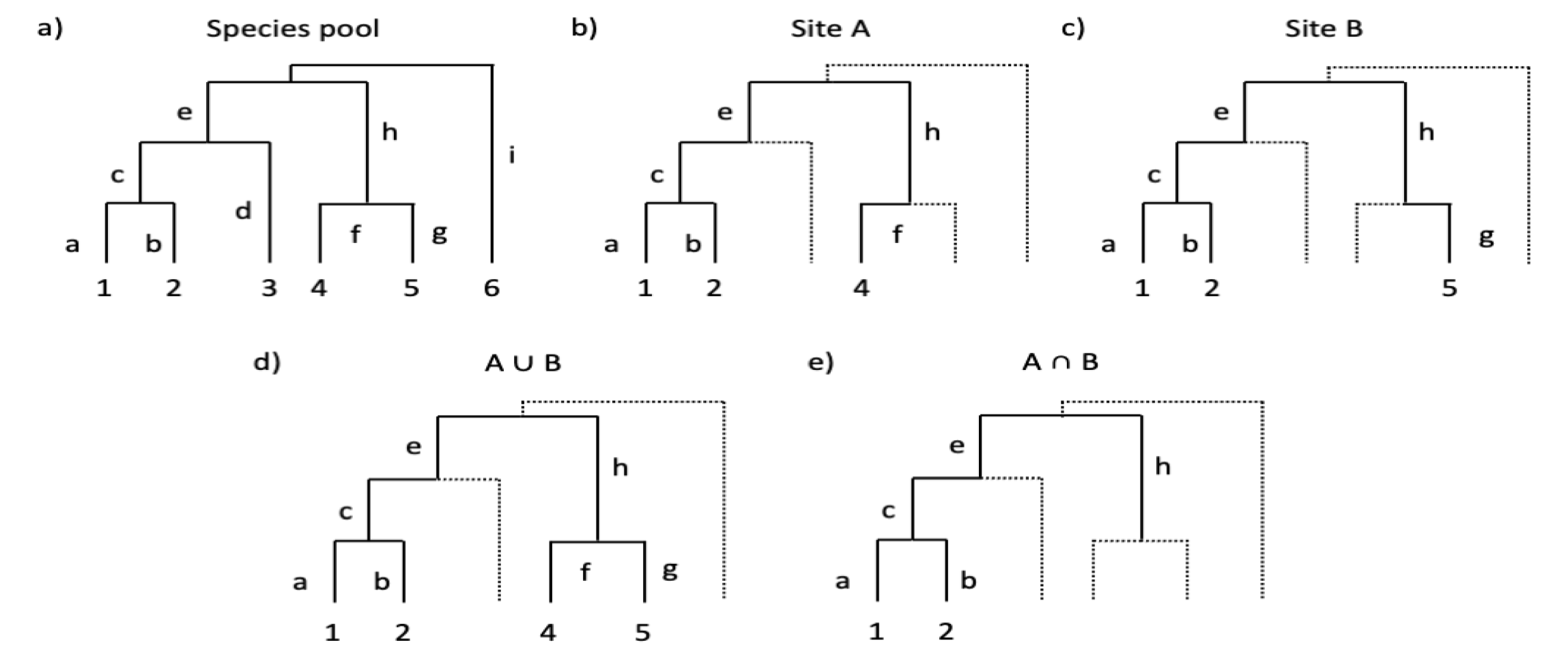
Components of the tree-based approach for the computation of trait turnover between two sites whose species are part of a larger species pool. a) Dendrogram for all species in the species pool. b) Dendrogram for species present in site A, after trimming the global tree from a). c) Dendrogram for species present in site B. d) Dendrogram for species present in either site A or site B. Since species 3 does not occur at any of the two sites, branch d is not included. e) Dendrogram for species present in both sites A and B. Although species 4 and 5 are present in only one of the two sites, branch h appears in both dendrograms, and is therefore conserved.

For each site, a sub-tree including only the species present is generated by trimming the overall tree (Figure 2b,c). It is then possible to compute the trees corresponding to the union and the intersection of the two sub-trees (Figure 2d,e). We can then adapt Eqs 1 and 2 to compute the Jaccard and Williams indices, by using the total length of remaining branches. Importantly, the sub-trees must be computed from the original tree generated from the entire species pool, not those computed from only the residing species. This conserves the internal branches in the union and the intersection of the two sub-trees, even if these internal branches do not lead to any present species (see branch h in Figure 2e). Therefore, for the example of Figure 2, Eqs 1 & 2 become:

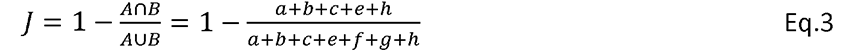

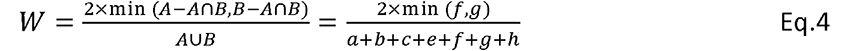

The tree-based method therefore offers the advantage over the convex hull method that all species will be accounted for when computing the β diversity indices. However, the clustering algorithms often generate branch lengths between species in the dendrogram that differ from the original distances in the trait space, which will necessarily influence the value of any β diversity index (Loiseau et al., 2017).

#### 2.1.3. Kernel density hypervolumes (KDH)

Mammola & Cardoso (2020) introduced the use of kernel density hypervolumes (KDH) for computing indices of species turnover. This approach uses KDEs to generate polytopes that are often non-convex (and can even be disjunct) and can be seen as a trait envelope around the species points in the trait space. The recommended method is based on a Gaussian estimator of the KDE (Mammola & Cardoso, 2020) and follows a series of four steps (see Blonder et al., 2018 for further details): (i) points are drawn randomly within a hypersphere around each species point in the trait space; (ii) these points are resampled to uniform density; (iii) a KDE is computed from these points; (iv) a threshold (typically 95%) is applied to truncate the KDE and define the polytope, from which hypervolumes can be computed. The indices are then computed as per Eqs 1 & 2. In our analyses, we used the kernel.beta() function from the BAT R package V.2.8.1 (Cardoso et al., 2015, 2022) to apply the KDH method.

This method, although more computationally intensive than the convex hull method, allows to account for the distribution of species points in the trait space to define the polytopes and therefore the hypervolumes used in the computation of the turnover indices (Figure 1b-d,f-h). As a result, the KDH method is less sensitive to outliers.

The KDH method nonetheless has some caveats. First, the choice of the threshold used to construct the polytope will necessarily influence the components of the β diversity indices, and therefore the final values. Second, by resampling random points to uniform density, some information about the distribution of species points in the trait space is lost. Finally, in the current implementation of the method in the BAT R package V.2.8.1 (Cardoso et al., 2015, 2022), the radius of the hyperspheres within which random points are drawn around the species points and the bandwidth used during the computation of the KDE (the bandwidth is a parameter that determines how smooth the KDE will be) are determined based on the species point distribution of each community separately using the estimate_bandwidth() function from the hypervolume R package. As a result, the more similar species are to each other, the closer random points will be to each other and the KDE will show a steeper gradient (Figures A1-A24, see especially Figures A1, A9 and A17). In other words, using different bandwidths and resampling random points to uniform density gives different weights to a species depending on how different its traits are from those of other species in the community. For a β diversity index to be unbiased we argue that all species should have the same weight when relative abundance and intraspecific trait variation are not concerned.

### 2.2 A kernel integral method (KIM)

Here we propose a novel computational method to compute trait turnover in the trait space, to solve the issues associated with the KDH method. Our method computes β diversity indices from the kernels themselves, therefore removing the influence of the threshold used to generate the polytope, and uses different kernels than those used in the KDH method. The KIM method consists of using only steps (i) and (iii) from the KDH method: (i) points (typically 1000, but the number can be adjusted to account for species abundance, for example) are drawn randomly within a hypersphere around each species point in the trait space (the diameter of the hypersphere can be the same for all species, or reflect intraspecific trait variability); (iii) a KDE is computed from these points, and rescaled between [0,1]. From the KDE, we then propose the following equations to compute the Jaccard dissimilarity index and the Williams replacement index:

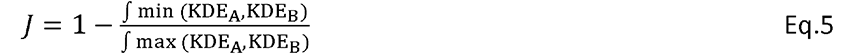

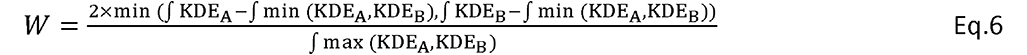

where KDE_A_ and KDE_B_ are the KDEs for communities A and B, and ∫ KDE_A_ is the integral of the KDE for community A over all dimensions of the trait space. This is similar in essence to the index of niche overlap proposed by Mouillot et al. (2005). In practice, since KDEs are computed as multi-dimensional matrices, an integral is simply computed as the sum of all elements of the matrix. The minimum and the maximum of two KDEs are analogue to the intersection and the union of the polytope in the KDH method (Figure 3).

**Figure 3.**
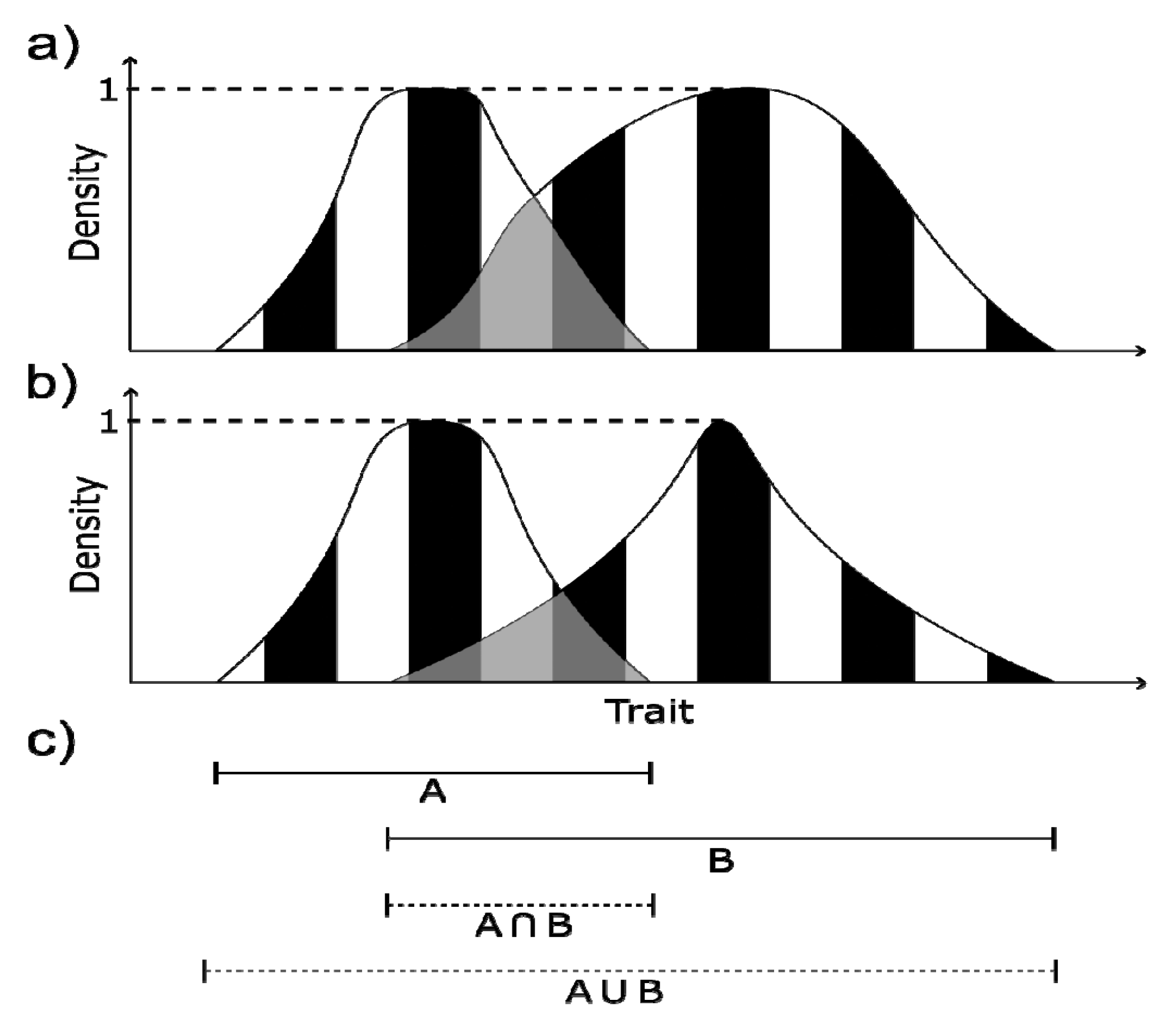
Details of the computation of trait turnover for two pairs of communities (one pair in (a) and one in (b)) following different approaches. In each graph (a,b), the curves fictional KDEs (Kernel density estimators) for the two communities, in one dimension (for simplification, we assume their densities are 0 beyond intersecting the horizontal axis). Using the KIM (Kernel integration method) formula, The Jaccard index is computed as one minus the area in grey divided by the striped area (Eq. 5), and the value is different for the two pairs of communities in (a) and (b) (as is the value of Willams replacement index, Eq. 6). (c) The horizontal segments represent one-dimensional polytopes (defined using the values where the KDEs intersect the horizontal axis for simplification), used to compute the Jaccard or Williams indices using the KDH (kernel density hypervolume) method (Eqs 1, 2). Contrary to KIM, the KDH method only generates a single value for each index for the two pairs of communities.

This kernel integral method enables us to overcome the limitations of the KDH method mentioned above. First, there is no need to define a threshold: if the KDE is estimated over a large enough area or volume, the local kernel density will approach zero and the integral will therefore converge. Second, the radius within which the random points are drawn is the same for all communities (but a suitable value must be chosen, which can be adjusted to account for intraspecific trait variability). Finally, because there is no resampling to uniform density, the distribution of species points in the trait space will be reflected more accurately in the KDE.

### 2.3. Test of the different methods on theoretical data and expectations

We have described above the theoretical advantages and caveats of each of the different methods. We implemented seven different methods in total to explore how the differences between the characteristics of the methods can influence the results (Table 1):

- A convex hull method (hereafter COVHULL).
- A tree-based method (hereafter TREE).
- The original kernel density hypervolume method with community-specific bandwidths and uniform resampling (hereafter KDH V1).
- A modified kernel density hypervolume method computed with the same bandwidth for each pair of communities and uniform resampling (hereafter KDH V2), to explore the influence of the bandwidth on the outcome.
- A kernel integral method using kernel densities estimated with community-specific bandwidths and uniform resampling (hereafter KIM V1), to explore the influence of the kernel-based vs the polytope-based formulas (Figure 3).
- A kernel integral method using kernel densities estimated with the same bandwidth for both communities within a pair and uniform resampling (hereafter KIM V2), to further explore the influence of the kernel-based vs the polytope-based formulas.
- A kernel integral method estimated with the same bandwidth for both communities within a pair and without uniform resampling (hereafter KIM V3).

**Table 1.** Summary of the methods used to compute trait turnover.

| Method | Summary | Advantages | Caveats | References |
| --- | --- | --- | --- | --- |
| Convex hull method (COVHULL) | A minimum convex polytope (MCP) encompassing all species points in the trait space is computed for each community. The hypervolumes are used in the computation of the $\beta$ diversity indices. | The distance between the original species points in the trait space is representative of how functionally different the species are. | Only the species with extreme trait values are used in the computation of the $\beta$ diversity indices. Sensitive to outliers. | (Cornwell et al., 2006; Villéger et al., 2008, 2013) |
| Tree-based method (TREE) | A dendrogram is computed for the species pool based on the distance between all species based on their traits. The resulting tree is trimmed for each community, and the lengths of branches are used in the computation of the $\beta$ diversity indices. | All species contribute to the computation of the $\beta$ diversity indices. Not as sensitive to outliers as the convex hull method. | The distance between the species computed from the dendrograms is somewhat different from the original trait difference between species for the traits considered in the analyses. | (Cardoso et al., 2014) |
| Kernel density hypervolume (KDH) V1 | A polytope is computed for each community based on a kernel density estimator (KDE). The hypervolumes are used in the computation of the $\beta$ diversity indices. | The distance between the original species points in the trait space is representative of how functionally different the species are. The polytope accounts for how species are distributed in the trait space. | Sensitive to the threshold used to compute the MCP. The random points are resampled to uniform distribution, which diminishes the effect of the species point distributions in the trait space. A different bandwidth is used for each community to generate the random points and compute the KDE. As a result, the KDE is influenced by the distance between the species in the trait space. | (Mammola & Cardoso, 2020) |
| Kernel density hypervolume (KDH) V2 | As KDH V1, but using the same bandwidth for each pair of communities. | As KDH V1, plus independence from how different species are from each other in each community. | As KDH V1, except for the issue related to using different bandwidths. | This article |
| Kernel Integral Method (KIM) V1 | The $\beta$ diversity indices are directly computed from KDEs of species points in the trait space. The KDEs are based on the same procedure as KDH V1. | As KDH V1, but with the advantage of being insensitive to the threshold used to compute the MCP. | | This article |
| Kernel Integral Method (KIM) V2 | As KIM V1, but using the same bandwidth for each pair of communities. | As KDH V2, but with the advantage of being insensitive to the threshold used to compute the MCP. |  | This article |
| Kernel Integral Method (KIM) V3 | The $\beta$ diversity indices are directly computed from kernel density estimators of species points in the trait space, but the KDEs are estimated directly from points randomly drawn around species points in the trait space, without resampling to uniform density. | As KDH V1 & V2, but reflecting more accurately the distribution of species points in the trait space. The radius of the hyperspheres still needs to be determined. | | This article |

For each method, we computed Jaccard dissimilarity and Williams replacement, as defined in Eqs 1-6. We then examined how these seven methods behaved in a set of theoretical contexts for which we can make predictions of how an index of turnover should behave to capture trait differences between communities.

In total, we simulated 72 different pairs of communities (Figures 4 and A1) and computed our 14 indices (the Jaccard and Williams indices for each of the seven methods) for each pair. For simplicity and computational efficiency, we used a trait space defined by two theoretical traits. Each community was first delimited by a MCP represented by four species arranged as a square. The MCPs were either of different sizes (square side of lengths 4 and 2; Figure 4) or of the same size (square side of length 4; Figure A1). They were either nested within each other, partially overlapping, or disjunct. For each of these configurations, we generated a community by randomly drawing species points within the MCPs according to three patterns: (i) the species points were located in a small area (square sides of length 1) in opposite corners of the MCPs (hereafter called the “different” point distribution); (ii) the species points were randomly drawn within the MCPs (hereafter called the “random” point distribution); (iii) the species points were located in a small area (squares of length 1) in the closest corners of the MCPs (hereafter called the “similar” point distribution). We tested these 18 configurations for 10, 40, 70 and 100 species points, and performed analyses 50 times for each of the resulting 72 configurations (2 MCP size setups x 3 relative positions x 3 random point distributions x 4 sets of point numbers x 50 repeats = 72 configurations x 50 repeats).

**Figure 4.**
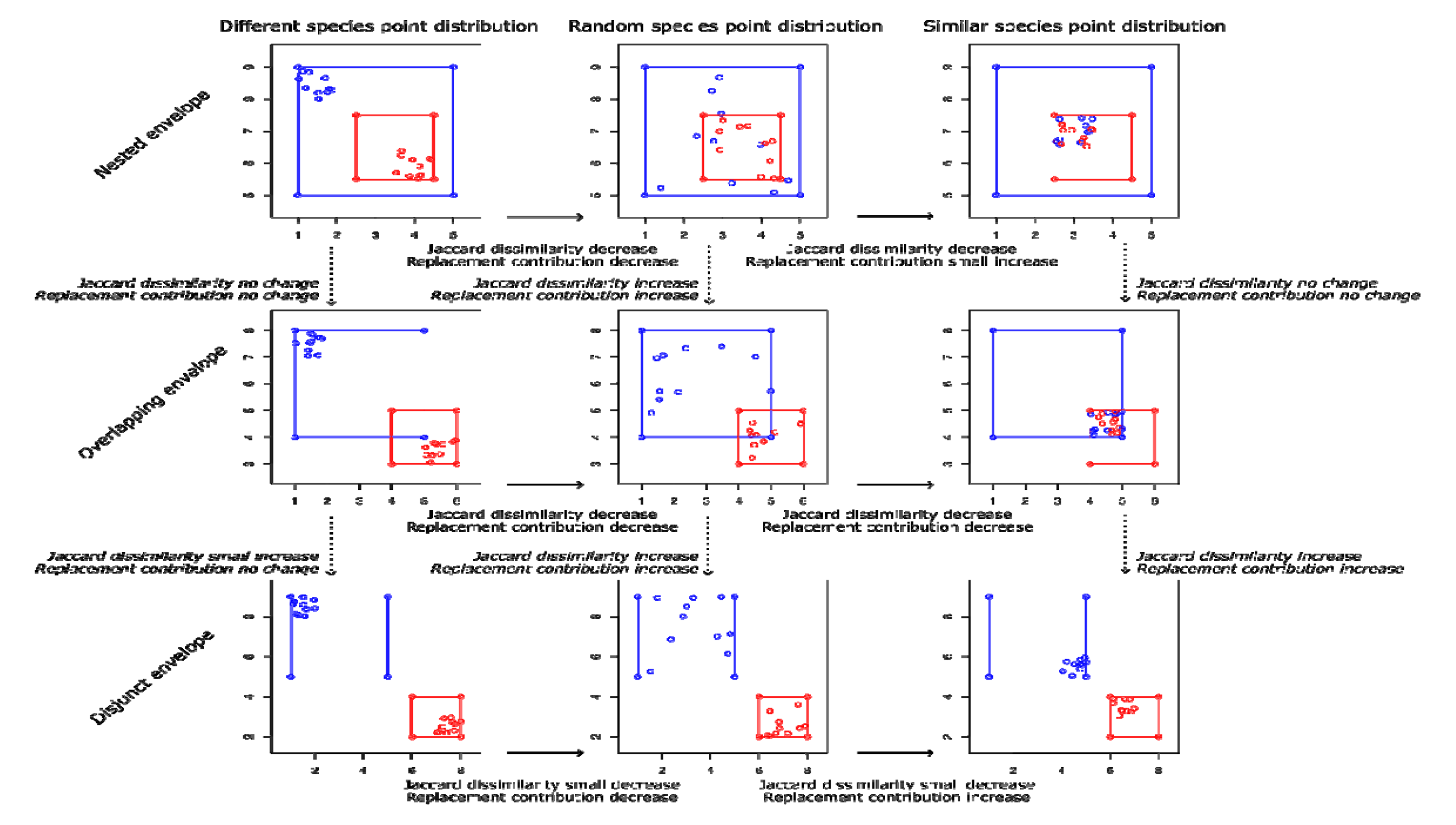
One of the 50 instances of the nine theoretical configurations of pairs of communities for different sizes of the MCPs (see Figure A1 for same size MCPs), using only 10 randomly drawn species points for clarity (point distributions for 40, 70 and 100 species were also generated). We varied how the MCPs overlapped (“nested”, “overlapping” or “disjunct”), and how the species points are distributed within the MCPs (“different” – distributed in opposite corners of the MCPs –, “random” – randomly distributed within the MCPs – or “similar” – distributed either within the same small area, or in the closest corners of the MCPs).

There is one obvious difference between these theoretical communities and communities that would be analysed for real-world applications: real-world communities belonging to the same ecological system such as those described in the next section will usually share species, resulting in many species points overlapping in the trait space. Here we used independent random species distributions in the trait space for the two communities in order to have more flexibility in these species distributions, to explore in detail how each of the seven methods would behave across a wide variety of extreme configurations, and better test disentangle the implications of their computational specificities.

This flexibility allows us to describe how a β diversity index should behave based on what it is supposed to capture from these theoretical configurations. These expectations are depicted in Figures 5, A2, A3, and their justification provided in Table A1. In summary, Jaccard dissimilarity should increase as most species points in the two communities move away from each other. The replacement component should decrease if the difference in area covered by the two sets of species points increases. These patterns should be especially clear for large numbers of species, i.e. for high densities of species points. For few species and low species point density (e.g. when only 10 species points were randomly drawn in the trait space), we expect these patterns to be weak, as the stochastic element of the species point distributions may obscure the results.

**Figure 5.**
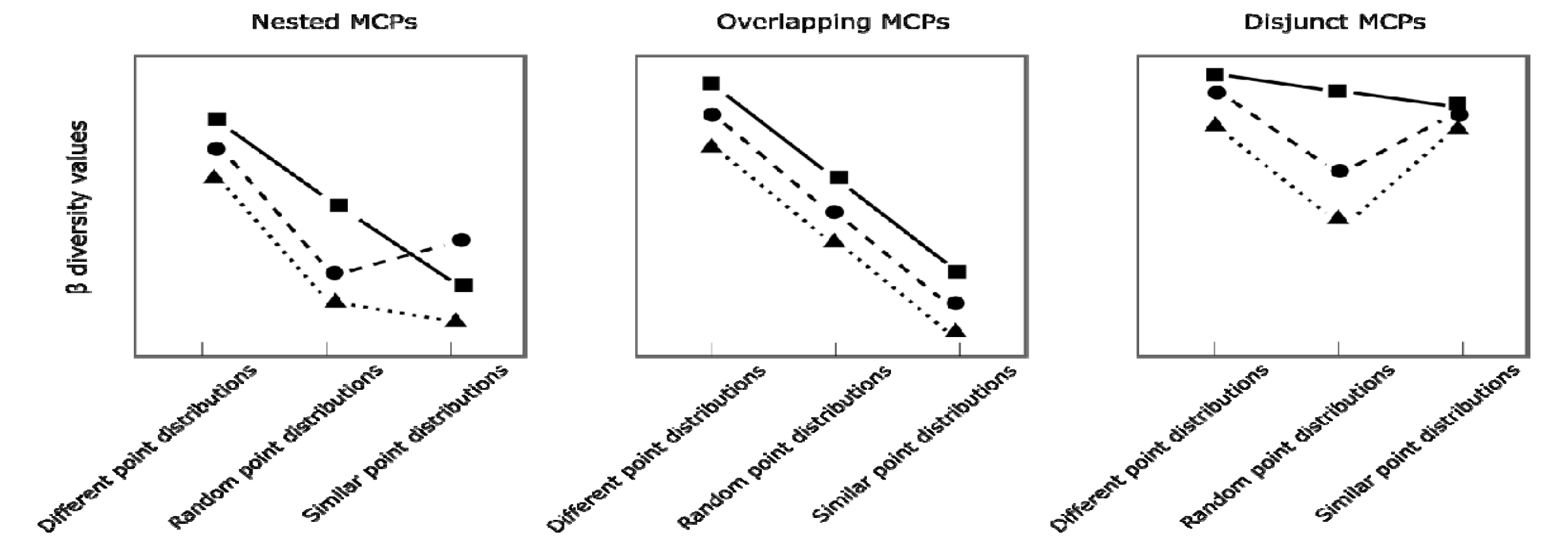
Qualitative differences in β values for trait turnover predicted under different simulated configurations of species points in the trait space when the MCPs of the two communities have different sizes. Jaccard dissimilarity (squares, solid lines), Williams replacement (triangles, dotted lines), and contribution of replacement to overall turnover, computed as Williams replacement divided by Jaccard dissimilarity (circles, dashed lines). Overall, β values, are expected to decrease as the overlap between the trait profiles of the two simulated communities increases. Jaccard dissimilarity accounts for differences in the spread of the trait profiles in the trait space (i.e. trait richness), whereas Williams replacement is independent from differences in trait richness. For detailed explanations about the changes in β values, see Table A1 in SI.

### 2.4. Established non-native plants in French Polynesia

To examine how the different methods may lead to different conclusions when analysing real data, we examined the trait diversity of plant species introduced to the Pacific islands of French Polynesia, comparing trait turnover across islands and archipelagos using each method. We extracted data from PacIFlora (Wohlwend et al., 2021). For French Polynesia, PacIFlora contains data on naturalised non-native plant species across the 80 Pacific islands over five archipelagos: The Society Islands, the Gambier Islands, the Tuamotu Islands, the Tubuai Islands, and the Marquesas. However, careful examination of the database revealed that some species were not naturalised but cultivated or endemic. We therefore only used the 417 naturalised species in PaciFlora appearing in the Appendix of Fourdrigniez & Meyer (2008).

For these 417 species, data on species woodiness (woody vs. herbaceous species), seed mass, plant height and specific leaf area (SLA) were extracted from multiple trait databases, including TRY (Kattge et al., 2011, 2020), LEDA (Kleyer et al., 2008), PLANTATT (Hill et al., 2004), Austraits (Falster et al., 2021), BIEN (Maitner, 2022), EcoFlora (Fitter & Peat, 1994), and BROT (Tavşanoğlu & Pausas, 2018). Seed mass, plant height and SLA have been used to characterise different plant life strategies (Díaz et al., 2016; Westoby, 1998). When different databases contained different values, we used the mean for seed mass, plant height and SLA, and the most frequent category for woodiness. Data on plant woodiness was available for all 417 species. Trait data for seed mass and plant height were only available for 250 out of the 417 species. Data for seed mass, plant height and SLA were only available for 124 out of 417 species. We therefore performed three sets of analyses: (i) a set for the 250 species with data on seed mass and plant height, (ii) a set for the 124 species with data on the three traits, and (iii) a set for the same 124 species, using data on seed mass and plant height only, to assess the robustness of the results to data availability and trait selection. In the following we present and discuss mainly results for seed mass and plant height for the 250 species (see Figure D1 for the distribution of plant species in this two-dimensional trait space), as it represents more than half of the species and should be less biased despite using only two traits.

Prior to analysis, seed mass, plant height and SLA were log-transformed and rescaled between [0,1], so that the traits would be more uniformly distributed in the trait space. We then computed the Jaccard dissimilarity and the Williams replacement indices for all species together, and for woody and herbaceous species separately, to have a more comprehensive assessment of potential differences between methods, as woody and herbaceous plants tend to characterise different parts of the global spectrum of plant forms and functions (Díaz et al., 2016). We also computed these indices for all French Polynesian islands, and for each archipelago separately.

Finally, the behaviours of the different indices were analysed using randomisation tests. We randomised the presence-absence matrices for all islands and for each archipelago by keeping species occupancy and island richness constant (i.e. the sim9 algorithm from (Gotelli, 2000)), and compared the Jaccard dissimilarity and Williams replacement indices generated by the 7 methods for the original matrices to the indices computed over 10 randomised matrices for each original matrix (the number of randomisations was constrained by computation time).

The purpose of these analyses on real data was only to examine how results may differ between methods for more complex data than used in the theoretical analyses, to assess each method’s range of sensitivity. As each archipelago contains multiple different islands whose combinations will fall across a large spectrum of trait profile configurations, it was not possible to define a priori expectations and the purpose is therefore not to determine which methods are in line or not with a priori expectations, contrary to the theoretical analyses.

## 3. Results

### 3.1. Theoretical data

Overall, KDH and KIM tended to converge towards similar values and behaviours as the number of species points increased (Figures 6-8, C1-C12), corresponding to theoretical expectations (Figures 5, A2, A3). In contrast, the convex hull and the tree-based methods generated indices of turnover different from the other methods and from the theoretical expectations. The main differences between observed and expected values for all methods, except the tree-based method, were for the contribution of replacement to overall turnover (computed as the ratio of the Williams replacement index to the Jaccard dissimilarity index) for MCPs of the same size in the nested / random and the nested / similar configurations, which was lower than the expected value of 1 (Figures A3, C6). This is likely due to the fact these are the configurations for which the values of the Jaccard index are small, and small changes in Williams replacement index due to stochasticity in the distribution of species points in the trait space will be disproportionally large.

**Figure 6.**
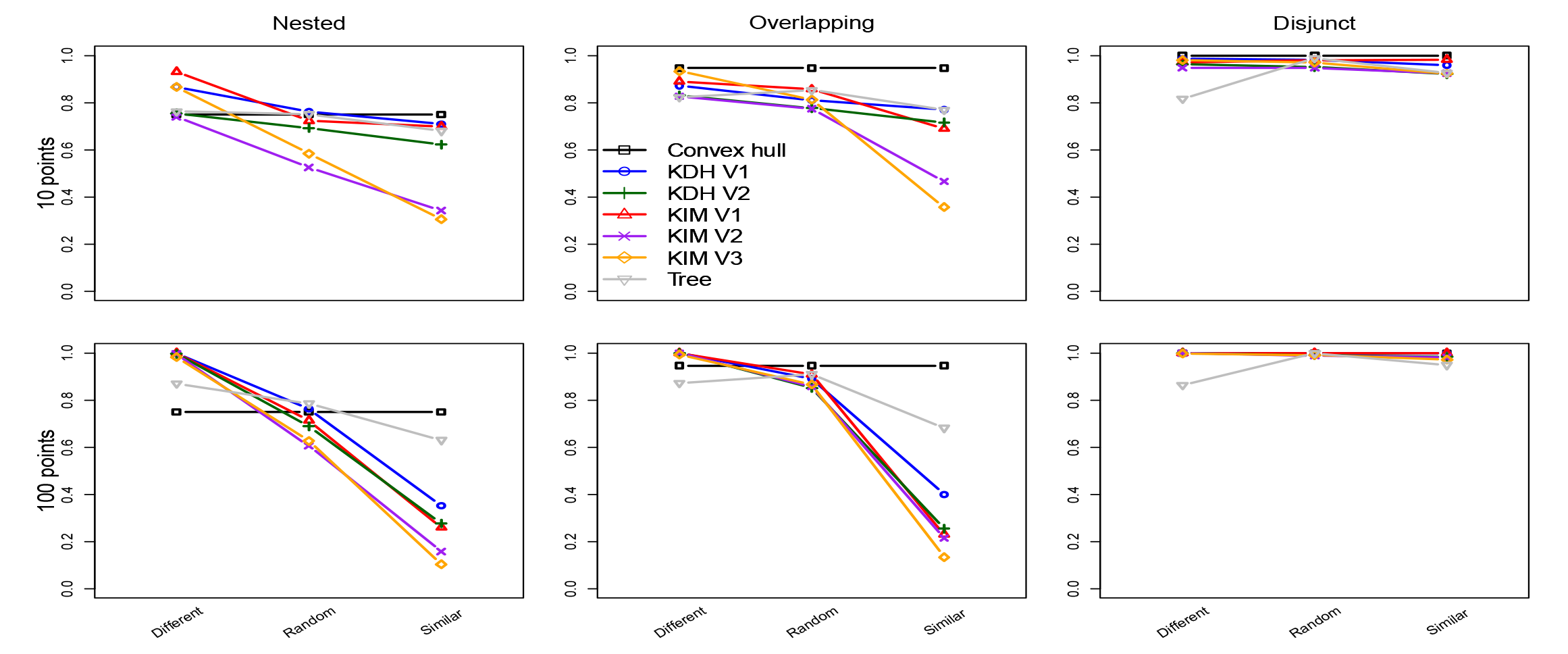
Differences in Jaccard dissimilarity between the seven methods summarised in Table 1, for MCPs of different sizes, for 10 and 100 species points (see Appendix C for the full set of results).

For all indices of turnover, the three KIM methods generated values above 0.5 and above other methods when the point distributions were different from each other (i.e. the “Different” point distributions under all MCP configurations, and for all three point distributions under the “Disjunct” MCP configuration). KIM V3 generated values below 0.5 and below other methods when the point distributions were similar from each other (i.e. the “Similar” point distributions under all MCP configurations), and intermediate values otherwise, in-between the values generated by the other methods. These results suggest KIM V3 can better distinguish between different species point distributions in the trait space (Figures 6-8, C1-C12). The KIM V3 method also tended to be less sensitive to the number of species points than the other KDH and KIM methods, with values and behaviours being similar from 10 to 100 species points.

When communities had MCPs of the same size, the KIM V1 and V2 methods generated similar results to the KDH V1 and V2 methods, respectively, for all β diversity indices (Figures C4-C6, C10-C12). However, when the MCPs had different sizes, the KIM methods tended to generate values more similar to each other than to the KDH methods (Figures 6-8, C1-C3, C7-C8).

Adjusting the bandwidth to be common between communities in each pair in the computation of the kernels for the KDH and KIM methods (i.e. switching from V1 to V2) resulted in lower dissimilarity values, both for the Jaccard dissimilarity index and the Williams replacement index, for all configurations. This is because the radius of the hyperspheres and therefore the steepness of the kernels were the same for both communities in the V2 methods, increasing similarity. The effect of removing the resampling of random points to uniform density (i.e. from KIM V2 to KIM V3) had an often larger and more variable effect than adjusting the bandwidth, as the values generated by KIM V3 could be either larger, smaller or in between those of the KIM V1 and V2 methods.

### 3.2. Established non-native plants in French Polynesia

Raw values of Jaccard dissimilarity, of Williams replacement and of the contribution of replacement to turnover differed greatly between the different methods. Maximum differences in values between methods were around 0.6 for Jaccard dissimilarity, 0.2 for Williams replacement, and 0.8 for the contribution of replacement to turnover (Figures 9, C2, C3). The KIM V3 and the KDH V2 methods generated the lowest Jaccard dissimilarity, and KIM V1 and TREE the highest. In contrast, KIM V3 consistently generated much higher values for the contribution of replacement to turnover than other methods, as expected from the fact that it better accounts for differences in species point distributions in the trait space. Results showed similar trends for all the combinations of traits and number of species used in the analyses (Figures D2, D3).

Importantly, compared to the other methods, the KIM methods sometimes generated a different ranking between archipelagos for Jaccard dissimilarity. This is especially true for woody species, for which KIM V3 suggests that trait turnover was higher for the Gambier than for any other archipelagos, for all combinations of traits, and for both Jaccard and Williams replacement indices (Figures 9, D2, D3). By contrast, the other methods generated results that were more variable depending on the combination of traits and species used.

Randomisation of presence-absence matrices show that the KDH V2, KIM V2, KIM V3 and Tree methods tended to generate more consistent values for Jaccard dissimilarity compared to the convex hull, KDH V1 and KIM V1 methods (Figure D4). For Williams replacement, values were also more consistent across randomised matrices for KIM V3 than for the other methods, especially for herbaceous species.

## 4. Discussion

Here we compared existing and novel methods to compute trait turnover for simulated and empirical data, to illustrate how differences in the computational aspects of these methods reflect different aspects of trait diversity and can affect inferences made from trait diversity comparisons.

### 4.1. Theoretical aspects of trait turnover computation

Comparing the seven methods using simulated data, for which we had complete control of the community trait profiles in the trait space, revealed the important effects of the computational specifics of each method on the value of trait β diversity. In particular, we assessed the effect of conserving trait distance between species by comparing the tree-based method, which distorts the trait distance between species in the dendrogram (Maire et al., 2015), to the trait space-based methods. The tree-based method consistently generated results most different from theoretical expectations (Figures 5-8). It tended to either underestimate or overestimate dissimilarity in the trait profiles of communities in the MCPs, and tended to generate high values for the Williams replacement index.

Interestingly, the convex-hull methods produced results similar to the tree-based method for Jaccard dissimilarity, also generating results that differed from theoretical expectations. This is because CONVHULL only uses a subset of the species in the trait space. Although using a trait space approach better conserves trait distance between species than a tree-based approach when all species are included, this property is broken when species are ignored in the computation of trait turnover. Therefore, although the CONVHULL and tree-based methods have been contrasted in the literature and have been shown to generate different results (e.g. Loiseau et al., 2017), we show that neither of these two methods accurately reflects the trait profile of a community.

In contrast, the other five trait space methods compared in this article (KDH V1-V2 and KIM V1-V3) generated results more in line with theoretical expectations. These methods therefore offer a more consistent representation of the community trait profile in the trait space, i.e. they better capture the contribution of all species to the assessment of turnover. The computational aspects of these approaches to estimating trait turnover have nonetheless important effects on the generated β values, with potential consequences for inferences about made trait turnover in an assemblage or community.

Specifically, we explored three computational aspects of these methods: (i) the use of polytopes vs kernel integrals (KDH V1 vs KIM V1; Eqs 1 & 2 vs Eqs 5 & 6); (ii) the use of the same or different bandwidths for each community in a pair when computing the KDE (KDH V1 vs V2 and KIM V1 vs V2); and (iii) the use of point resampling when computing the KDE (KIM V2 vs V3). All three aspects proved to have important effects on the β diversity values calculated. Using kernel integrals, the same bandwidth and not resampling (i.e. using KIM V3) generated results most in line with theoretical expectations.

The respective effects of these three computational aspects on trait turnover (β values) depend on the index used (Jaccard dissimilarity or Williams replacement) and on the configuration of the community trait profiles. For example, Jaccard dissimilarity is sensitive to the difference in bandwidth between communities (Figure 6). This is because using different bandwidths changes the shape of the KDEs (akin to making the distributions larger or narrower in Figure 3) and generates polytopes with different areas (akin to changing the lengths of A and B in Figure 3). Consequently, Jaccard dissimilarity values reflect this artificial difference in trait richness, but not by Williams replacement. In contrast, for Williams replacement, the use of polytopes or kernel integrals proved to be the most important factor (Figure 7). This is because kernel integrals better reflect small variations in the shape of the KDE (akin to changing the shape of the distributions and the overlapping area in Figure 3) and thus better capture trait replacement. Similarly, resampling also affected Williams replacement for communities with an “overlapping” configuration (Figure 7), especially for species-poor communities. This is because, when compared to the more uniformly distributed trait profile of species-rich communities, each species has a greater weight on the shape of the trait profile in species-poor communities, and the idiosyncrasy in the position of different species can drastically change trait profiles if resampling is not applied.

**Figure 7.**
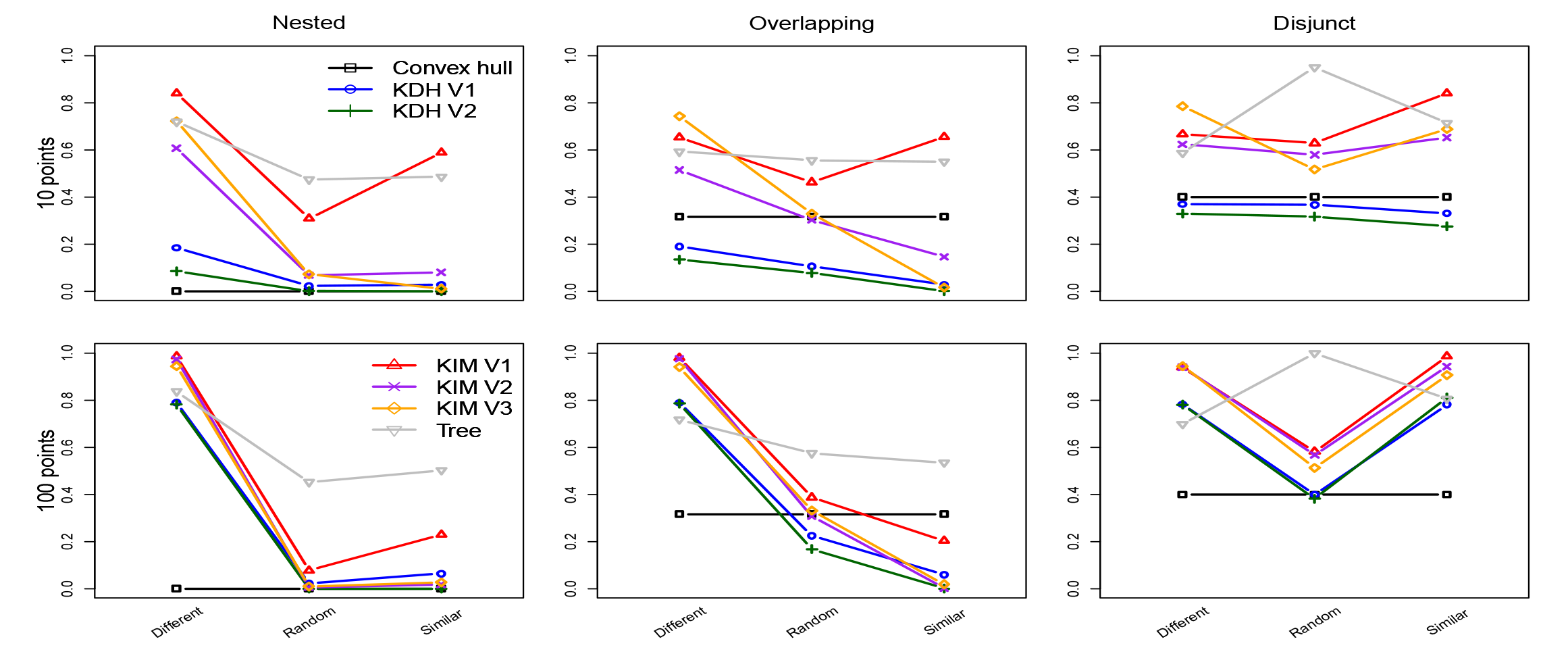
Changes in Williams replacement index for the seven methods summarised in Table 1, for MCPs of different sizes, for 10 and 100 species points (see Appendix C for the full set of results).

**Figure 8.**
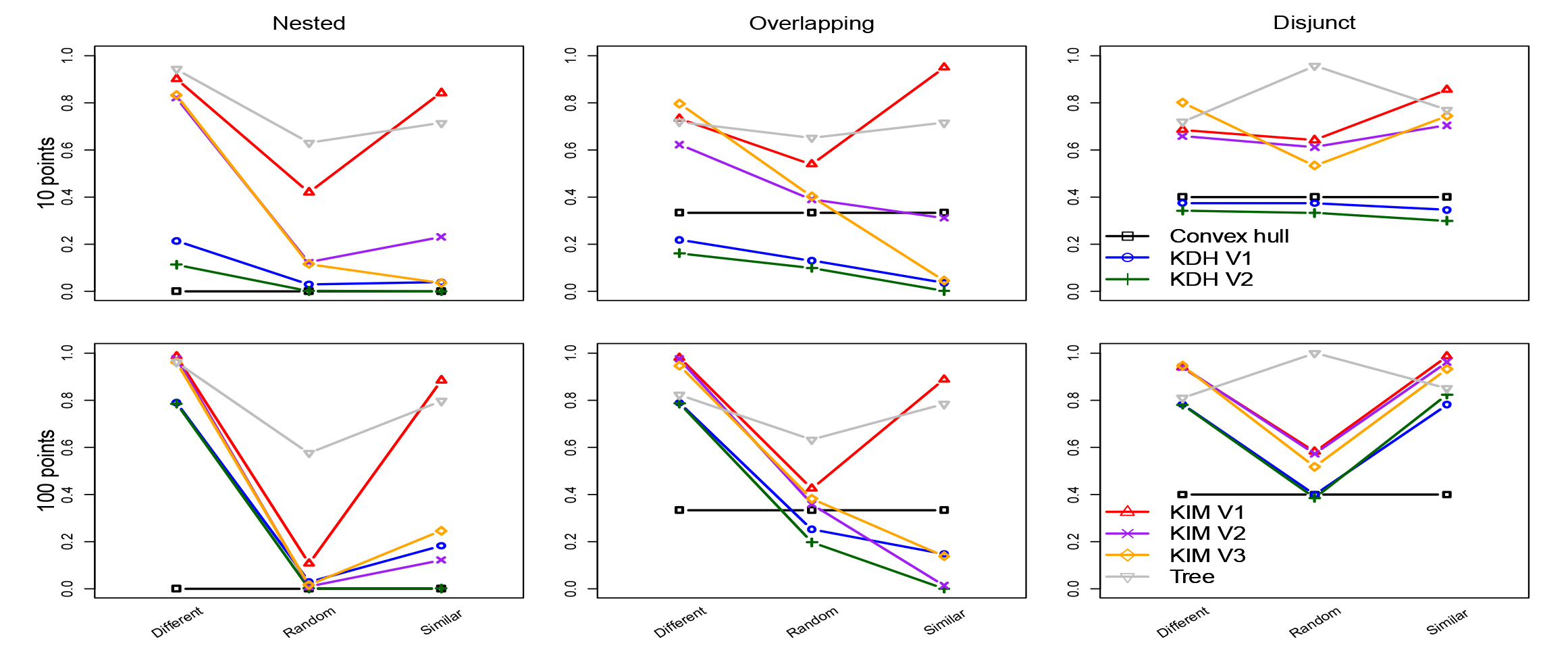
Changes in the contribution of replacement to overall turnover, computed as Williams replacement divided by Jaccard dissimilarity, for the seven methods summarised in Table 1, for MCPs of different sizes (see Appendix C for the full set of results).

**Figure 9.**
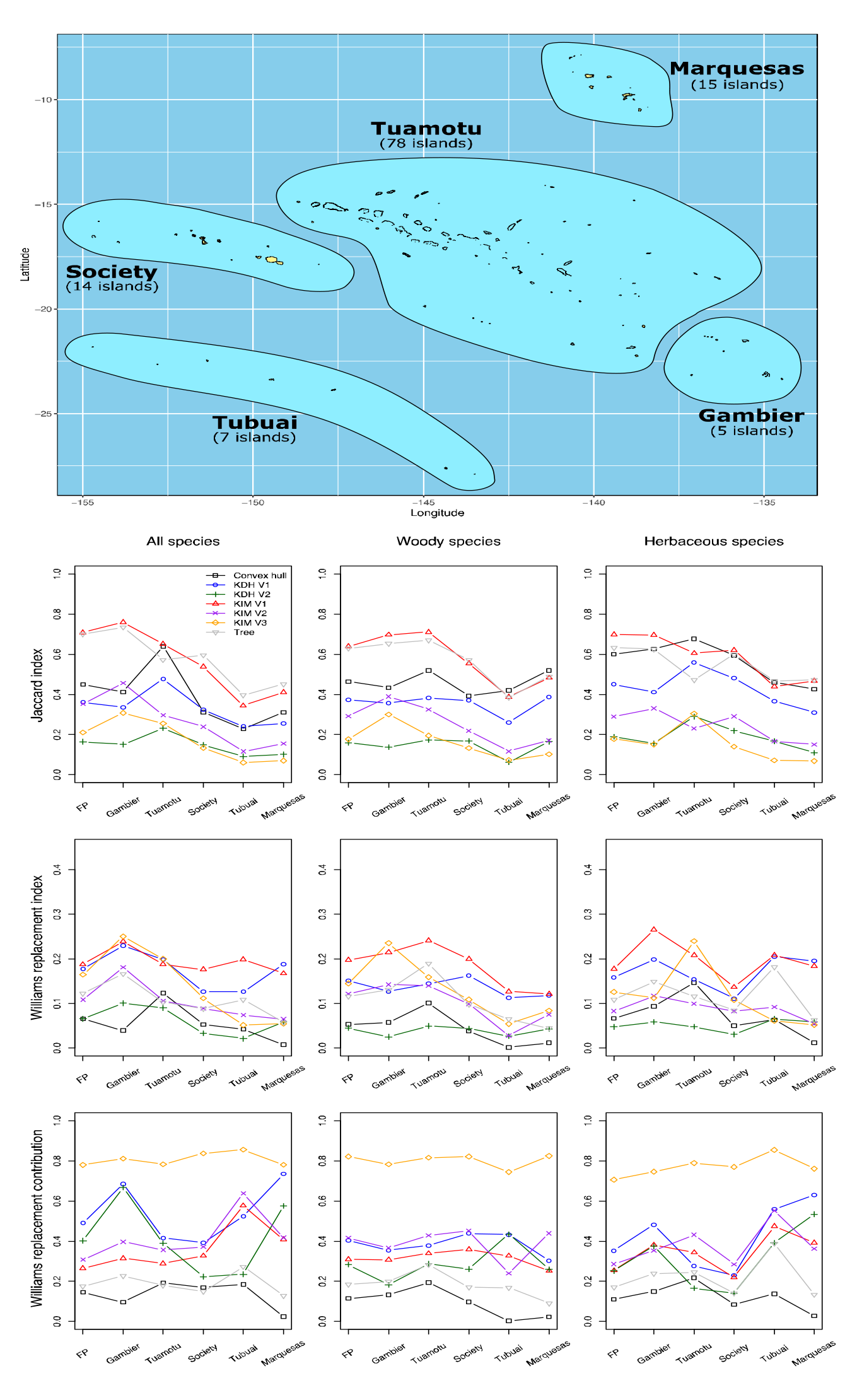
Application of seven trait turnover methods to empirical data on French Polynesian plant species. Jaccard dissimilarity index, Williams replacement index, and contribution of replacement to overall turnover, computed as Williams replacement divided by Jaccard dissimilarity, for the seven methods summarised in Table 1, for French Polynesia (FP) and its archipelagos, for all species, woody species and herbaceous species, using seed mass and plant height, for 250 out of 417 species (see Figures C2 and C3 for using seed mass, plant height and SLA on 124 species, and seed mass and plant height on 124 species). Note that the order of the islands is arbitrary, and lines between symbols are used as a visual aid and not to depict continuous change.

Both trait envelope, as captured by a convex hull, and kernel-based community trait profiles have complementary uses, and the choice of an analytical approach will depend on the research or management question. On the one hand, species with extreme trait values defining a trait envelope for a given community can help capture the whole range of trait values of species that may potentially join the community. The trait envelope may therefore be an important piece of information to assess the risk of potential invaders to a region (e.g. the join the locals vs. the try harder hypothesis; Tecco et al., 2010), or an indicator of the loss of trait extremes. The CONVHULL method is appropriate for such applications. On the other hand, capturing the trait distribution of all species in a community in the trait space provides a more comprehensive description of trait diversity and is necessary for identifying community assembly processes (Falster et al., 2017). The distributional profile of species in the trait space can also highlight gaps within the trait envelope, where introduced species with corresponding traits could have a higher chance to establish (i.e. the “empty niche hypothesis”; MacArthur, 1970; Molofsky et al., 2022). This community trait profile thus reflects ecosystem resilience and trait redundancy (Hui et al., 2021; Mouillot et al., 2021). Our results suggest that the KIM V3 method is most informative and least biased for addressing trait diversity questions.

### 4.2. Empirical test of methods using plant data from French Polynesia

The empirical testing of these methods provided further insight on the behaviour of the different methods for a mixture of configurations of species points in the trait space (Figure D1), and on how using different indices can lead to different conclusions. The KIM V3 method consistently generated much higher values for the contribution of replacement to turnover than other methods, even when randomising site-by-species matrices (Figures 9, D2-D4), suggesting that the higher Jaccard dissimilarity values generated by the other methods may reflect an overestimation of the contribution of trait richness difference (the complement of replacement) to turnover. Importantly, depending on the method used, one could either conclude that most islands of an archipelago are very different from each other in terms of trait diversity (e.g. Jaccard dissimilarity values > 0.5 for KIM V1 for the Gambier, Tuamotu and Society archipelagos), or very similar (Jaccard dissimilarity values mostly < 0.2 for KIM V3). These different conclusions, in addition to the different rankings generated by the different methods could be crucial for conservation decisions. For example, assuming management actions are influenced by species traits, low trait dissimilarity between islands, as indicated by KIM V3, would suggest that a similar management approach is appropriate across most islands, simplifying management and potentially improving management efficiency. In addition, the high contribution of replacement to turnover suggests that existing differences in community trait profiles are unlikely to be the result of differences in colonisation pressure, and may point towards either idiosyncratic or niche-driven factors.

## 5. Final recommendations / Conclusion

The kernel integral method presented here computes trait β diversity by directly integrating KDEs. Out of the three different indices this method can generate, KIM V3 implements the same bandwidth for the paired communities without resampling random points to uniform distribution. KIM V3 generates values that better reflect the distribution of species in the trait space (i.e. the community trait profiles) than methods based on convex hulls or dendrograms, and also better than other methods based on KDEs. The approach is also flexible, information rich and readily adapted to account for relative abundance between species and intraspecific trait variation, by using different numbers of random points and radii to generate the KDEs. Together with the convex hull method to inform on the trait envelope, and tree-based approaches for quantifying phylogenetic diversity, and the kernel integral method using the same bandwidth and non-uniform point distribution provide a complementary set of metrics for understanding patterns of trait diversity and turnover.

## Acknowledgements

We thank Jean-Yves Meyer for his time and expertise to refine the database on naturalised alien plants species in French Polynesian islands. CH is supported by the NRF (89967), the NERC (NE/V007548/1) and the Horizon Europe (101059592).

## References

Ackerly, D. D., & Cornwell, W. K. (2007). A trait-based approach to community assembly: Partitioning of species trait values into within- and among-community components. Ecology Letters, 10(2), 135–145. https://doi.org/10.1111/j.1461-0248.2006.01006.x

Anderson, M. J., Crist, T. O., Chase, J. M., Vellend, M., Inouye, B. D., Freestone, A. L., Sanders, N. J., Cornell, H. V., Comita, L. S., & Davies, K. F. (2011). Navigating the multiple meanings of β diversity: A roadmap for the practicing ecologist. Ecology Letters, 14(1), 19–28.

Baselga, A. (2010). Partitioning the turnover and nestedness components of beta diversity. Global Ecology and Biogeography, 19(1), 134–143.

Baselga, A., & Leprieur, F. (2015). Comparing methods to separate components of beta diversity. Methods in Ecology and Evolution, 6(9), 1069–1079. https://doi.org/10.1111/2041-210X.12388

Blonder, B., Morrow, C. B., Maitner, B., Harris, D. J., Lamanna, C., Violle, C., Enquist, B. J., & Kerkhoff, A. J. (2018). New approaches for delineating n-dimensional hypervolumes. Methods in Ecology and Evolution, 9(2), 305–319. https://doi.org/10.1111/2041-210X.12865

Cadotte, M. W., Carscadden, K., & Mirotchnick, N. (2011). Beyond species: Functional diversity and the maintenance of ecological processes and services. Journal of Applied Ecology, 48(5), 1079–1087. https://doi.org/10.1111/j.1365-2664.2011.02048.x

Cardoso, P., Mammola, S., Rigal, F., & Carvalho, J. (2022). BAT: Biodiversity Assessment Tools. R package version 2.8.1. https://CRAN.R-project.org/package=BAT

Cardoso, P., Rigal, F., & Carvalho, J. C. (2015). BAT – Biodiversity Assessment Tools, an R package for the measurement and estimation of alpha and beta taxon, phylogenetic and functional diversity. Methods in Ecology and Evolution, 6(2), 232–236. https://doi.org/10.1111/2041-210X.12310

Cardoso, P., Rigal, F., Carvalho, J. C., Fortelius, M., Borges, P. A. V., Podani, J., & Schmera, D. (2014). Partitioning taxon, phylogenetic and functional beta diversity into replacement and richness difference components. Journal of Biogeography, 41(4), 749–761. https://doi.org/10.1111/jbi.12239

Carmona, C. P., Azcárate, F. M., de Bello, F., Ollero, H. S., Lepš, J., & Peco, B. (2012). Taxonomical and functional diversity turnover in Mediterranean grasslands: Interactions between grazing, habitat type and rainfall. Journal of Applied Ecology, 49(5), 1084–1093. https://doi.org/10.1111/j.1365-2664.2012.02193.x

Carvalho, J. C., Cardoso, P., Borges, P. A. V., Schmera, D., & Podani, J. (2013). Measuring fractions of beta diversity and their relationships to nestedness: A theoretical and empirical comparison of novel approaches. Oikos, 122(6), 825–834. https://doi.org/10.1111/j.1600-0706.2012.20980.x

Carvalho, J. C., Cardoso, P., & Gomes, P. (2012). Determining the relative roles of species replacement and species richness differences in generating beta-diversity patterns. Global Ecology and Biogeography, 21(7), 760–771. https://doi.org/10.1111/j.1466-8238.2011.00694.x

Chao, A., Chazdon, R. L., Colwell, R. K., & Shen, T.-J. (2005). A new statistical approach for assessing similarity of species composition with incidence and abundance data. Ecology Letters, 8(2), 148–159. https://doi.org/10.1111/j.1461-0248.2004.00707.x

Chao, A., Chiu, C.-H., Villéger, S., Sun, I.-F., Thorn, S., Lin, Y.-C., Chiang, J.-M., & Sherwin, W. B. (2019). An attribute-diversity approach to functional diversity, functional beta diversity, and related (dis)similarity measures. Ecological Monographs, 89(2), e01343. https://doi.org/10.1002/ecm.1343

Cornwell, W. K., Schwilk, D. W., & Ackerly, D. D. (2006). A trait-based test for habitat filtering: Convex hull volume. Ecology, 87(6), 1465–1471. https://doi.org/10.1890/0012-9658(2006)87[1465:ATTFHF]2.0.CO;2

Deane, D. C., Xing, D., Hui, C., McGeoch, M., & He, F. (2022). A null model for quantifying the geometric effect of habitat subdivision on species diversity. Global Ecology and Biogeography, 31(3), 440–453. https://doi.org/10.1111/geb.13437

Devictor, V., Mouillot, D., Meynard, C., Jiguet, F., Thuiller, W., & Mouquet, N. (2010). Spatial mismatch and congruence between taxonomic, phylogenetic and functional diversity: The need for integrative conservation strategies in a changing world. Ecology Letters, 13(8), 1030–1040.

Díaz, S., Kattge, J., Cornelissen, J. H. C., Wright, I. J., Lavorel, S., Dray, S., Reu, B., Kleyer, M., Wirth, C., Colin Prentice, I., Garnier, E., Bönisch, G., Westoby, M., Poorter, H., Reich, P. B., Moles, A. T., Dickie, J., Gillison, A. N., Zanne, A. E., … Gorné, L. D. (2016). The global spectrum of plant form and function. Nature, 529(7585), 167–171. https://doi.org/10.1038/nature16489

Falster, D. S., Brännström, Å., Westoby, M., & Dieckmann, U. (2017). Multitrait successional forest dynamics enable diverse competitive coexistence. Proceedings of the National Academy of Sciences, 114(13), E2719–E2728.

Falster, D. S., Gallagher, R., Wenk, E. H., Wright, I. J., Indiarto, D., Andrew, S. C., Baxter, C., Lawson, J., Allen, S., Fuchs, A., Monro, A., Kar, F., Adams, M. A., Ahrens, C. W., Alfonzetti, M., Angevin, T., Apgaua, D. M. G., Arndt, S., Atkin, O. K., … Ziemińska, K. (2021). AusTraits, a curated plant trait database for the Australian flora. Scientific Data, 8(1), 254. https://doi.org/10.1038/s41597-021-01006-6

Fitter, A. H., & Peat, H. J. (1994). The Ecological Flora Database. Journal of Ecology, 82(2), 415–425. JSTOR. https://doi.org/10.2307/2261309

Fourdrigniez, M., & Meyer, J.-Y. (2008). Liste et caractéristiques des plantes introduites naturalisées et envahissantes en Polynésie française (No. 17; Contributions à La Biodiversité de Polynésie Française). Délégation à la Recherche, Papeete, 64 pages.

Gotelli, N. J. (2000). NULL MODEL ANALYSIS OF SPECIES CO-OCCURRENCE PATTERNS. Ecology, 81(9), 2606–2621. https://doi.org/10.1890/0012-9658(2000)081[2606:NMAOSC]2.0.CO;2

Gross, N., Bagousse-Pinguet, Y. L., Liancourt, P., Berdugo, M., Gotelli, N. J., & Maestre, F. T. (2017). Functional trait diversity maximizes ecosystem multifunctionality. Nature Ecology & Evolution, 1(5), 0132. https://doi.org/10.1038/s41559-017-0132

Hill, M. O., Preston, C. D., & Roy, D. B. (2004). PLANTATT - Attributes of British and Irish Plants—Spreadsheet. Centre for Ecology and Hydrology.

Hillebrand, H., & Matthiessen, B. (2009). Biodiversity in a complex world: Consolidation and progress in functional biodiversity research. Ecology Letters, 12(12), 1405–1419. https://doi.org/10.1111/j.1461-0248.2009.01388.x

Hui, C., Richardson, D. M., Landi, P., Minoarivelo, H. O., Roy, H. E., Latombe, G., Jing, X., CaraDonna, P. J., Gravel, D., Beckage, B., & Molofsky, J. (2021). Trait positions for elevated invasiveness in adaptive ecological networks. Biological Invasions, 23(6), 1965–1985. https://doi.org/10.1007/s10530-021-02484-w

Irl, S. D. H., Schweiger, A. H., Medina, F. M., Fernández-Palacios, J. M., Harter, D. E. V., Jentsch, A., Provenzale, A., Steinbauer, M. J., & Beierkuhnlein, C. (2017). An island view of endemic rarity—Environmental drivers and consequences for nature conservation. Diversity and Distributions, 23(10), 1132–1142. https://doi.org/10.1111/ddi.12605

Jaccard, P. (1908). Nouvelles recherches sur la distribution florale. Bull. Soc. Vaud. Sci. Nat., 44, 223–270.

Kattge, J., Bönisch, G., Díaz, S., Lavorel, S., Prentice, I. C., Leadley, P., Tautenhahn, S., Werner, G. D. A., Aakala, T., Abedi, M., Acosta, A. T. R., Adamidis, G. C., Adamson, K., Aiba, M., Albert, C. H., Alcántara, J. M., Alcázar C, C., Aleixo, I., Ali, H., … Wirth, C. (2020). TRY plant trait database – enhanced coverage and open access. Global Change Biology, 26(1), 119–188. https://doi.org/10.1111/gcb.14904

Kattge, J., Díaz, S., Lavorel, S., Prentice, I. C., Leadley, P., Bönisch, G., Garnier, E., Westoby, M., Reich, P. B., Wright, I. J., Cornelissen, J. H. C., Violle, C., Harrison, S. P., Van Bodegom, P. M., Reichstein, M., Enquist, B. J., Soudzilovskaia, N. A., Ackerly, D. D., Anand, M., … Wirth, C. (2011). TRY – a global database of plant traits. Global Change Biology, 17(9), 2905–2935. https://doi.org/10.1111/j.1365-2486.2011.02451.x

Kleyer, M., Bekker, R. M., Knevel, I. C., Bakker, J. P., Thompson, K., Sonnenschein, M., Poschlod, P., Van Groenendael, J. M., Klimeš, L., Klimešová, J., Klotz, S., Rusch, G. M., Hermy, M., Adriaens, D., Boedeltje, G., Bossuyt, B., Dannemann, A., Endels, P., Götzenberger, L., … Peco, B. (2008). The LEDA Traitbase: A database of life-history traits of the Northwest European flora. Journal of Ecology, 96(6), 1266–1274. https://doi.org/10.1111/j.1365-2745.2008.01430.x

Kunin, W. E., Harte, J., He, F., Hui, C., Jobe, R. T., Ostling, A., Polce, C., Šizling, A., Smith, A. B., Smith, K., Storch, D., Even, T., Karl-Inne, U., Werner, U., & Varun, V. (2018). Upscaling biodiversity: Estimating the species–area relationship from small samples. Ecological Monographs, 88, 170–187.

Laureto, L. M. O., Cianciaruso, M. V., & Samia, D. S. M. (2015). Functional diversity: An overview of its history and applicability. Natureza & Conservação, 13(2), 112–116. https://doi.org/10.1016/j.ncon.2015.11.001

Loiseau, N., Legras, G., Gaertner, J., Verley, P., Chabanet, P., & Mérigot, B. (2017). Performance of partitioning functional beta-diversity indices: Influence of functional representation and partitioning methods. Global Ecology and Biogeography, 26(6), 753–762.

MacArthur, R. (1970). Species packing and competitive equilibrium for many species. Theoretical Population Biology, 1(1), 1–11.

Maire, E., Grenouillet, G., Brosse, S., & Villéger, S. (2015). How many dimensions are needed to accurately assess functional diversity? A pragmatic approach for assessing the quality of functional spaces. Global Ecology and Biogeography, 24(6), 728–740. https://doi.org/10.1111/geb.12299

Maitner, B. (2022). BIEN: Tools for Accessing the Botanical Information and Ecology Network Database. R package version 1.2.5. https://CRAN.R-project.org/package=BIEN

Mammola, S., & Cardoso, P. (2020). Functional diversity metrics using kernel density n-dimensional hypervolumes. Methods in Ecology and Evolution, 11(8), 986–995. https://doi.org/10.1111/2041-210X.13424

McGill, B. J., Enquist, B. J., Weiher, E., & Westoby, M. (2006). Rebuilding community ecology from functional traits. Trends in Ecology & Evolution, 21(4), 178–185.

Middleton-Welling, J., Dapporto, L., García-Barros, E., Wiemers, M., Nowicki, P., Plazio, E., Bonelli, S., Zaccagno, M., Šašić, M., Liparova, J., Schweiger, O., Harpke, A., Musche, M., Settele, J., Schmucki, R., & Shreeve, T. (2020). A new comprehensive trait database of European and Maghreb butterflies, Papilionoidea. Scientific Data, 7(1), 351. https://doi.org/10.1038/s41597-020-00697-7

Molofsky, J., Park, D. S., Richardson, D. M., Keller, S. R., Beckage, B., Mandel, J. R., Boatwright, J. S., & Hui, C. (2022). Optimal differentiation to the edge of trait space (EoTS). Evolutionary Ecology, 36(5), 743–752. https://doi.org/10.1007/s10682-022-10192-7

Mouillot, D., Loiseau, N., Grenié, M., Algar, A. C., Allegra, M., Cadotte, M. W., Casajus, N., Denelle, P., Guéguen, M., Maire, A., Maitner, B., McGill, B. J., McLean, M., Mouquet, N., Munoz, F., Thuiller, W., Villéger, S., Violle, C., & Auber, A. (2021). The dimensionality and structure of species trait spaces. Ecology Letters, 24(9), 1988–2009. https://doi.org/10.1111/ele.13778

Mouillot, D., Stubbs, W., Faure, M., Dumay, O., Tomasini, J. A., Wilson, J. B., & Chi, T. D. (2005). Niche overlap estimates based on quantitative functional traits: A new family of non-parametric indices. Oecologia, 145(3), 345–353. https://doi.org/10.1007/s00442-005-0151-z

Petchey, O. L., & Gaston, K. J. (2006). Functional diversity: Back to basics and looking forward. Ecology Letters, 9(6), 741–758. https://doi.org/10.1111/j.1461-0248.2006.00924.x

R Core Team. (2022). R: A language and environment for statistical computing. R Foundation for Statistical Computing, Vienna, Austria. https://www.R-project.org/

Siefert, A., Ravenscroft, C., Weiser, M. D., & Swenson, N. G. (2013). Functional beta-diversity patterns reveal deterministic community assembly processes in eastern North American trees. Global Ecology and Biogeography, 22(6), 682–691. https://doi.org/10.1111/geb.12030

Sobral, F. L., Lees, A. C., & Cianciaruso, M. V. (2016). Introductions do not compensate for functional and phylogenetic losses following extinctions in insular bird assemblages. Ecology Letters, 19(9), 1091–1100. https://doi.org/10.1111/ele.12646

Socolar, J. B., Gilroy, J. J., Kunin, W. E., & Edwards, D. P. (2016). How should beta-diversity inform biodiversity conservation? Trends in Ecology & Evolution, 31(1), 67–80.

Swenson, N. G., Erickson, D. L., Mi, X., Bourg, N. A., Forero-Montaña, J., Ge, X., Howe, R., Lake, J. K., Liu, X., Ma, K., Pei, N., Thompson, J., Uriarte, M., Wolf, A., Wright, S. J., Ye, W., Zhang, J., Zimmerman, J. K., & Kress, W. J. (2012). Phylogenetic and functional alpha and beta diversity in temperate and tropical tree communities. Ecology, 93(sp8), S112–S125. https://doi.org/10.1890/11-0402.1

Tavşanoğlu, Ç., & Pausas, J. G. (2018). A functional trait database for Mediterranean Basin plants. Scientific Data, 5(1), 180135. https://doi.org/10.1038/sdata.2018.135

Tecco, P. A., Díaz, S., Cabido, M., & Urcelay, C. (2010). Functional traits of alien plants across contrasting climatic and land-use regimes: Do aliens join the locals or try harder than them? Journal of Ecology, 98(1), 17–27.

Tobias, J. A., Sheard, C., Pigot, A. L., Devenish, A. J. M., Yang, J., Sayol, F., Neate-Clegg, M. H. C., Alioravainen, N., Weeks, T. L., Barber, R. A., Walkden, P. A., MacGregor, H. E. A., Jones, S. E. I., Vincent, C., Phillips, A. G., Marples, N. M., Montaño-Centellas, F. A., Leandro-Silva, V., Claramunt, S., … Schleuning, M. (2022). AVONET: morphological, ecological and geographical data for all birds. Ecology Letters, 25(3), 581–597. https://doi.org/10.1111/ele.13898

Villéger, S., Grenouillet, G., & Brosse, S. (2013). Decomposing functional β-diversity reveals that low functional β-diversity is driven by low functional turnover in European fish assemblages. Global Ecology and Biogeography, 22(6), 671–681. https://doi.org/10.1111/geb.12021

Villéger, S., Maire, E., & Leprieur, F. (2017). On the risks of using dendrograms to measure functional diversity and multidimensional spaces to measure phylogenetic diversity: A comment on Sobral et al. (2016). Ecology Letters, 20(4), 554–557. https://doi.org/10.1111/ele.12750

Villéger, S., Mason, N. W. H., & Mouillot, D. (2008). New multidimensional functional diversity indices for a multifaceted framework in functional ecology. Ecology, 89(8), 2290–2301. https://doi.org/10.1890/07-1206.1

Westoby, M. (1998). A leaf-height-seed (LHS) plant ecology strategy scheme. Plant and Soil, 199(2), 213–227. https://doi.org/10.1023/A:1004327224729

Williams, P. H. (1996). Mapping variations in the strength and breadth of biogeographic transition zones using species turnover. Proceedings of the Royal Society B: Biological Sciences, 263(1370), 579–588. https://doi.org/10.1098/rspb.1996.0087

Wohlwend, M. R., Craven, D., Weigelt, P., Seebens, H., Winter, M., Kreft, H., Dawson, W., Essl, F., van Kleunen, M., & Pergl, J. (2021). Data descriptor: Pacific introduced flora (PaciFLora). Biodiversity Data Journal, 9.

