## Supplementary material for "A kernel integral method to remove biases in estimating trait turnover": Supplemetary Material

**Appendix A.** Methodological figures

**Appendix B.** Graphical outputs underlying the computation of the β diversity indices under the different functional-space approaches

**Appendix C.** Supplementary results for the theoretical simulations

**Appendix D.** Supplementary results for French Polynesia data

**Appendix A.** Methodological figures

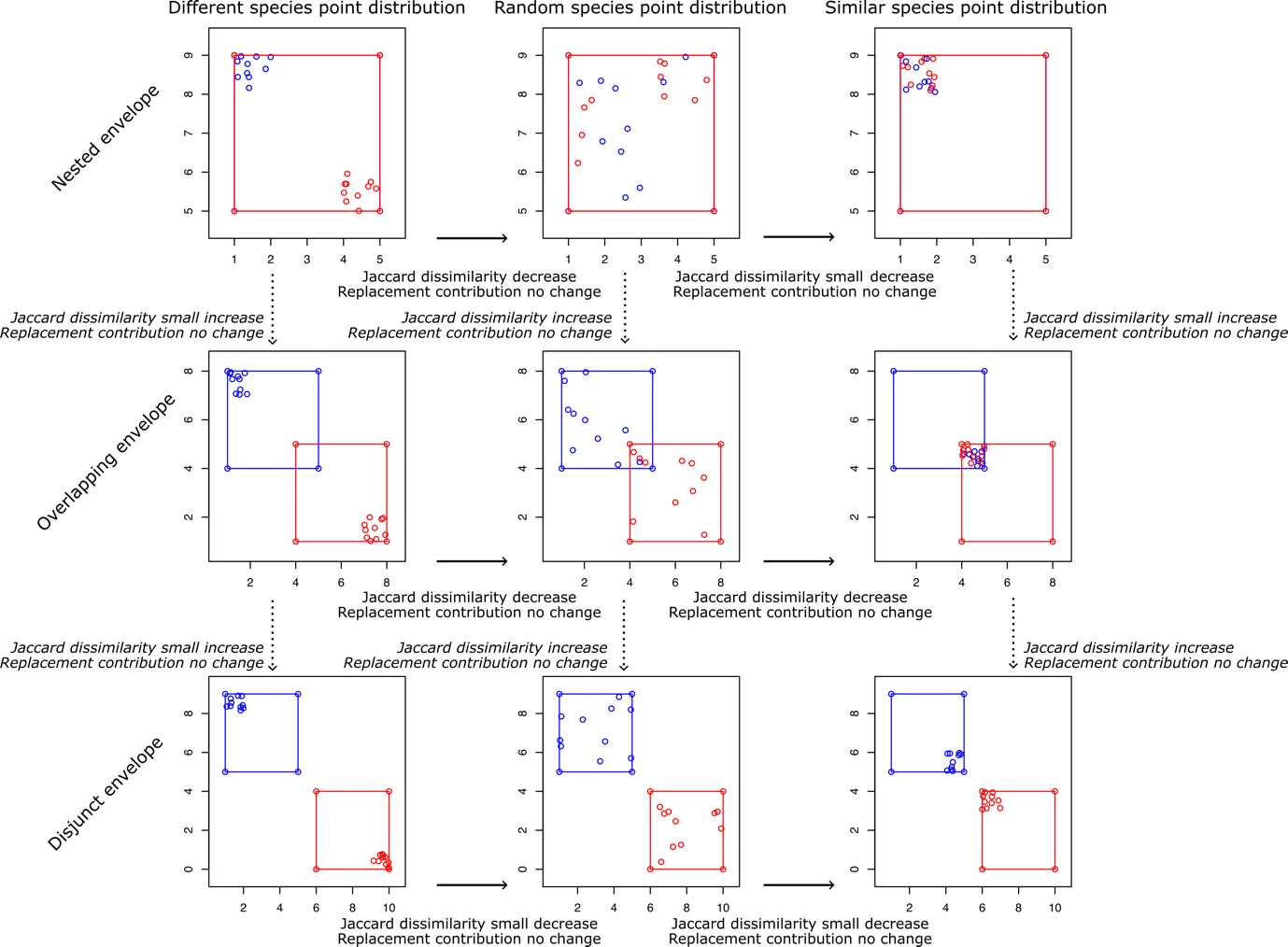

**Figure A1.** The nine theoretical configurations of pairs of communities for the same size of the MCPs, using only 10 randomly drawn points for clarity. We varied how the MCPs overlapped (“nested”, “overlapping” or “disjunct”), and how the points are distributed within the MCPs (“different” – distributed in opposite corners of the MCPs –, “random” – randomly distributed within the MCPs – or “similar” – distributed in the closest corners of the MCPs). These were repeated for 10, 40, 70 and 100 random points, and for 50 replicates.

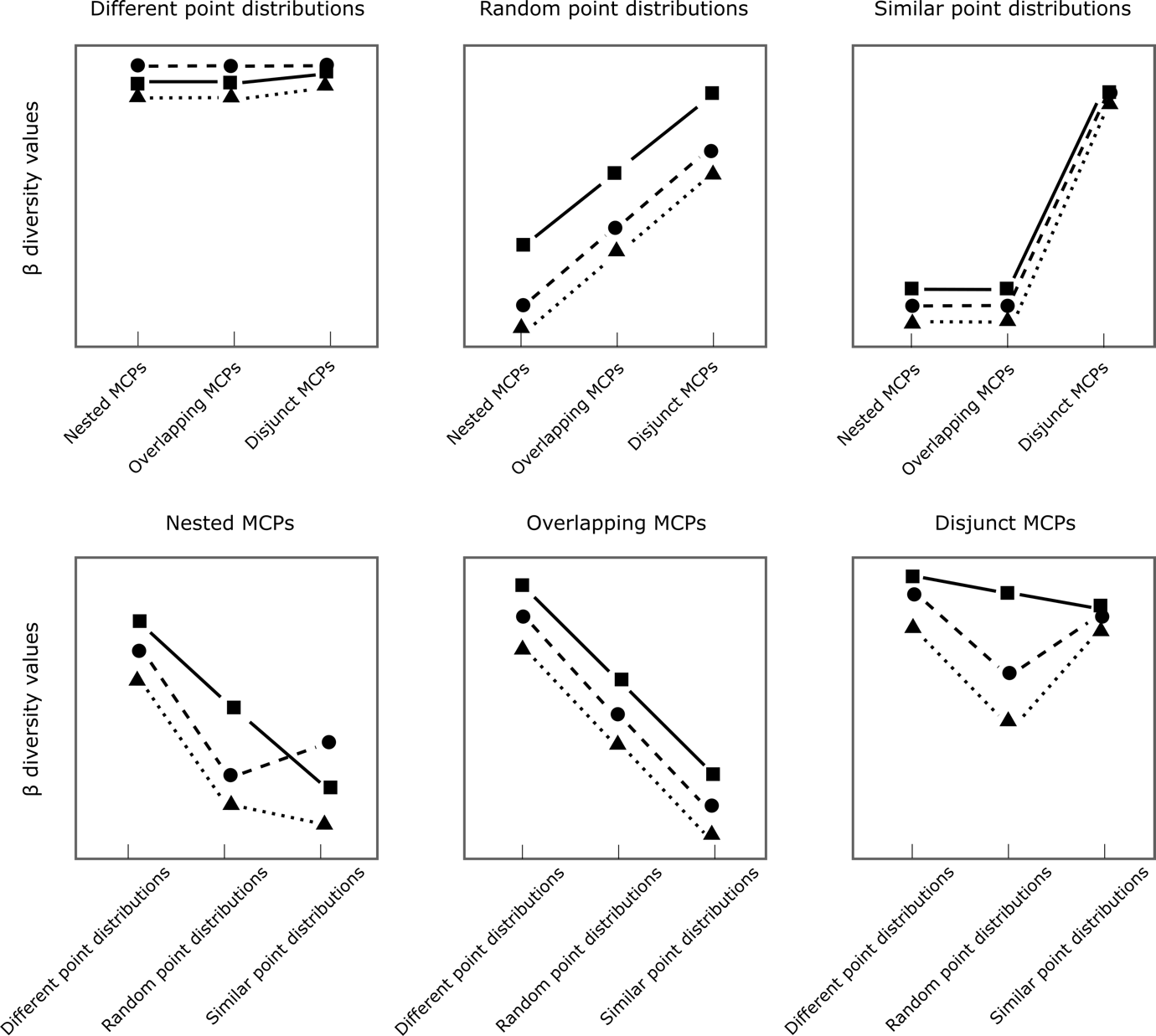

**Figure A2.** Expected qualitative differences in β diversity values for different configurations of species points in the functional space when the MCPs of the two communities have different sizes. Jaccard dissimilarity (squares, solid lines), Williams replacement (triangles, dotted lines), and contribution of replacement to overall turnover, computed as Williams replacement divided by Jaccard dissimilarity (circles, dashed lines).

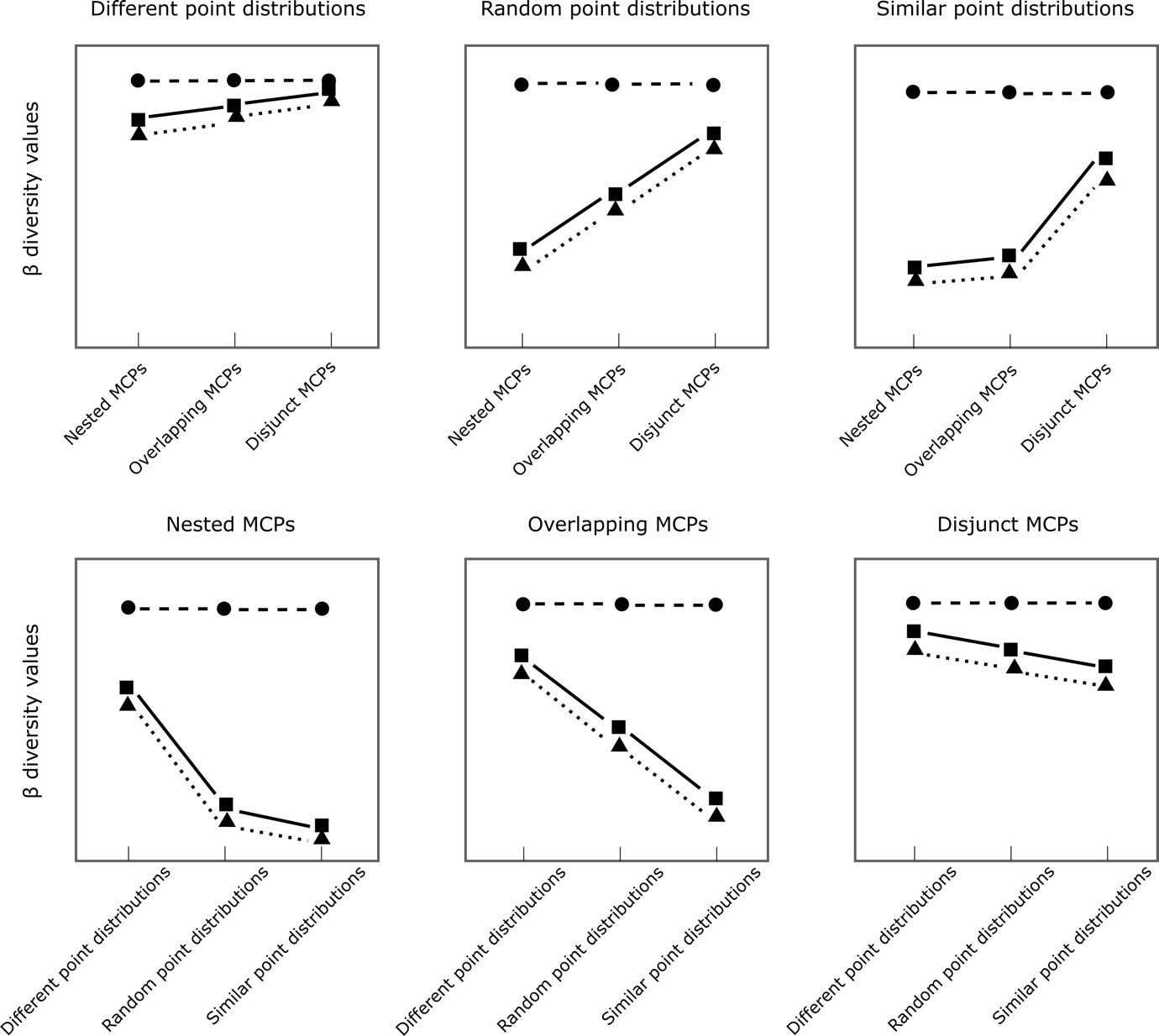

**Figure A3.** Expected qualitative differences in β diversity values for different configurations of species points in the functional space when the MCPs of the two communities have the same size. Jaccard dissimilarity (squares, solid lines), Williams replacement (triangles, dotted lines), and contribution of replacement to overall turnover, computed as Williams replacement divided by Jaccard dissimilarity (circles, dashed lines).

**Table A1.** Justifications for expected changes in functional turnover for the different species point configurations in the functional space. Blue and red communities refer to the colours in Figures 4 and 5.

| Transition | Expectations for Jaccard dissimilarity | Justification | Expectations for Williams replacement (%) | Justification |
| --- | --- | --- | --- | --- |
| 1. **Different size Minimum Convex Polytopes (MCPs)** | | | | |
| Different: Nested → Overlapping | No change | Most of the species points move away from each other, but this is compensated by the fact that the species points in the red community get closer to the extreme point in the bottom-right of the blue community. | No change | The area of the functional space covered by the two communities is very similar, and only differs for three species, and turnover is caused almost exclusively by replacement. |
| Different: Overlapping → Similar | Small increase | The species points from the two communities move away from each other, but since the dissimilarity was already high, the increase is small. | No change | The area of the functional space covered by the two communities is very similar, and only differs for three species, and turnover is caused almost exclusively by replacement. |
| Random: Nested → Overlapping | Increase | The overlap between the point distributions from the two communities decreases. | Increase | The turnover was first to differences in functional area covered by the species points in the nested configuration, and is now due in part to differences in overlapping of the point distribution. |
| Random: Overlapping → Similar | Increase | The overlap between the point distributions from the two communities decreases. | Increase | Turnover is high because the parts of the functional space covered by the species points are disjunct, not because of differences in area covered. |
| Similar: Nested → Overlapping | No change | The overlap between the point distributions from the two communities remains constant despite changing position in the functional space. | No change | The area of the functional space covered by the two communities is very similar, and only differs for three species, and turnover is caused almost exclusively by replacement. |
| Similar: Overlapping → Similar | Increase | The overlap between the point distributions from the two communities decreases. | Increase | Turnover is high because the parts of the functional space covered by the species points are disjunct, not because of differences in area covered. |
| Nested: Different → Random | Decrease | The point distributions overlap more because species points in the blue community will now fall in the red MCP. | Decrease | Most of the turnover was due to different parts of the functional space being covered by species points to differences in covered area. |
| Nested: Random → Similar | Decrease | The point distributions overlap more because the species points all fall in the same small area of the functional space. | Small increase | The areas covered by the species points from the two communities becomes more similar to each other, and therefore proportionally replacement contributes more to turnover. |
| Overlapping: Different → Random | Decrease | The point distributions overlap more because some species points in the blue and red communities will now fall in the overlapping area of the MCPs. | Decrease | The part of the functional space covered by the two communities was mostly disjunct and due for the most part to replacement. As it now overlaps, the contribution of functional richness difference increase, so the contribution of replacement decreases. |
| Overlapping: Random → Similar | Decrease | The point distributions overlap more because all species points in the blue and red communities will now fall in the overlapping area of the MCPs. | Decrease | Turnover is very low, and mostly due to the three extreme species on the edge of each MCP, which cause a difference in functional richness. The contribution of species replacement therefore decreases. |
| Disjunct: Different → Random | Small decrease | The point distributions are still not overlapping, but species points are now closer to each other and the distributions are therefore slightly more similar. | Decrease | In both cases, the distributions of the species points cover different parts of the functional space, but in the random case the covered area differ much more. Functional richness contribution therefore increases, whereas the contribution of replacement decreases. |
| Disjunct: Random → Similar | Small decrease | The point distributions are still not overlapping, but species points are now closer to each other and the distributions are therefore slightly more similar. | Increase | The area covered by each set of species points become much more similar and therefore turnover becomes mostly due to replacement. |
| 2. **Same size Minimum Convex Polytopes** | | | | |
| Different: Nested → Overlapping | Small increase | The point distributions from the two communities are far from each other, and the small increase is due to the extreme point species moving away from each other. | No change | The area of the functional space covered by the two communities is the same, and turnover is caused almost exclusively by replacement. |
| Different: Overlapping → Similar | Small increase | The point distributions from the two communities are far from each other, and the small increase is due to the extreme point species moving away from each other. | No change | The area of the functional space covered by the two communities is the same, and turnover is caused almost exclusively by replacement. |
| Random: Nested → Overlapping | Increase | The overlap between the point distributions from the two communities decreases. | No change | The area of the functional space covered by the two communities is the same, and turnover is caused almost exclusively by replacement. |
| Random: Overlapping → Similar | Increase | The overlap between the point distributions from the two communities decreases. | No change | The area of the functional space covered by the two communities is the same, and turnover is caused almost exclusively by replacement. |
| Similar: Nested → Overlapping | Small increase | The overlap between the point distributions from the two communities remains similar for most species points, but the small increase is due to the extreme point species moving away from each other. | No change | The area of the functional space covered by the two communities is the same, and turnover is caused almost exclusively by replacement. |
| Similar: Overlapping → Similar | Increase | The overlap between the point distributions from the two communities decreases. | No change | The area of the functional space covered by the two communities is the same, and turnover is caused almost exclusively by replacement. |
| Nested: Different → Random | Decrease | The overlap between the point distributions from the two communities increases. | No change | The area of the functional space covered by the two communities is the same, and turnover is caused almost exclusively by replacement. |
| Nested: Random → Similar | Small decrease | The overlap between the point distributions from the two communities increases due to the area in which most species points are distributed getting smaller. | No change | The area of the functional space covered by the two communities is the same, and turnover is caused almost exclusively by replacement. |
| Overlapping: Different → Random | Decrease | The overlap between the point distributions from the two communities increases. | No change | The area of the functional space covered by the two communities is the same, and turnover is caused almost exclusively by replacement. |
| Overlapping: Random → Similar | Decrease | The overlap between the point distributions from the two communities increases. | No change | The area of the functional space covered by the two communities is the same, and turnover is caused almost exclusively by replacement. |
| Disjunct: Different → Random | Small decrease | The point distributions from the two communities increases, but the species points do not overlap, hence only a small decrease. | No change | The area of the functional space covered by the two communities is the same, and turnover is caused almost exclusively by replacement. |
| Disjunct: Random → Similar | Small decrease | The point distributions from the two communities increases, but the species points do not overlap, hence only a small decrease. | No change | The area of the functional space covered by the two communities is the same, and turnover is caused almost exclusively by replacement. |

**Appendix B.** Graphical outputs underlying the computation of the β diversity indices under the different functional-space approaches

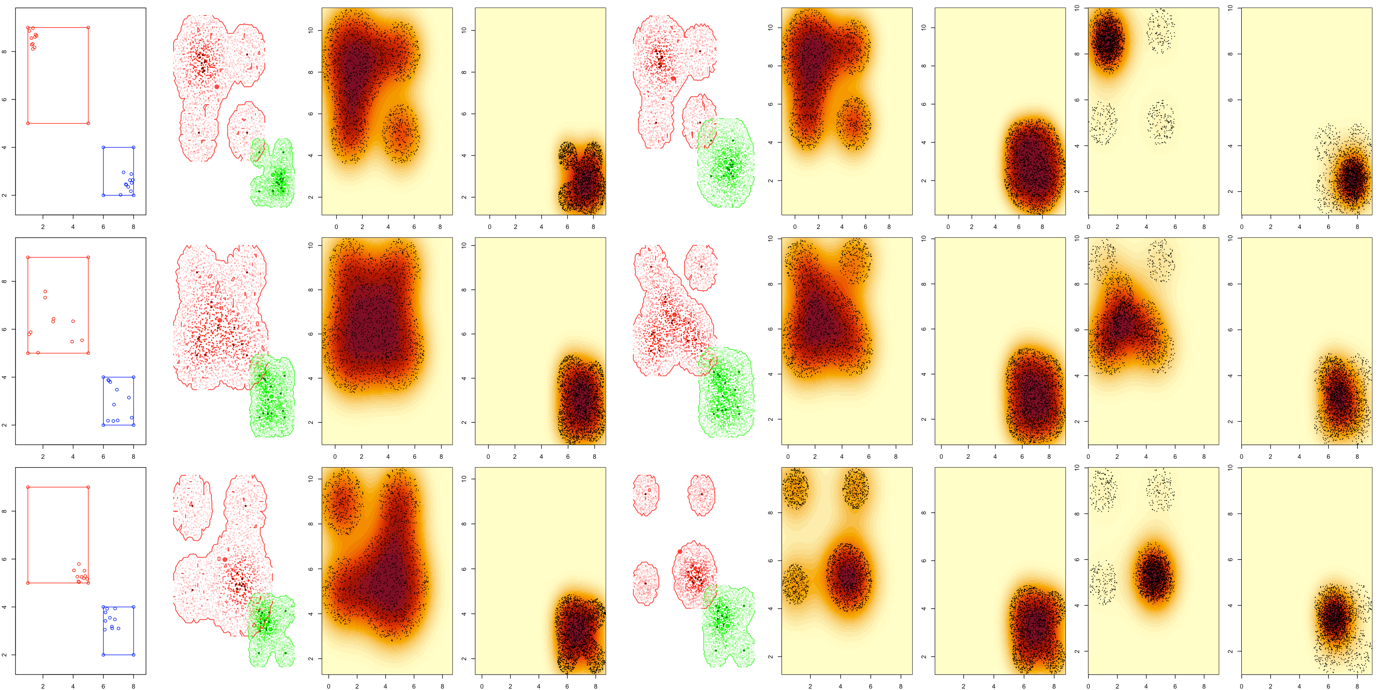

**Figure B1.** A set of theoretical point distribution used to examine the behaviour of the different indices: disjunct MCPs of different sizes, 10 points. In the first row, the points are located far from each other within the MCPs. In the second row, the points are randomly distributed within the MCPs. In the last row, the points are located close each other within the MCPs. The first column shows the point distributions and MCPs in the functional space. The second column shows the polytopes generated by the KDH approach Version 1. The third and fourth columns show the kernels and a sample of the random points from which they are computed, used in the KDH approach Version 1 and the kernel-based approach Version 1. The fifth column shows the polytopes generated by the KDH approach Version 2. The sixth and seventh columns show the kernels and a sample of the random points from which they are computed, used in the KDH approach Version 2 and the kernel-based approach Version 2. The eighth and ninth columns show the kernels and a sample of the random points from which they are computed, used in the kernel-based approach Version 3.

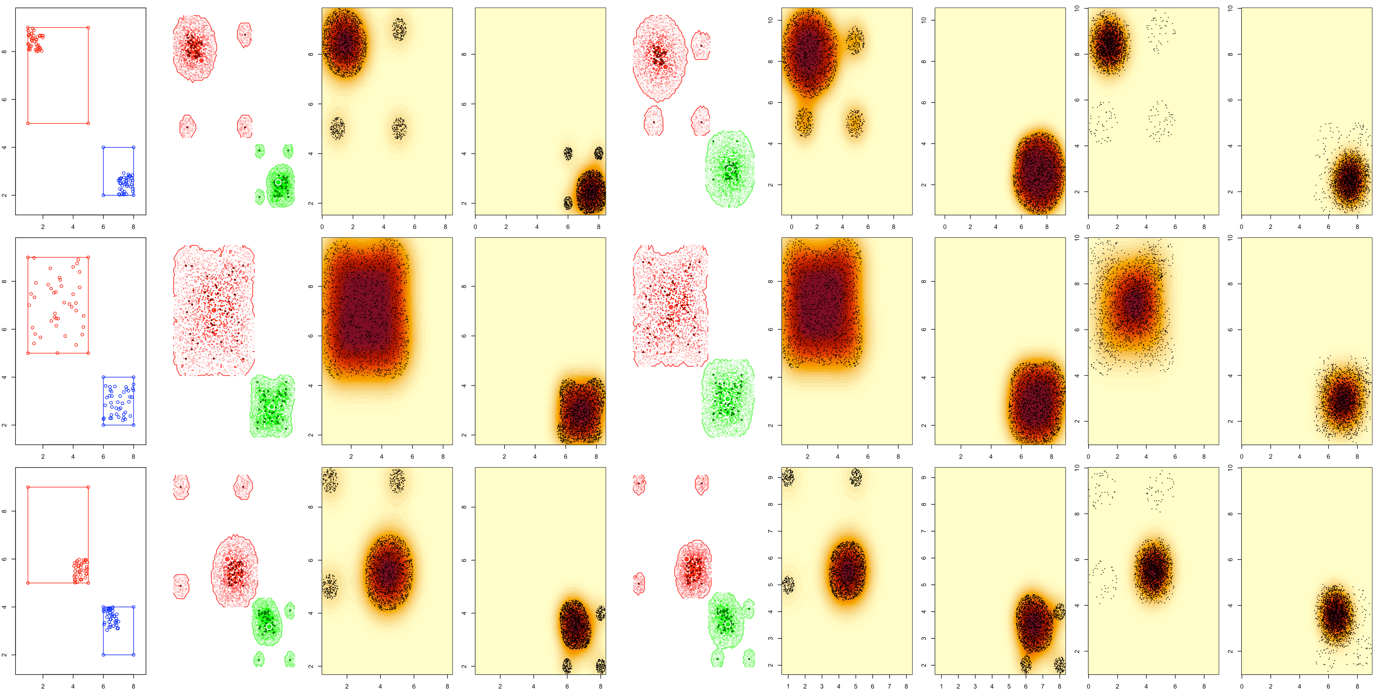

**Figure B2.** A set of theoretical point distribution used to examine the behaviour of the different indices: disjunct MCPs of different sizes, 40 points. In the first row, the points are located far from each other within the MCPs. In the second row, the points are randomly distributed within the MCPs. In the last row, the points are located close each other within the MCPs. The first column shows the point distributions and MCPs in the functional space. The second column shows the polytopes generated by the KDH approach Version 1. The third and fourth columns show the kernels and a sample of the random points from which they are computed, used in the KDH approach Version 1 and the kernel-based approach Version 1. The fifth column shows the polytopes generated by the KDH approach Version 2. The sixth and seventh columns show the kernels and a sample of the random points from which they are computed, used in the KDH approach Version 2 and the kernel-based approach Version 2. The eighth and ninth columns show the kernels and a sample of the random points from which they are computed, used in the kernel-based approach Version 3.

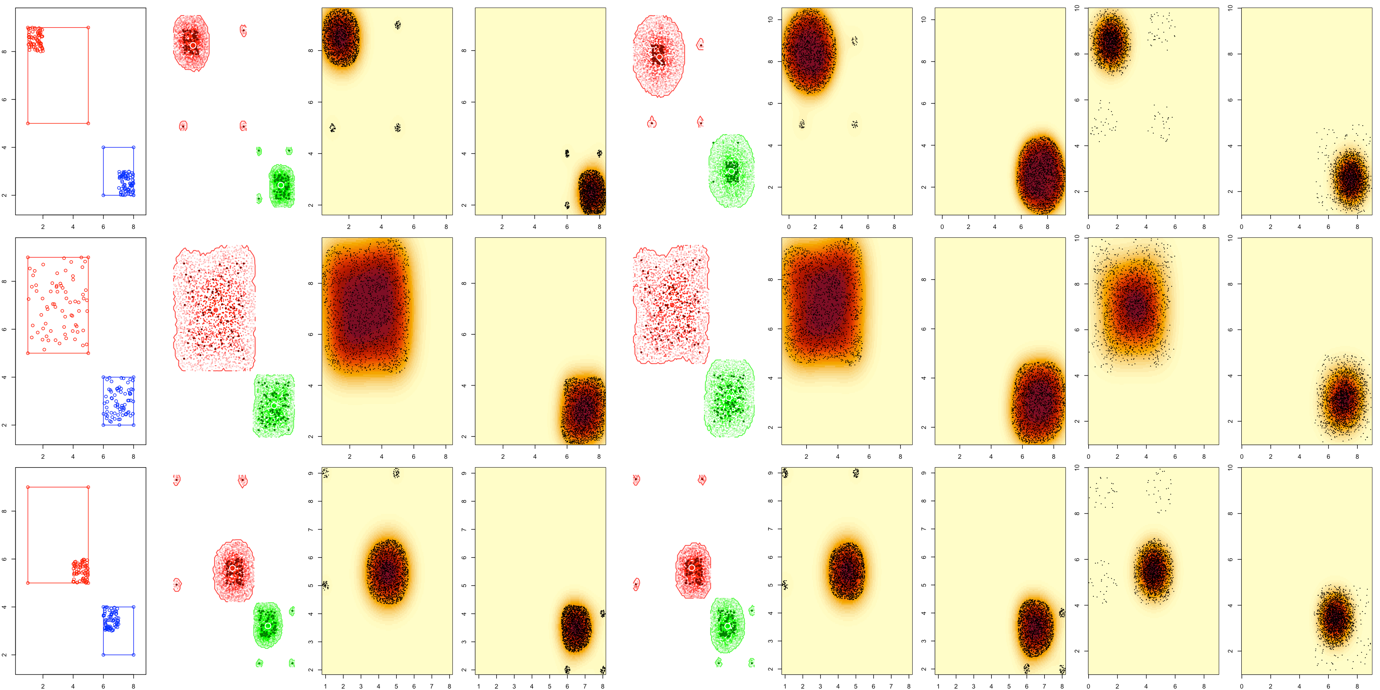

**Figure B3.** A set of theoretical point distribution used to examine the behaviour of the different indices: disjunct MCPs of different sizes, 70 points. In the first row, the points are located far from each other within the MCPs. In the second row, the points are randomly distributed within the MCPs. In the last row, the points are located close each other within the MCPs. The first column shows the point distributions and MCPs in the functional space. The second column shows the polytopes generated by the KDH approach Version 1. The third and fourth columns show the kernels and a sample of the random points from which they are computed, used in the KDH approach Version 1 and the kernel-based approach Version 1. The fifth column shows the polytopes generated by the KDH approach Version 2. The sixth and seventh columns show the kernels and a sample of the random points from which they are computed, used in the KDH approach Version 2 and the kernel-based approach Version 2. The eighth and ninth columns show the kernels and a sample of the random points from which they are computed, used in the kernel-based approach Version 3.

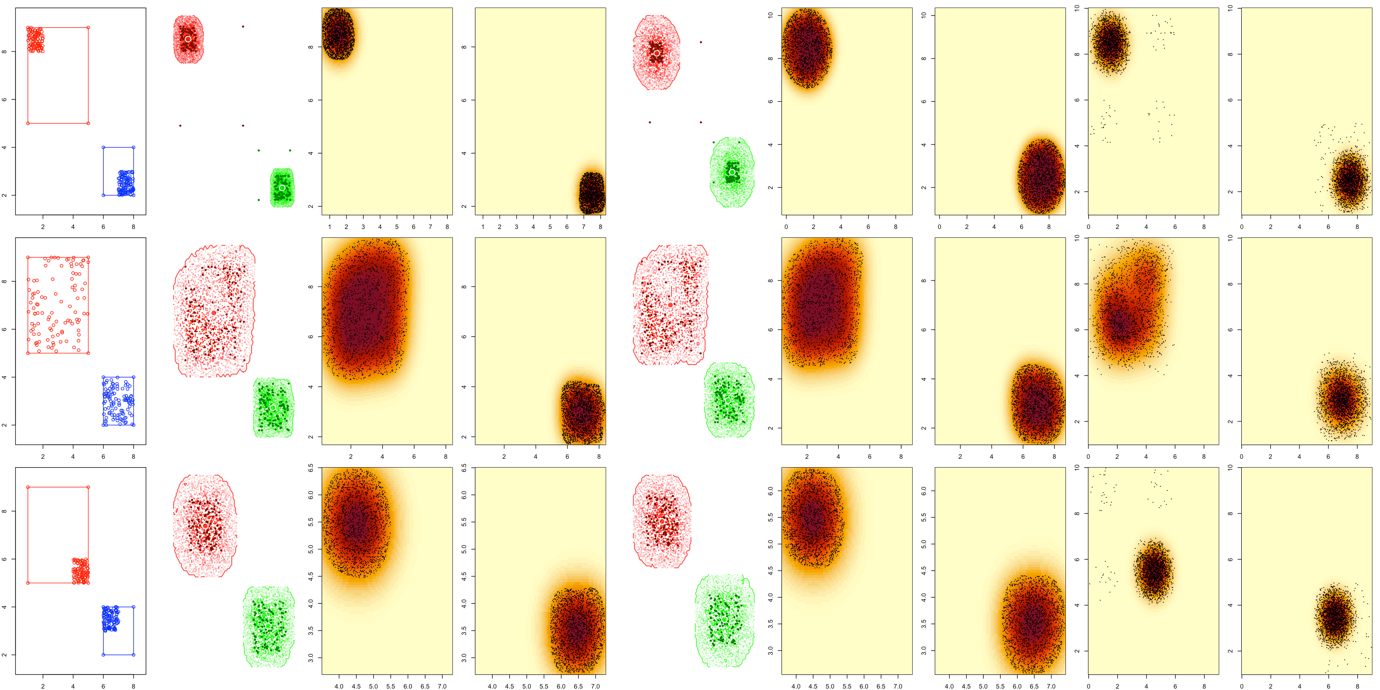

**Figure B4.** A set of theoretical point distribution used to examine the behaviour of the different indices: disjunct MCPs of different sizes, 100 points. In the first row, the points are located far from each other within the MCPs. In the second row, the points are randomly distributed within the MCPs. In the last row, the points are located close each other within the MCPs. The first column shows the point distributions and MCPs in the functional space. The second column shows the polytopes generated by the KDH approach Version 1. The third and fourth columns show the kernels and a sample of the random points from which they are computed, used in the KDH approach Version 1 and the kernel-based approach Version 1. The fifth column shows the polytopes generated by the KDH approach Version 2. The sixth and seventh columns show the kernels and a sample of the random points from which they are computed, used in the KDH approach Version 2 and the kernel-based approach Version 2. The eighth and ninth columns show the kernels and a sample of the random points from which they are computed, used in the kernel-based approach Version 3.

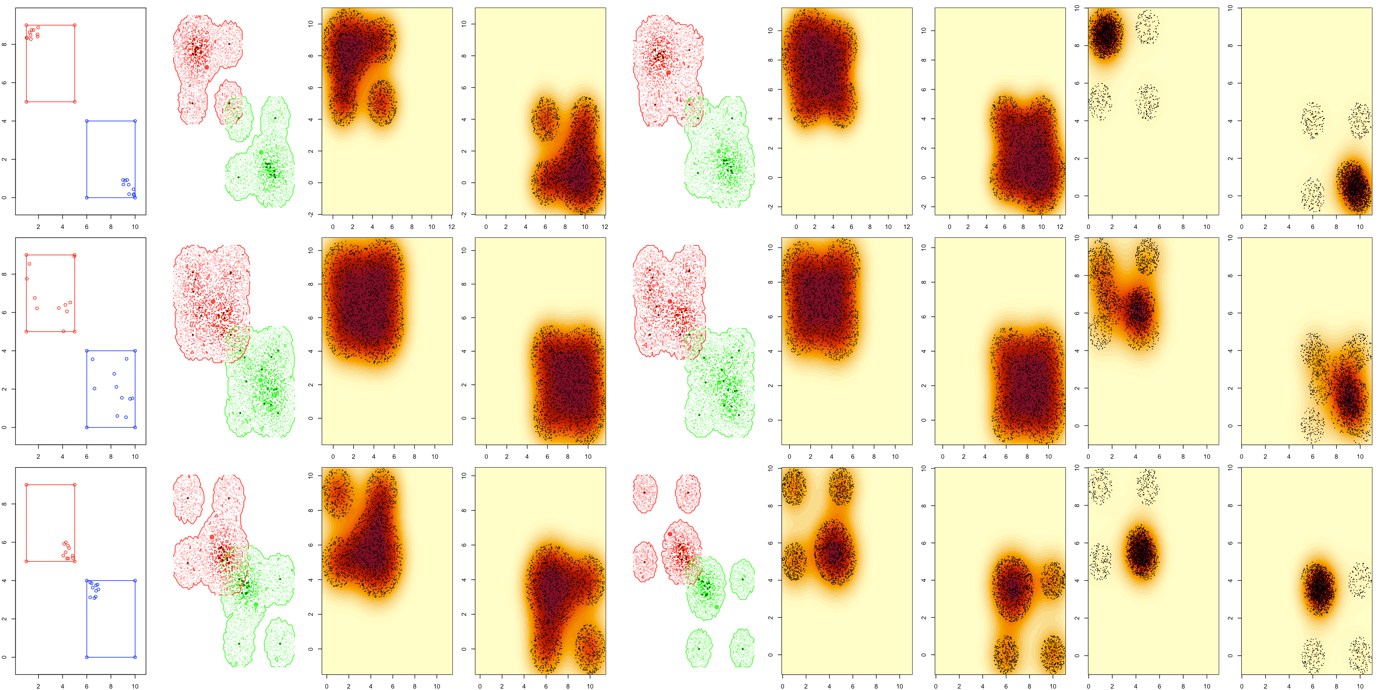

**Figure B5.** A set of theoretical point distribution used to examine the behaviour of the different indices: disjunct MCPs of same size, 10 points. In the first row, the points are located far from each other within the MCPs. In the second row, the points are randomly distributed within the MCPs. In the last row, the points are located close each other within the MCPs. The first column shows the point distributions and MCPs in the functional space. The second column shows the polytopes generated by the KDH approach Version 1. The third and fourth columns show the kernels and a sample of the random points from which they are computed, used in the KDH approach Version 1 and the kernel-based approach Version 1. The fifth column shows the polytopes generated by the KDH approach Version 2. The sixth and seventh columns show the kernels and a sample of the random points from which they are computed, used in the KDH approach Version 2 and the kernel-based approach Version 2. The eighth and ninth columns show the kernels and a sample of the random points from which they are computed, used in the kernel-based approach Version 3.

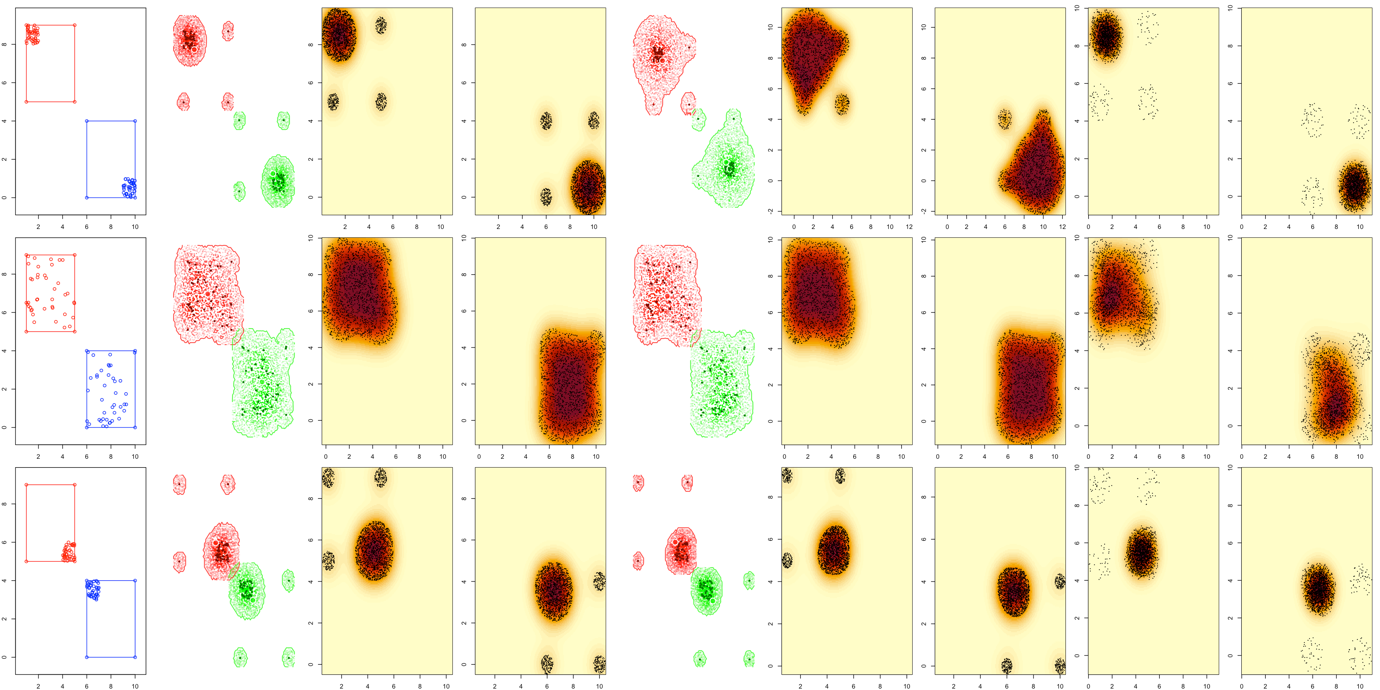

**Figure B6.** A set of theoretical point distribution used to examine the behaviour of the different indices: disjunct MCPs of same size, 40 points. In the first row, the points are located far from each other within the MCPs. In the second row, the points are randomly distributed within the MCPs. In the last row, the points are located close each other within the MCPs. The first column shows the point distributions and MCPs in the functional space. The second column shows the polytopes generated by the KDH approach Version 1. The third and fourth columns show the kernels and a sample of the random points from which they are computed, used in the KDH approach Version 1 and the kernel-based approach Version 1. The fifth column shows the polytopes generated by the KDH approach Version 2. The sixth and seventh columns show the kernels and a sample of the random points from which they are computed, used in the KDH approach Version 2 and the kernel-based approach Version 2. The eighth and ninth columns show the kernels and a sample of the random points from which they are computed, used in the kernel-based approach Version 3.

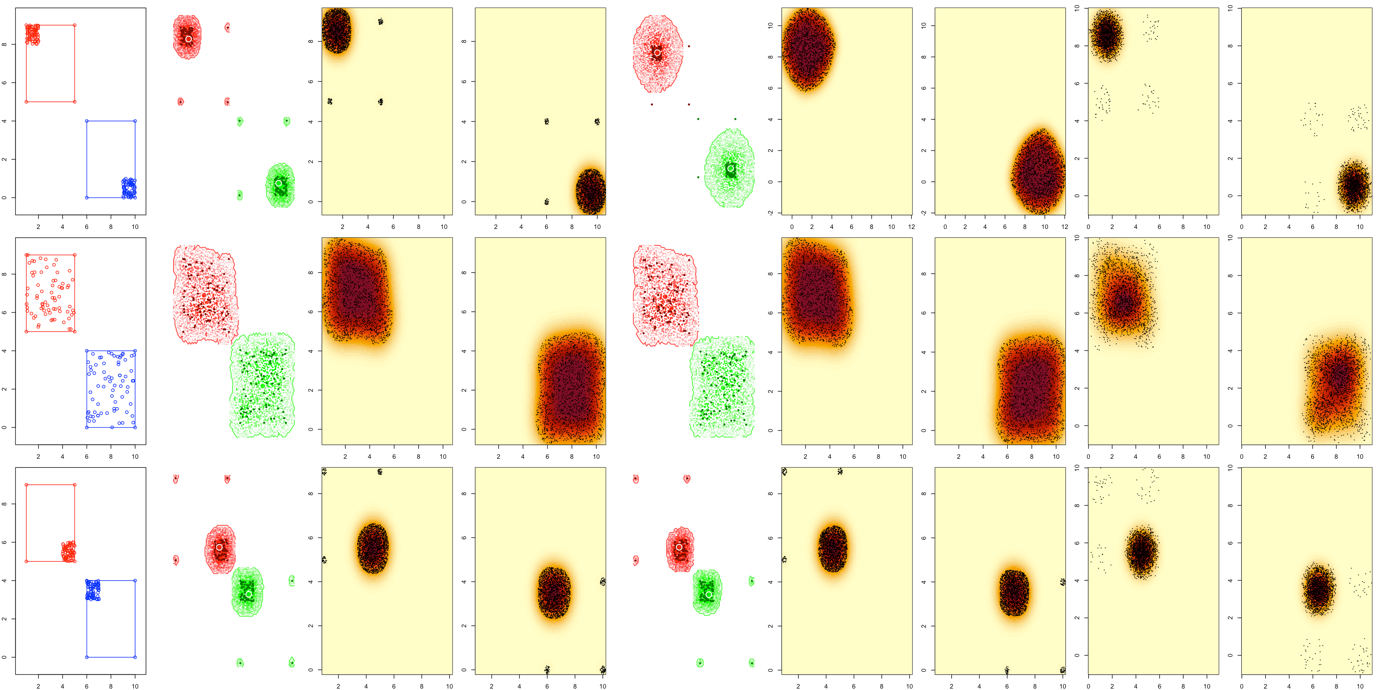

**Figure B7.** A set of theoretical point distribution used to examine the behaviour of the different indices: disjunct MCPs of same size, 70 points. In the first row, the points are located far from each other within the MCPs. In the second row, the points are randomly distributed within the MCPs. In the last row, the points are located close each other within the MCPs. The first column shows the point distributions and MCPs in the functional space. The second column shows the polytopes generated by the KDH approach Version 1. The third and fourth columns show the kernels and a sample of the random points from which they are computed, used in the KDH approach Version 1 and the kernel-based approach Version 1. The fifth column shows the polytopes generated by the KDH approach Version 2. The sixth and seventh columns show the kernels and a sample of the random points from which they are computed, used in the KDH approach Version 2 and the kernel-based approach Version 2. The eighth and ninth columns show the kernels and a sample of the random points from which they are computed, used in the kernel-based approach Version 3.

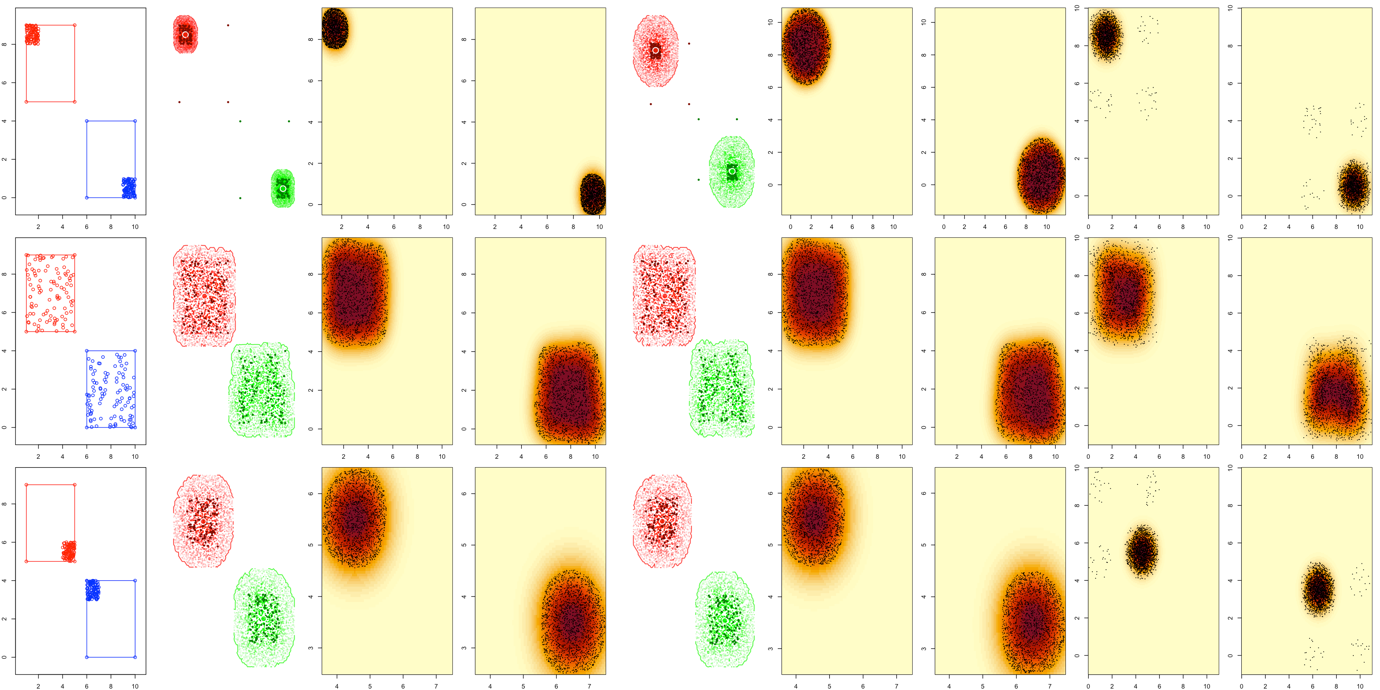

**Figure B8.** A set of theoretical point distribution used to examine the behaviour of the different indices: disjunct MCPs of same size, 100 points. In the first row, the points are located far from each other within the MCPs. In the second row, the points are randomly distributed within the MCPs. In the last row, the points are located close each other within the MCPs. The first column shows the point distributions and MCPs in the functional space. The second column shows the polytopes generated by the KDH approach Version 1. The third and fourth columns show the kernels and a sample of the random points from which they are computed, used in the KDH approach Version 1 and the kernel-based approach Version 1. The fifth column shows the polytopes generated by the KDH approach Version 2. The sixth and seventh columns show the kernels and a sample of the random points from which they are computed, used in the KDH approach Version 2 and the kernel-based approach Version 2. The eighth and ninth columns show the kernels and a sample of the random points from which they are computed, used in the kernel-based approach Version 3.

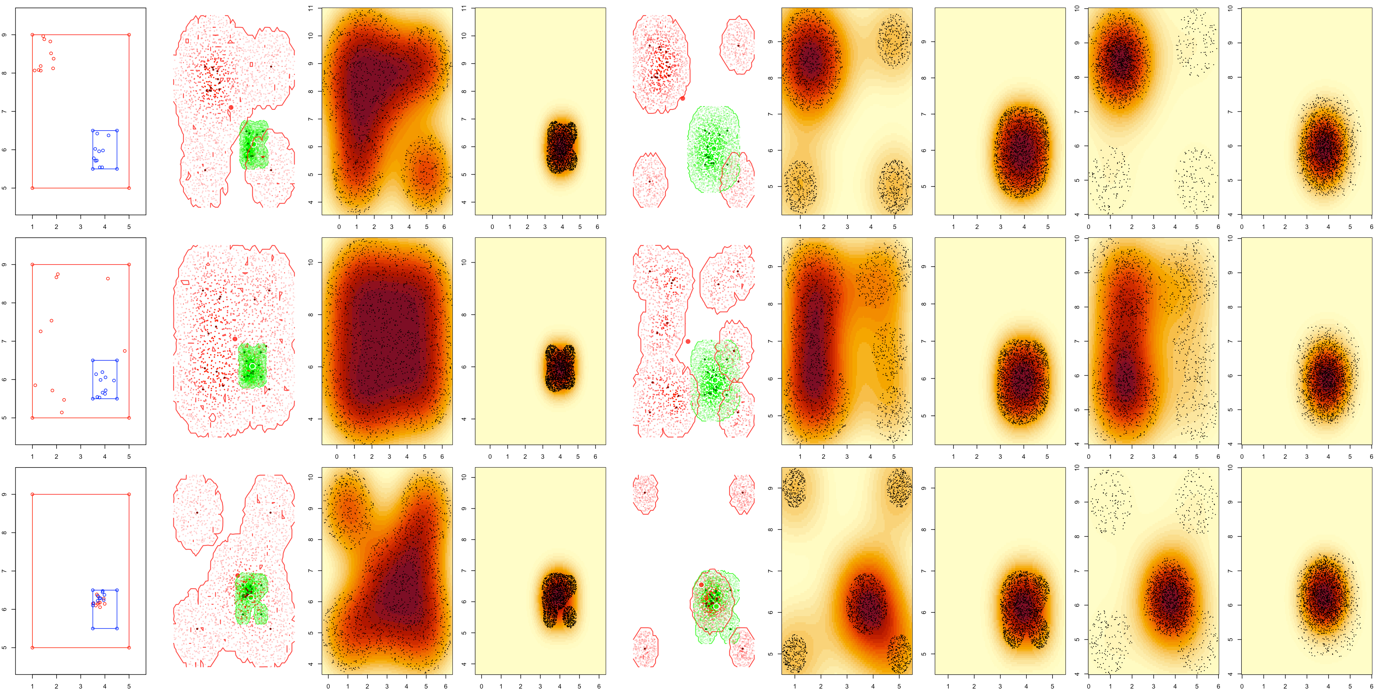

**Figure B9.** A set of theoretical point distribution used to examine the behaviour of the different indices: nested MCPs of different sizes, 10 points. In the first row, the points are located far from each other within the MCPs. In the second row, the points are randomly distributed within the MCPs. In the last row, the points are located close each other within the MCPs. The first column shows the point distributions and MCPs in the functional space. The second column shows the polytopes generated by the KDH approach Version 1. The third and fourth columns show the kernels and a sample of the random points from which they are computed, used in the KDH approach Version 1 and the kernel-based approach Version 1. The fifth column shows the polytopes generated by the KDH approach Version 2. The sixth and seventh columns show the kernels and a sample of the random points from which they are computed, used in the KDH approach Version 2 and the kernel-based approach Version 2. The eighth and ninth columns show the kernels and a sample of the random points from which they are computed, used in the kernel-based approach Version 3.

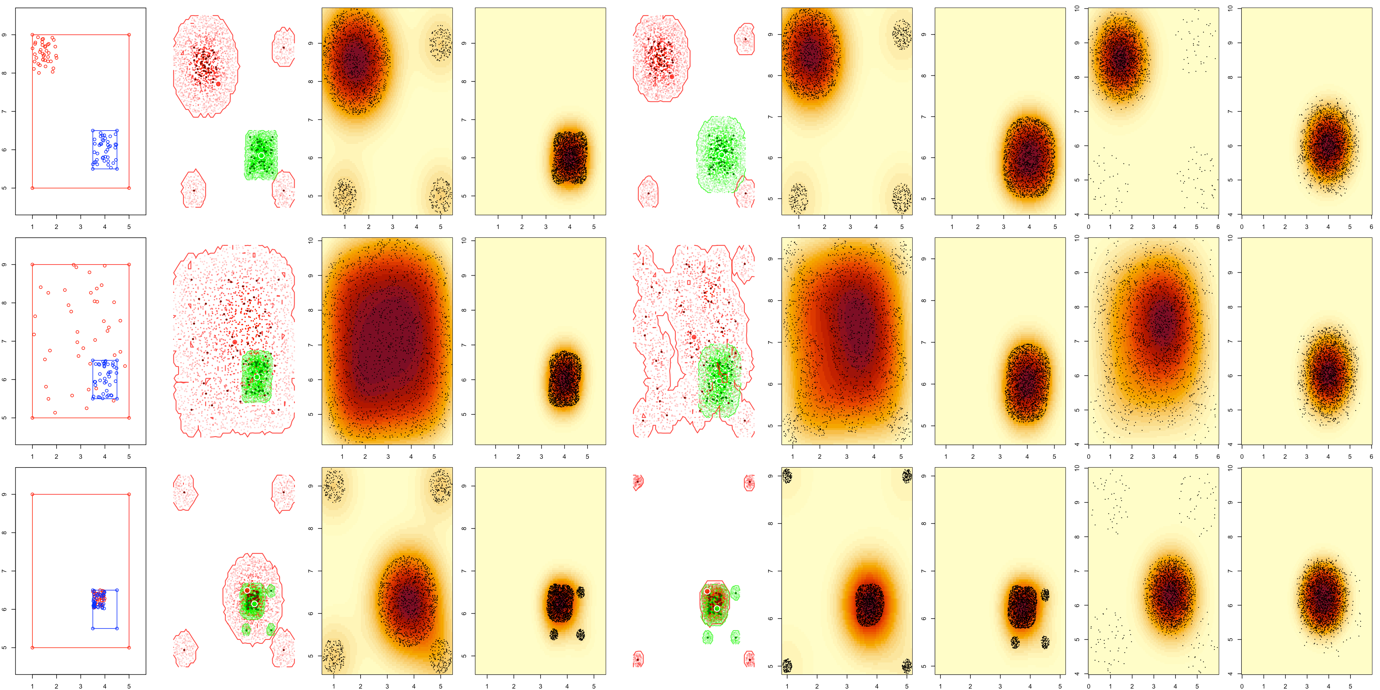

**Figure B10.** A set of theoretical point distribution used to examine the behaviour of the different indices: nested MCPs of different sizes, 40 points. In the first row, the points are located far from each other within the MCPs. In the second row, the points are randomly distributed within the MCPs. In the last row, the points are located close each other within the MCPs. The first column shows the point distributions and MCPs in the functional space. The second column shows the polytopes generated by the KDH approach Version 1. The third and fourth columns show the kernels and a sample of the random points from which they are computed, used in the KDH approach Version 1 and the kernel-based approach Version 1. The fifth column shows the polytopes generated by the KDH approach Version 2. The sixth and seventh columns show the kernels and a sample of the random points from which they are computed, used in the KDH approach Version 2 and the kernel-based approach Version 2. The eighth and ninth columns show the kernels and a sample of the random points from which they are computed, used in the kernel-based approach Version 3.

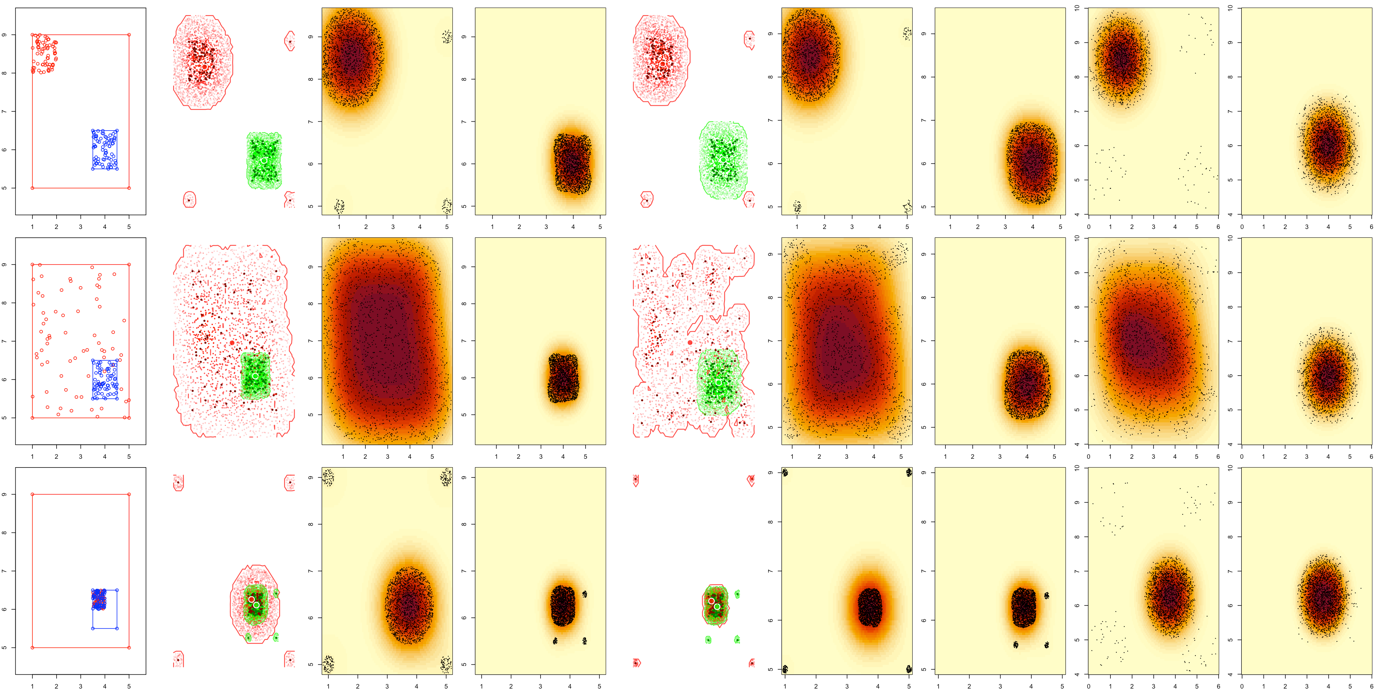

**Figure B11.** A set of theoretical point distribution used to examine the behaviour of the different indices: nested MCPs of different sizes, 70 points. In the first row, the points are located far from each other within the MCPs. In the second row, the points are randomly distributed within the MCPs. In the last row, the points are located close each other within the MCPs. The first column shows the point distributions and MCPs in the functional space. The second column shows the polytopes generated by the KDH approach Version 1. The third and fourth columns show the kernels and a sample of the random points from which they are computed, used in the KDH approach Version 1 and the kernel-based approach Version 1. The fifth column shows the polytopes generated by the KDH approach Version 2. The sixth and seventh columns show the kernels and a sample of the random points from which they are computed, used in the KDH approach Version 2 and the kernel-based approach Version 2. The eighth and ninth columns show the kernels and a sample of the random points from which they are computed, used in the kernel-based approach Version 3.

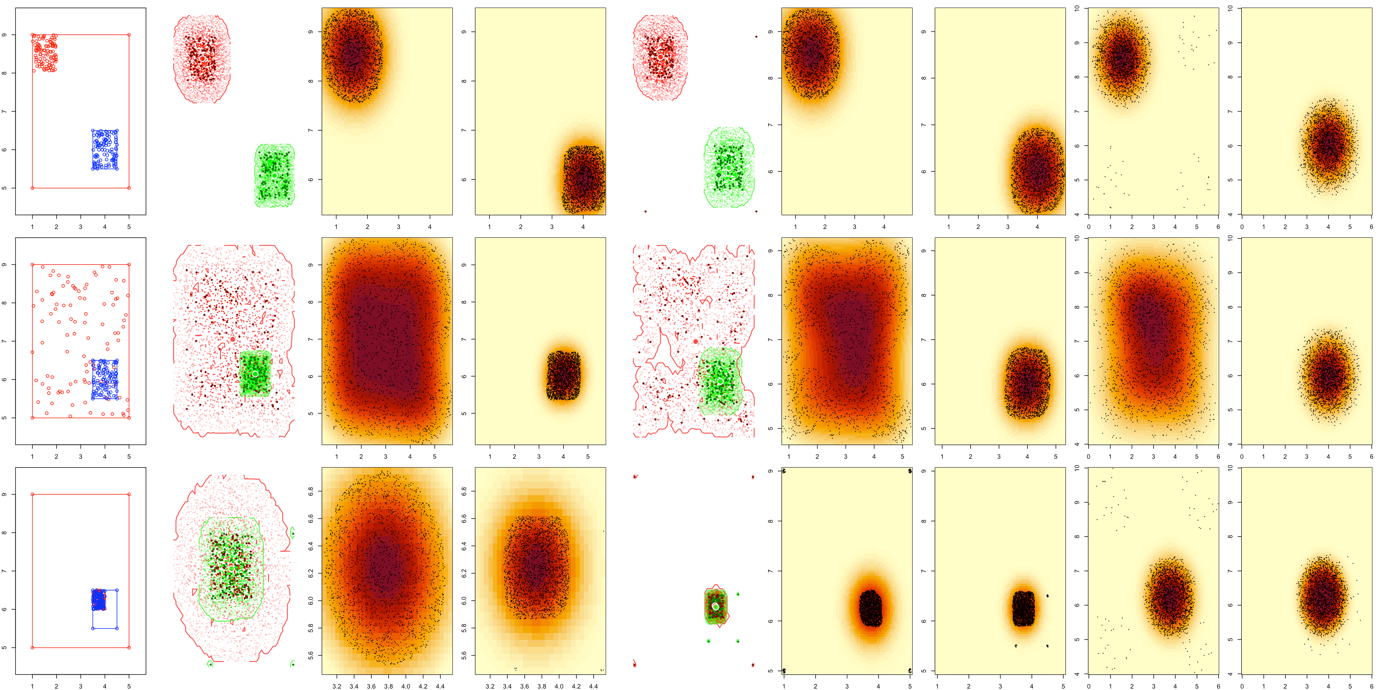

**Figure B12.** A set of theoretical point distribution used to examine the behaviour of the different indices: nested MCPs of different sizes, 100 points. In the first row, the points are located far from each other within the MCPs. In the second row, the points are randomly distributed within the MCPs. In the last row, the points are located close each other within the MCPs. The first column shows the point distributions and MCPs in the functional space. The second column shows the polytopes generated by the KDH approach Version 1. The third and fourth columns show the kernels and a sample of the random points from which they are computed, used in the KDH approach Version 1 and the kernel-based approach Version 1. The fifth column shows the polytopes generated by the KDH approach Version 2. The sixth and seventh columns show the kernels and a sample of the random points from which they are computed, used in the KDH approach Version 2 and the kernel-based approach Version 2. The eighth and ninth columns show the kernels and a sample of the random points from which they are computed, used in the kernel-based approach Version 3.

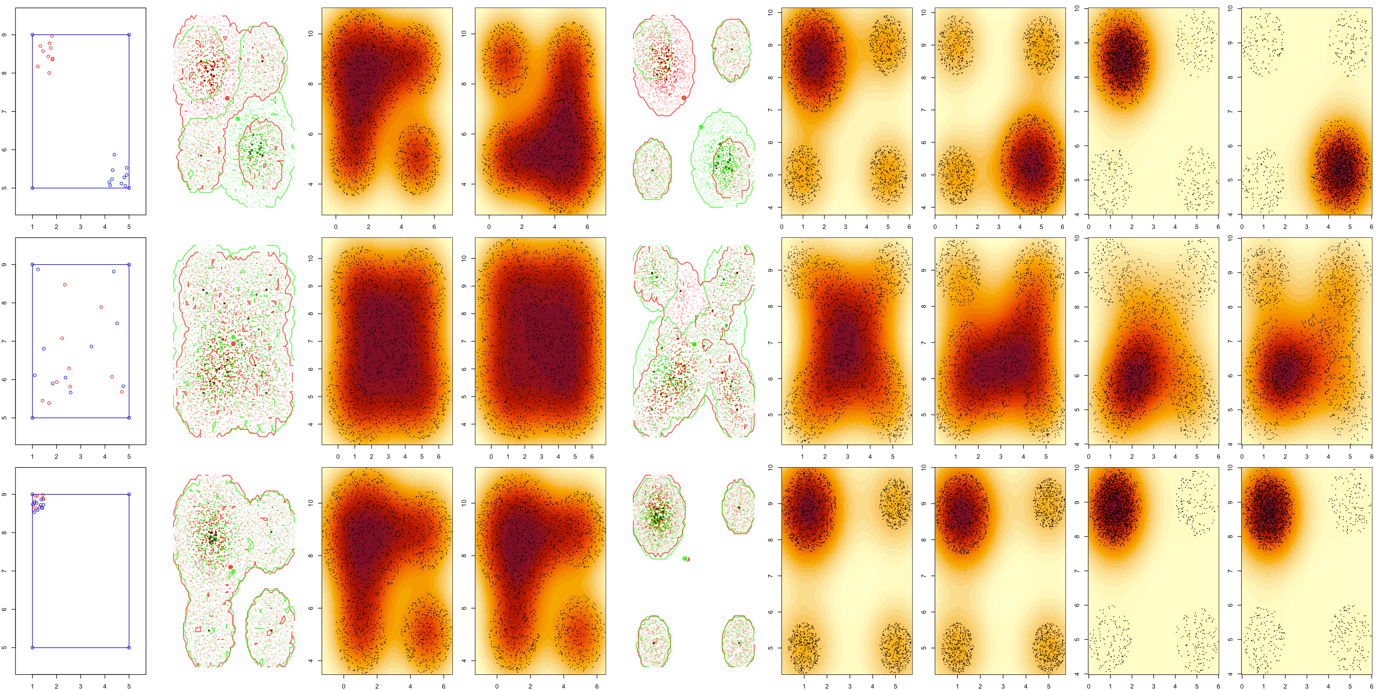

**Figure B13.** A set of theoretical point distribution used to examine the behaviour of the different indices: nested MCPs of same size, 10 points. In the first row, the points are located far from each other within the MCPs. In the second row, the points are randomly distributed within the MCPs. In the last row, the points are located close each other within the MCPs. The first column shows the point distributions and MCPs in the functional space. The second column shows the polytopes generated by the KDH approach Version 1. The third and fourth columns show the kernels and a sample of the random points from which they are computed, used in the KDH approach Version 1 and the kernel-based approach Version 1. The fifth column shows the polytopes generated by the KDH approach Version 2. The sixth and seventh columns show the kernels and a sample of the random points from which they are computed, used in the KDH approach Version 2 and the kernel-based approach Version 2. The eighth and ninth columns show the kernels and a sample of the random points from which they are computed, used in the kernel-based approach Version 3.

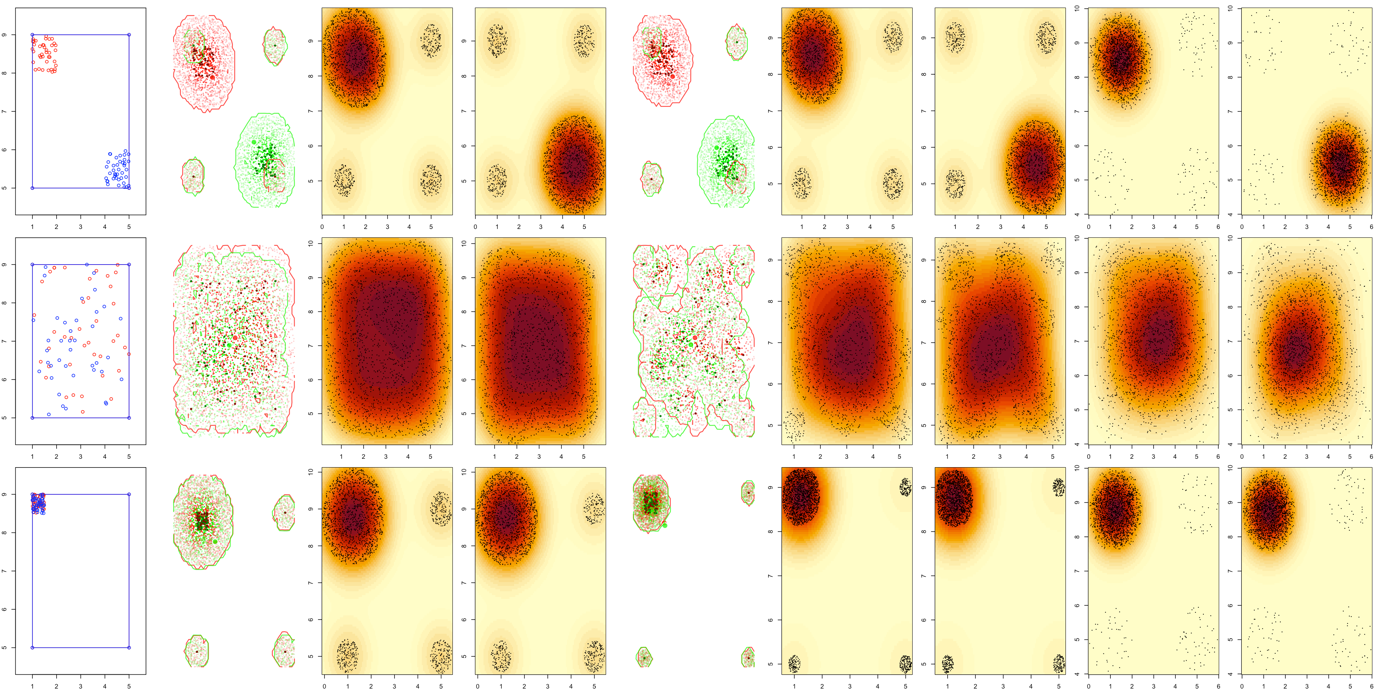

**Figure B14.** A set of theoretical point distribution used to examine the behaviour of the different indices: nested MCPs of same size, 40 points. In the first row, the points are located far from each other within the MCPs. In the second row, the points are randomly distributed within the MCPs. In the last row, the points are located close each other within the MCPs. The first column shows the point distributions and MCPs in the functional space. The second column shows the polytopes generated by the KDH approach Version 1. The third and fourth columns show the kernels and a sample of the random points from which they are computed, used in the KDH approach Version 1 and the kernel-based approach Version 1. The fifth column shows the polytopes generated by the KDH approach Version 2. The sixth and seventh columns show the kernels and a sample of the random points from which they are computed, used in the KDH approach Version 2 and the kernel-based approach Version 2. The eighth and ninth columns show the kernels and a sample of the random points from which they are computed, used in the kernel-based approach Version 3.

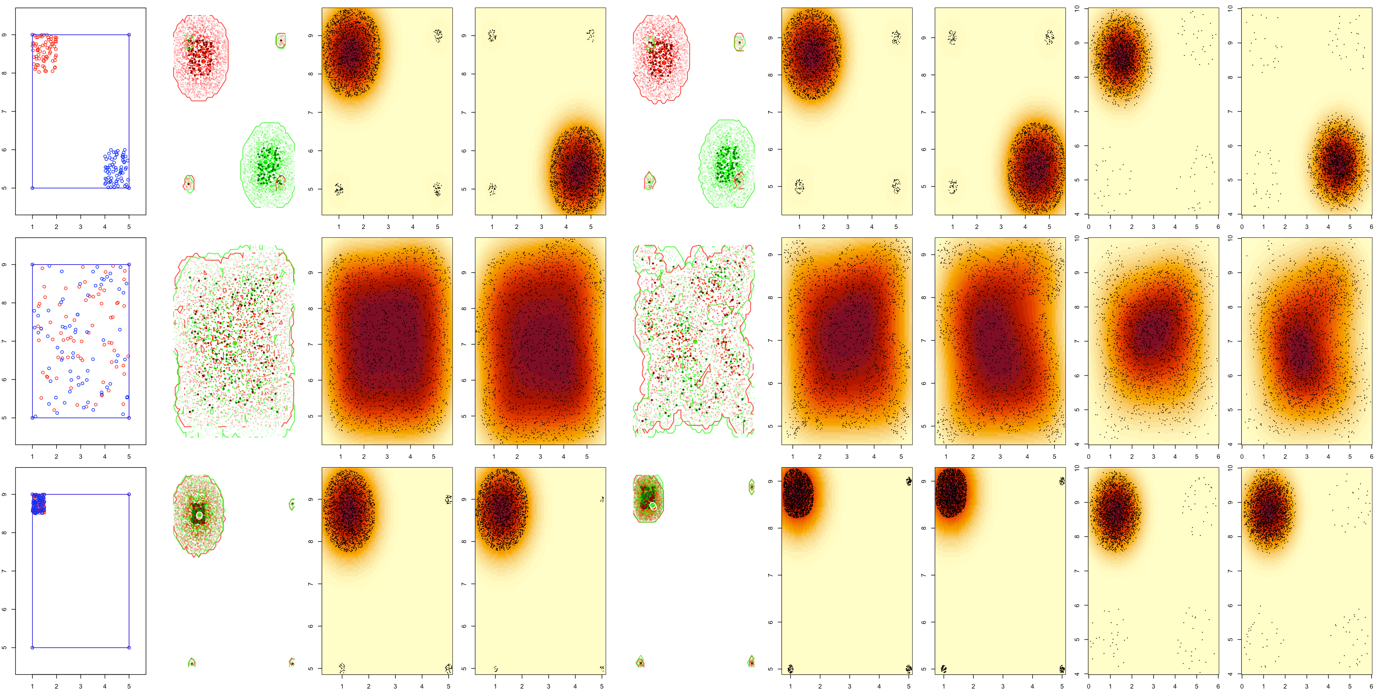

**Figure B15.** A set of theoretical point distribution used to examine the behaviour of the different indices: nested MCPs of same size, 70 points. In the first row, the points are located far from each other within the MCPs. In the second row, the points are randomly distributed within the MCPs. In the last row, the points are located close each other within the MCPs. The first column shows the point distributions and MCPs in the functional space. The second column shows the polytopes generated by the KDH approach Version 1. The third and fourth columns show the kernels and a sample of the random points from which they are computed, used in the KDH approach Version 1 and the kernel-based approach Version 1. The fifth column shows the polytopes generated by the KDH approach Version 2. The sixth and seventh columns show the kernels and a sample of the random points from which they are computed, used in the KDH approach Version 2 and the kernel-based approach Version 2. The eighth and ninth columns show the kernels and a sample of the random points from which they are computed, used in the kernel-based approach Version 3.

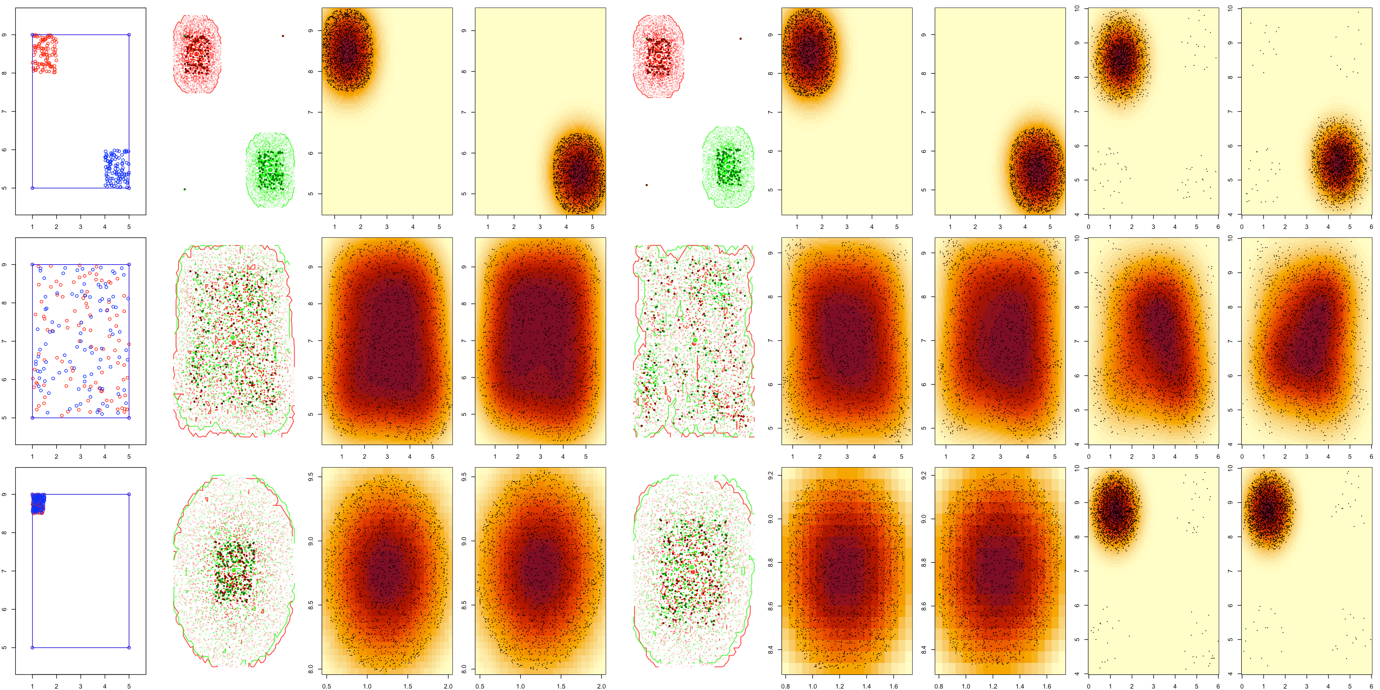

**Figure B16.** A set of theoretical point distribution used to examine the behaviour of the different indices: nested MCPs of same size, 100 points. In the first row, the points are located far from each other within the MCPs. In the second row, the points are randomly distributed within the MCPs. In the last row, the points are located close each other within the MCPs. The first column shows the point distributions and MCPs in the functional space. The second column shows the polytopes generated by the KDH approach Version 1. The third and fourth columns show the kernels and a sample of the random points from which they are computed, used in the KDH approach Version 1 and the kernel-based approach Version 1. The fifth column shows the polytopes generated by the KDH approach Version 2. The sixth and seventh columns show the kernels and a sample of the random points from which they are computed, used in the KDH approach Version 2 and the kernel-based approach Version 2. The eighth and ninth columns show the kernels and a sample of the random points from which they are computed, used in the kernel-based approach Version 3.

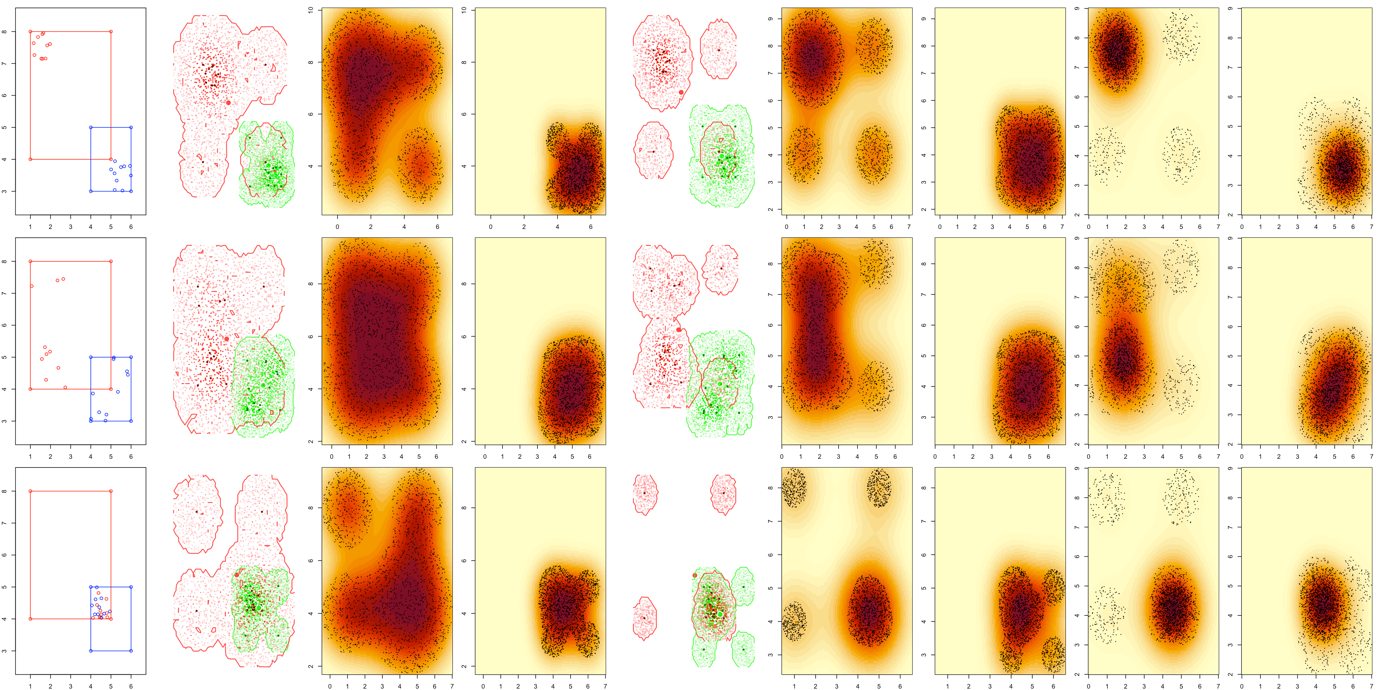

**Figure B17.** A set of theoretical point distribution used to examine the behaviour of the different indices: overlapping MCPs of different sizes, 10 points. In the first row, the points are located far from each other within the MCPs. In the second row, the points are randomly distributed within the MCPs. In the last row, the points are located close each other within the MCPs. The first column shows the point distributions and MCPs in the functional space. The second column shows the polytopes generated by the KDH approach Version 1. The third and fourth columns show the kernels and a sample of the random points from which they are computed, used in the KDH approach Version 1 and the kernel-based approach Version 1. The fifth column shows the polytopes generated by the KDH approach Version 2. The sixth and seventh columns show the kernels and a sample of the random points from which they are computed, used in the KDH approach Version 2 and the kernel-based approach Version 2. The eighth and ninth columns show the kernels and a sample of the random points from which they are computed, used in the kernel-based approach Version 3.

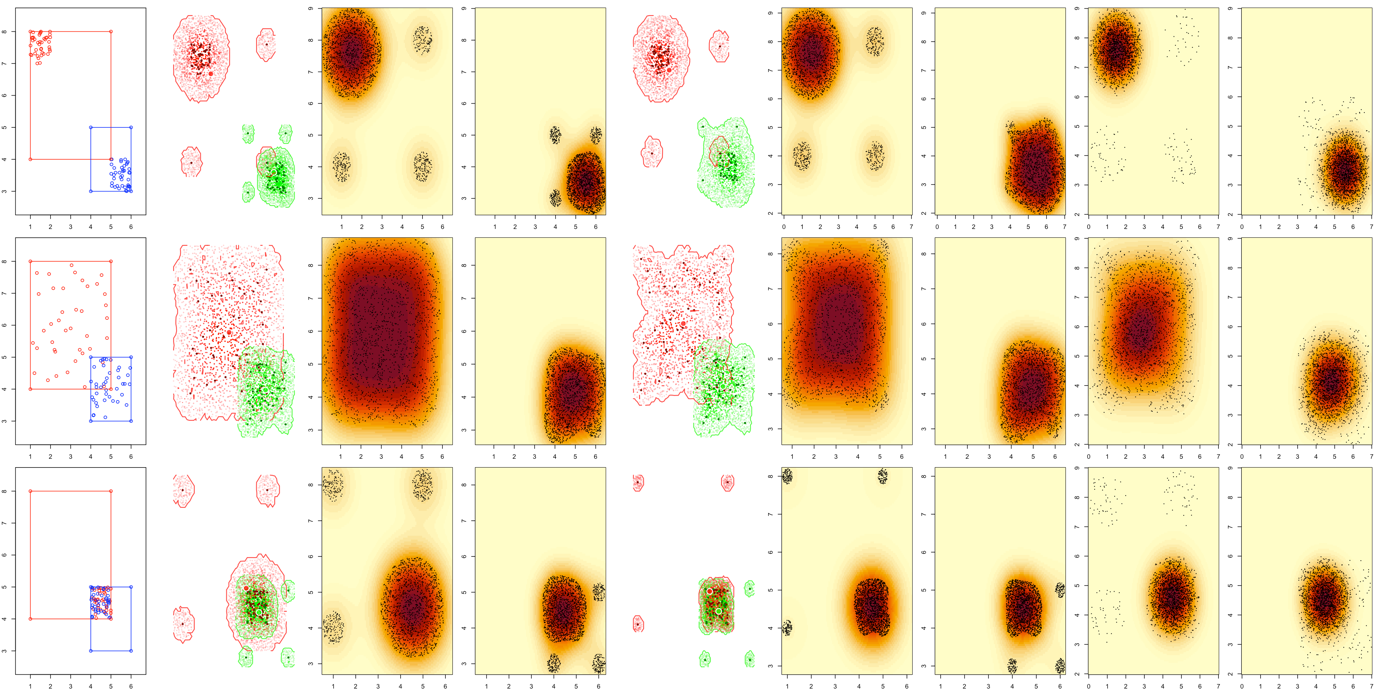

**Figure B18.** A set of theoretical point distribution used to examine the behaviour of the different indices: overlapping MCPs of different sizes, 40 points. In the first row, the points are located far from each other within the MCPs. In the second row, the points are randomly distributed within the MCPs. In the last row, the points are located close each other within the MCPs. The first column shows the point distributions and MCPs in the functional space. The second column shows the polytopes generated by the KDH approach Version 1. The third and fourth columns show the kernels and a sample of the random points from which they are computed, used in the KDH approach Version 1 and the kernel-based approach Version 1. The fifth column shows the polytopes generated by the KDH approach Version 2. The sixth and seventh columns show the kernels and a sample of the random points from which they are computed, used in the KDH approach Version 2 and the kernel-based approach Version 2. The eighth and ninth columns show the kernels and a sample of the random points from which they are computed, used in the kernel-based approach Version 3.

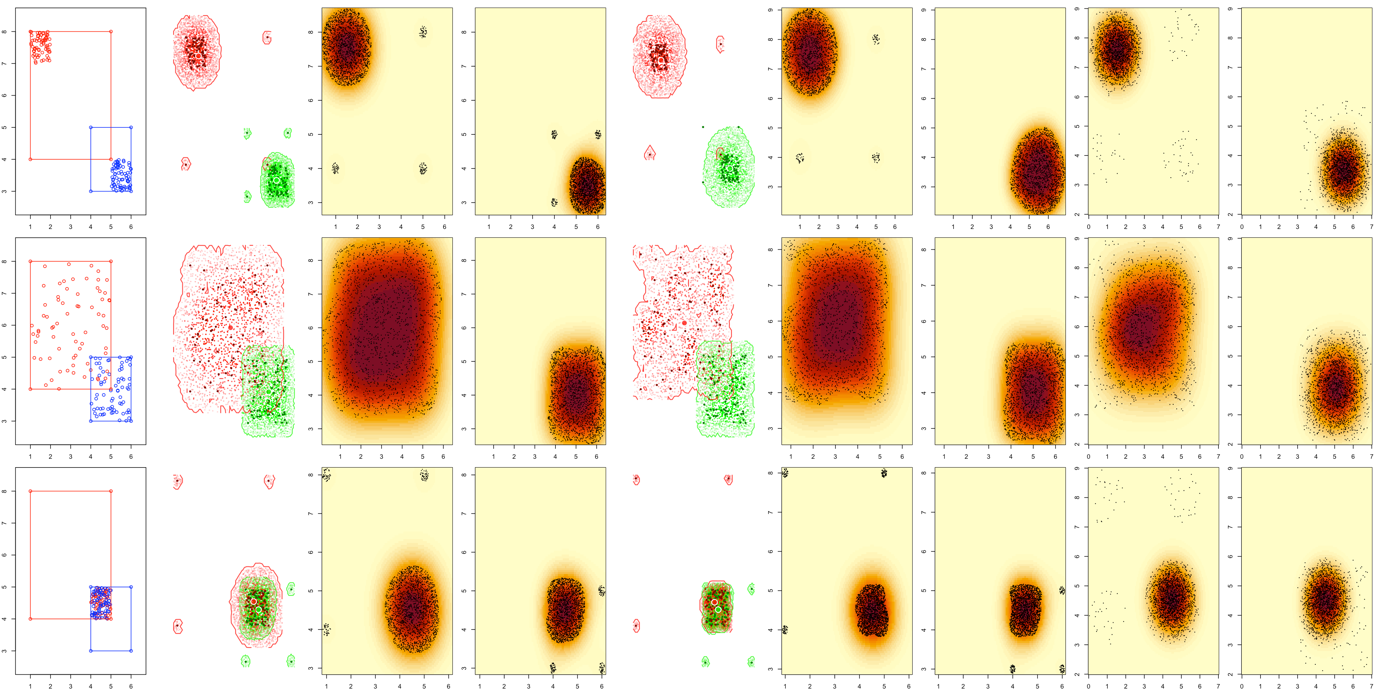

**Figure B19.** A set of theoretical point distribution used to examine the behaviour of the different indices: overlapping MCPs of different sizes, 70 points. In the first row, the points are located far from each other within the MCPs. In the second row, the points are randomly distributed within the MCPs. In the last row, the points are located close each other within the MCPs. The first column shows the point distributions and MCPs in the functional space. The second column shows the polytopes generated by the KDH approach Version 1. The third and fourth columns show the kernels and a sample of the random points from which they are computed, used in the KDH approach Version 1 and the kernel-based approach Version 1. The fifth column shows the polytopes generated by the KDH approach Version 2. The sixth and seventh columns show the kernels and a sample of the random points from which they are computed, used in the KDH approach Version 2 and the kernel-based approach Version 2. The eighth and ninth columns show the kernels and a sample of the random points from which they are computed, used in the kernel-based approach Version 3.

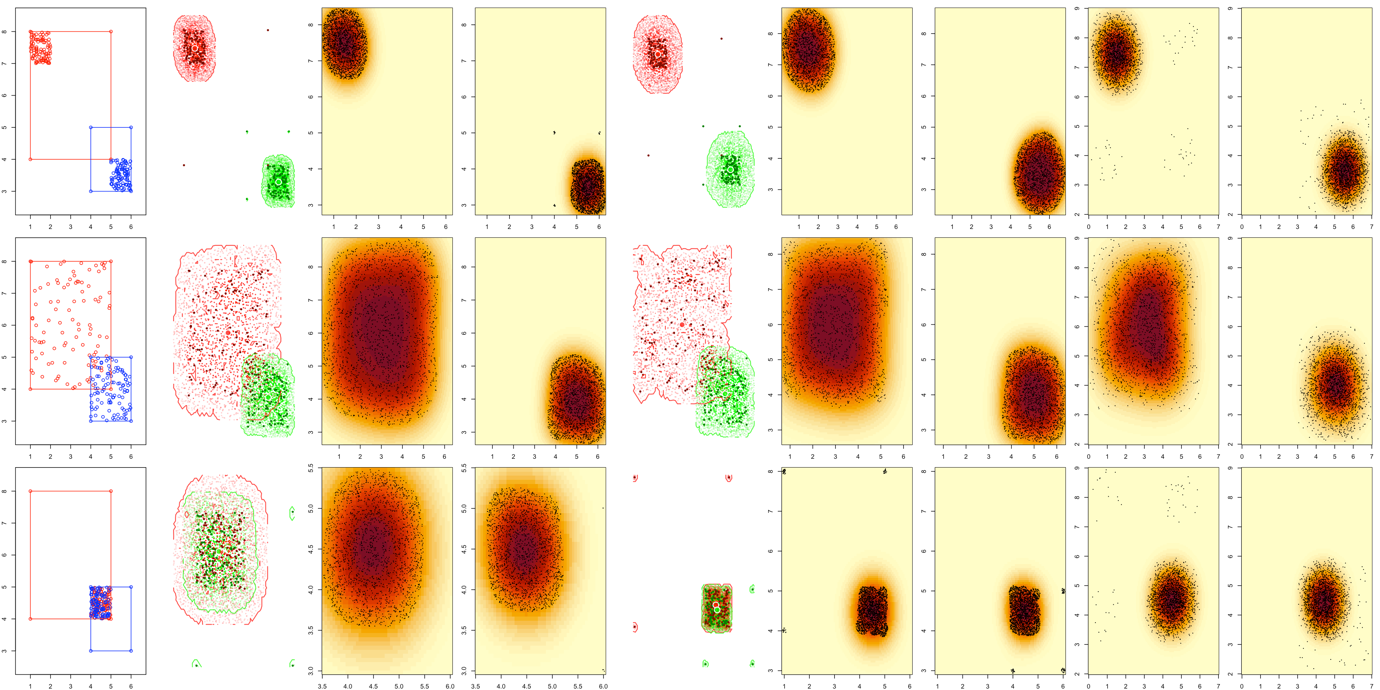

**Figure B20.** A set of theoretical point distribution used to examine the behaviour of the different indices: overlapping MCPs of different sizes, 100 points. In the first row, the points are located far from each other within the MCPs. In the second row, the points are randomly distributed within the MCPs. In the last row, the points are located close each other within the MCPs. The first column shows the point distributions and MCPs in the functional space. The second column shows the polytopes generated by the KDH approach Version 1. The third and fourth columns show the kernels and a sample of the random points from which they are computed, used in the KDH approach Version 1 and the kernel-based approach Version 1. The fifth column shows the polytopes generated by the KDH approach Version 2. The sixth and seventh columns show the kernels and a sample of the random points from which they are computed, used in the KDH approach Version 2 and the kernel-based approach Version 2. The eighth and ninth columns show the kernels and a sample of the random points from which they are computed, used in the kernel-based approach Version 3.

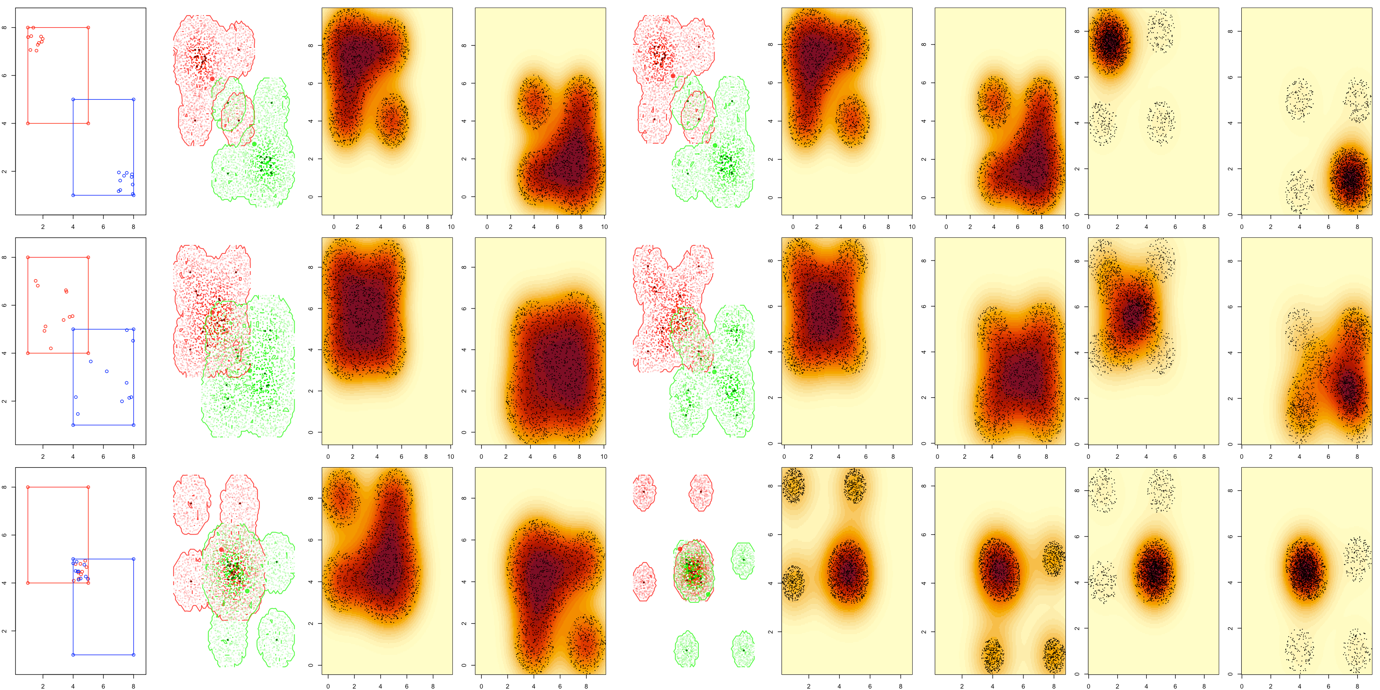

**Figure B21.** A set of theoretical point distribution used to examine the behaviour of the different indices: overlapping MCPs of same size, 10 points. In the first row, the points are located far from each other within the MCPs. In the second row, the points are randomly distributed within the MCPs. In the last row, the points are located close each other within the MCPs. The first column shows the point distributions and MCPs in the functional space. The second column shows the polytopes generated by the KDH approach Version 1. The third and fourth columns show the kernels and a sample of the random points from which they are computed, used in the KDH approach Version 1 and the kernel-based approach Version 1. The fifth column shows the polytopes generated by the KDH approach Version 2. The sixth and seventh columns show the kernels and a sample of the random points from which they are computed, used in the KDH approach Version 2 and the kernel-based approach Version 2. The eighth and ninth columns show the kernels and a sample of the random points from which they are computed, used in the kernel-based approach Version 3.

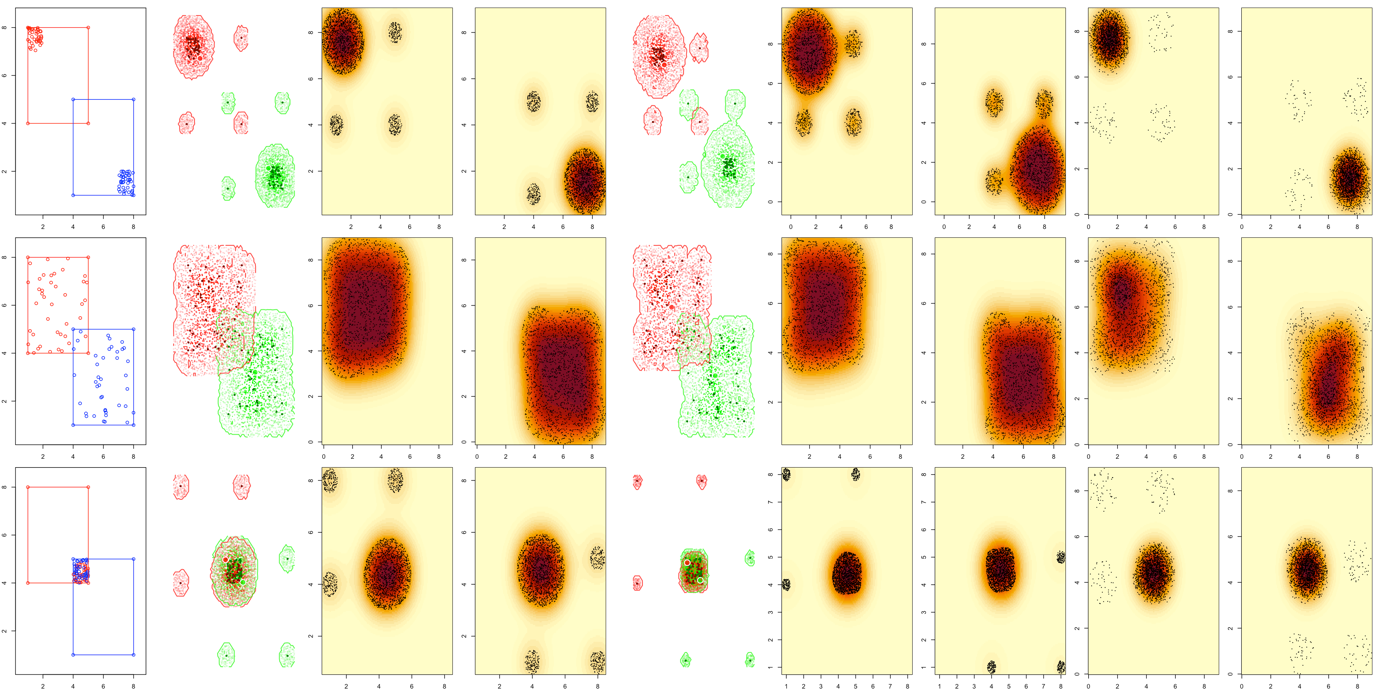

**Figure B22.** A set of theoretical point distribution used to examine the behaviour of the different indices: overlapping MCPs of same size, 40 points. In the first row, the points are located far from each other within the MCPs. In the second row, the points are randomly distributed within the MCPs. In the last row, the points are located close each other within the MCPs. The first column shows the point distributions and MCPs in the functional space. The second column shows the polytopes generated by the KDH approach Version 1. The third and fourth columns show the kernels and a sample of the random points from which they are computed, used in the KDH approach Version 1 and the kernel-based approach Version 1. The fifth column shows the polytopes generated by the KDH approach Version 2. The sixth and seventh columns show the kernels and a sample of the random points from which they are computed, used in the KDH approach Version 2 and the kernel-based approach Version 2. The eighth and ninth columns show the kernels and a sample of the random points from which they are computed, used in the kernel-based approach Version 3.

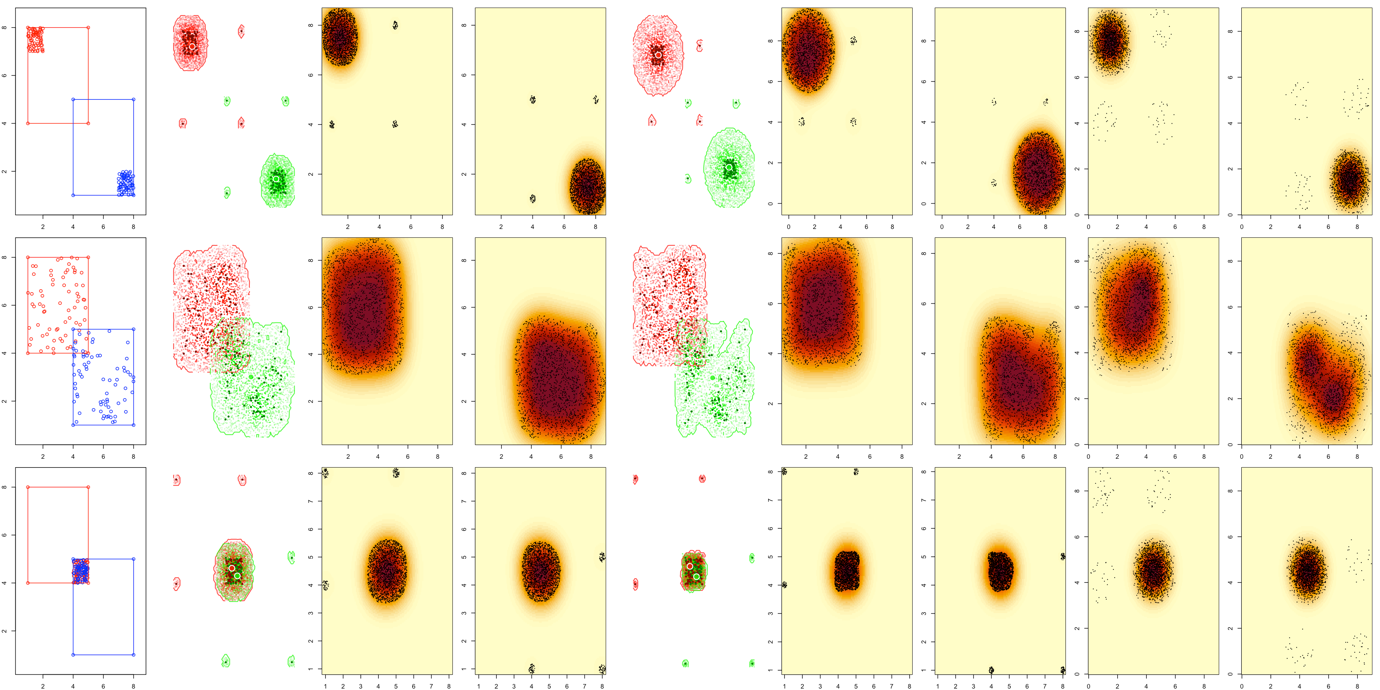

**Figure B23.** A set of theoretical point distribution used to examine the behaviour of the different indices: overlapping MCPs of same size, 70 points. In the first row, the points are located far from each other within the MCPs. In the second row, the points are randomly distributed within the MCPs. In the last row, the points are located close each other within the MCPs. The first column shows the point distributions and MCPs in the functional space. The second column shows the polytopes generated by the KDH approach Version 1. The third and fourth columns show the kernels and a sample of the random points from which they are computed, used in the KDH approach Version 1 and the kernel-based approach Version 1. The fifth column shows the polytopes generated by the KDH approach Version 2. The sixth and seventh columns show the kernels and a sample of the random points from which they are computed, used in the KDH approach Version 2 and the kernel-based approach Version 2. The eighth and ninth columns show the kernels and a sample of the random points from which they are computed, used in the kernel-based approach Version 3.

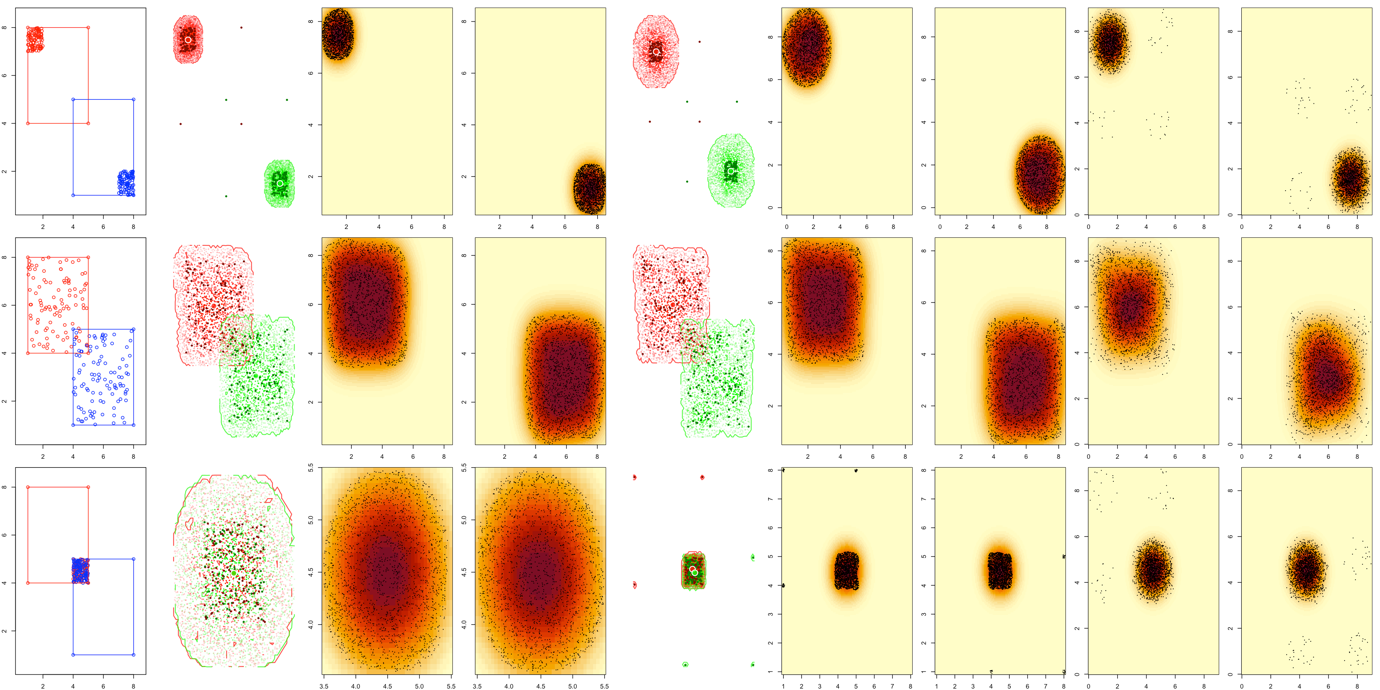

**Figure B24.** A set of theoretical point distribution used to examine the behaviour of the different indices: overlapping MCPs of same size, 100 points. In the first row, the points are located far from each other within the MCPs. In the second row, the points are randomly distributed within the MCPs. In the last row, the points are located close each other within the MCPs. The first column shows the point distributions and MCPs in the functional space. The second column shows the polytopes generated by the KDH approach Version 1. The third and fourth columns show the kernels and a sample of the random points from which they are computed, used in the KDH approach Version 1 and the kernel-based approach Version 1. The fifth column shows the polytopes generated by the KDH approach Version 2. The sixth and seventh columns show the kernels and a sample of the random points from which they are computed, used in the KDH approach Version 2 and the kernel-based approach Version 2. The eighth and ninth columns show the kernels and a sample of the random points from which they are computed, used in the kernel-based approach Version 3.

**Appendix C.** Supplementary results for the theoretical simulations

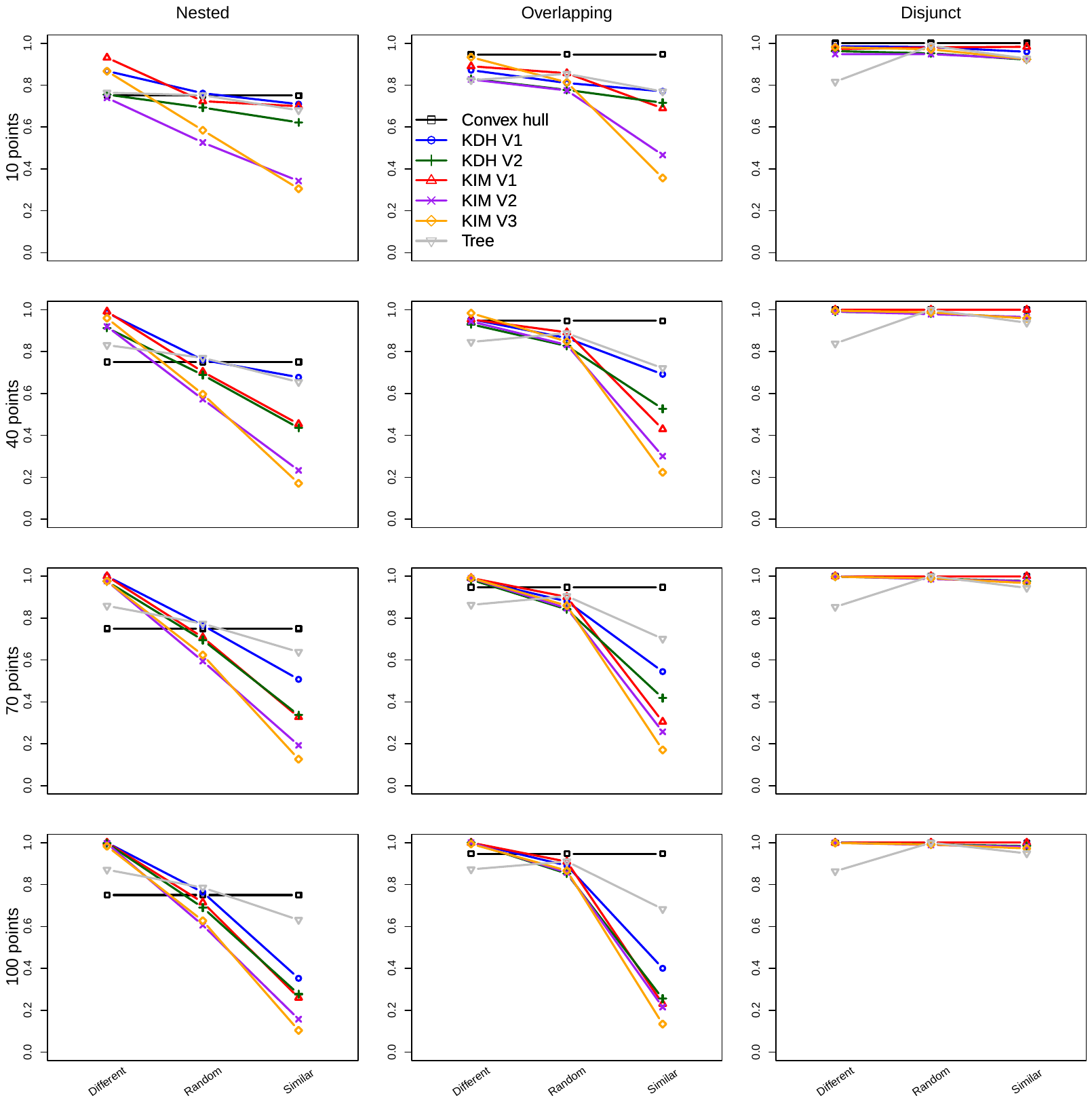

**Figure C1.** Differences in Jaccard dissimilarity between the seven approaches summarised in Table 1, for MCPs of different sizes.

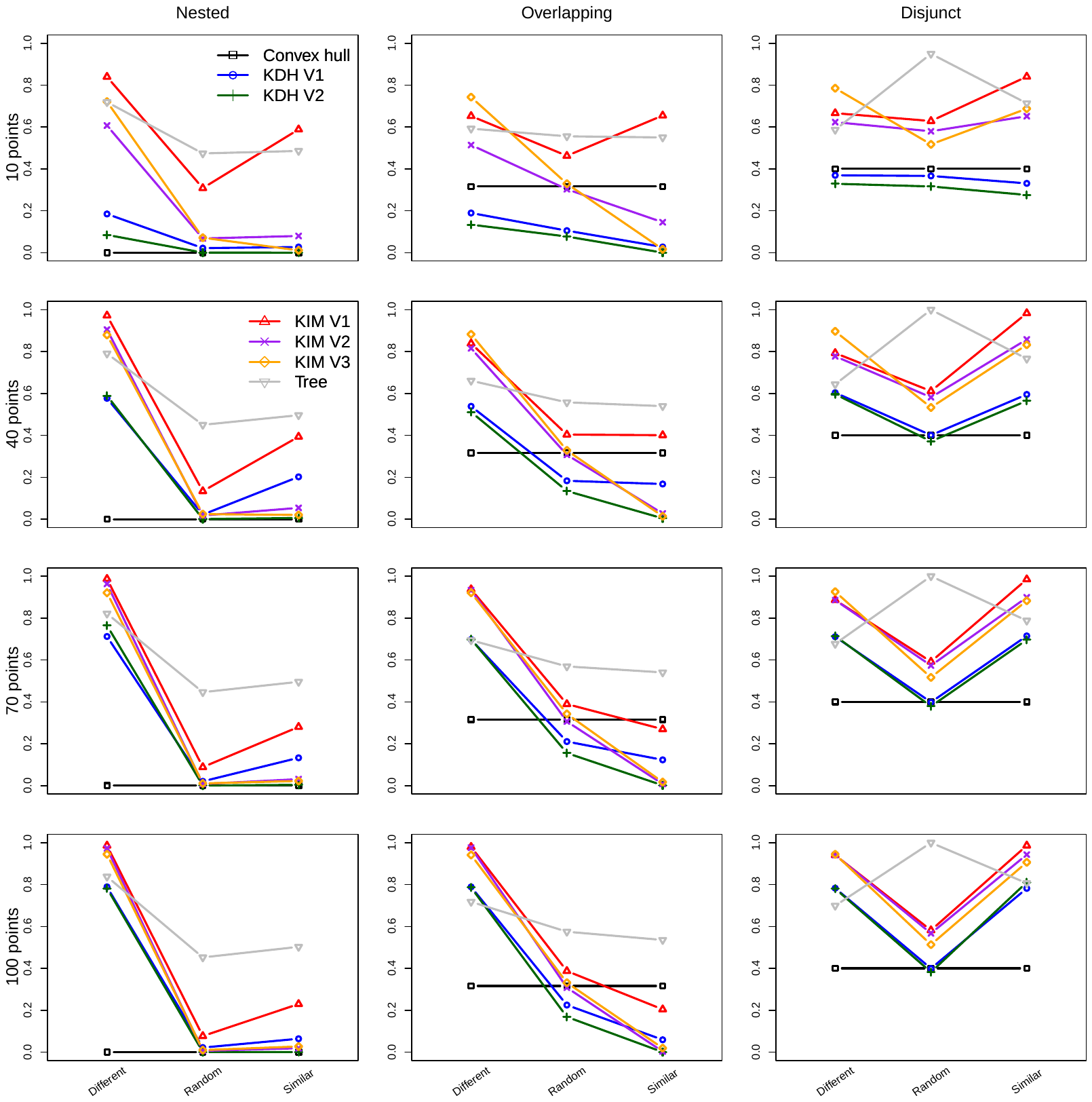

**Figure C2.** Changes in Williams replacement index for the seven approaches summarised in Table 1, for MCPs of different sizes.

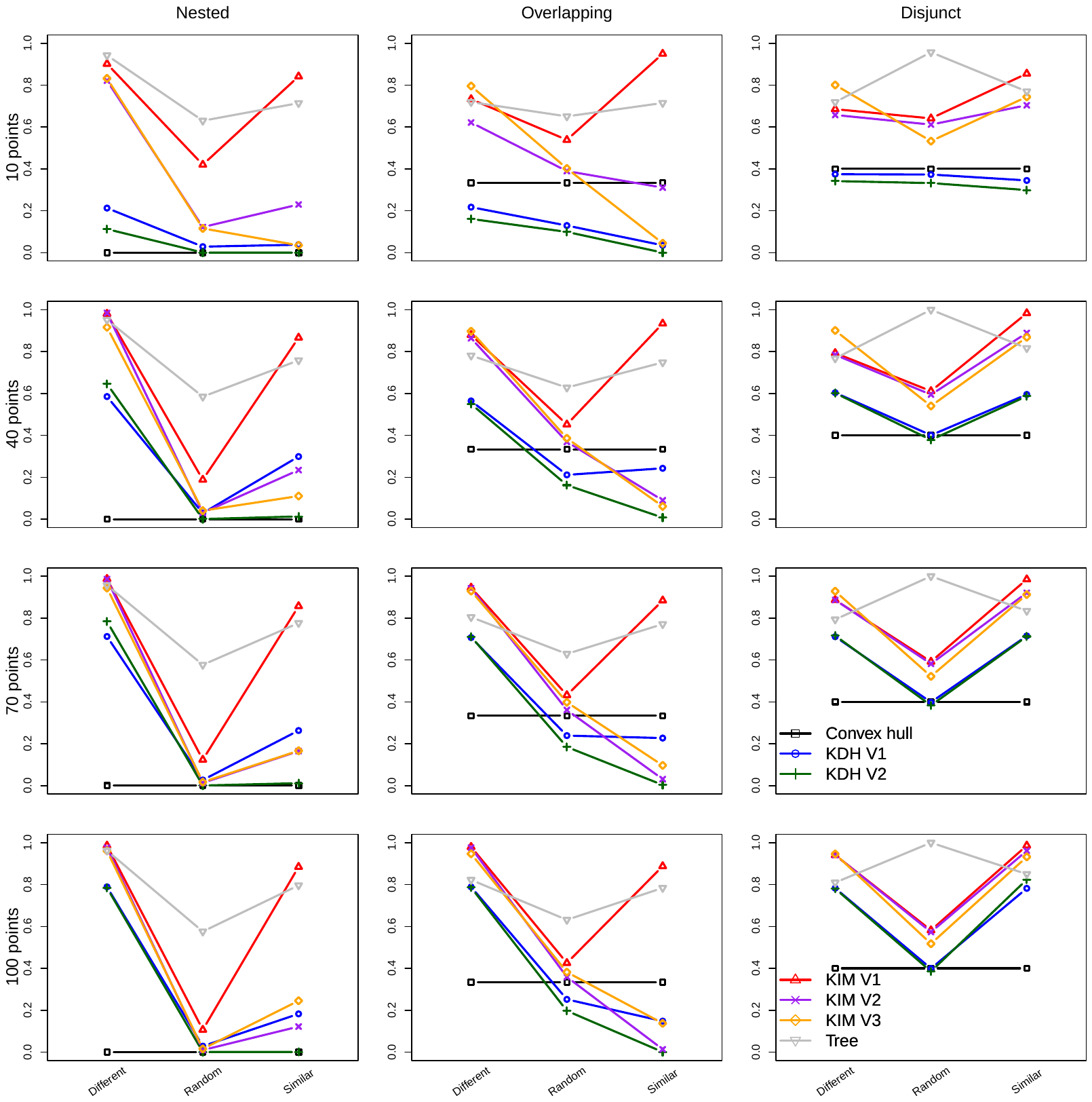

**Figure C3.** Changes in the contribution of replacement to overall turnover, computed as Williams replacement divided by Jaccard dissimilarity, for the seven approaches summarised in Table 1, for MCPs of different sizes.

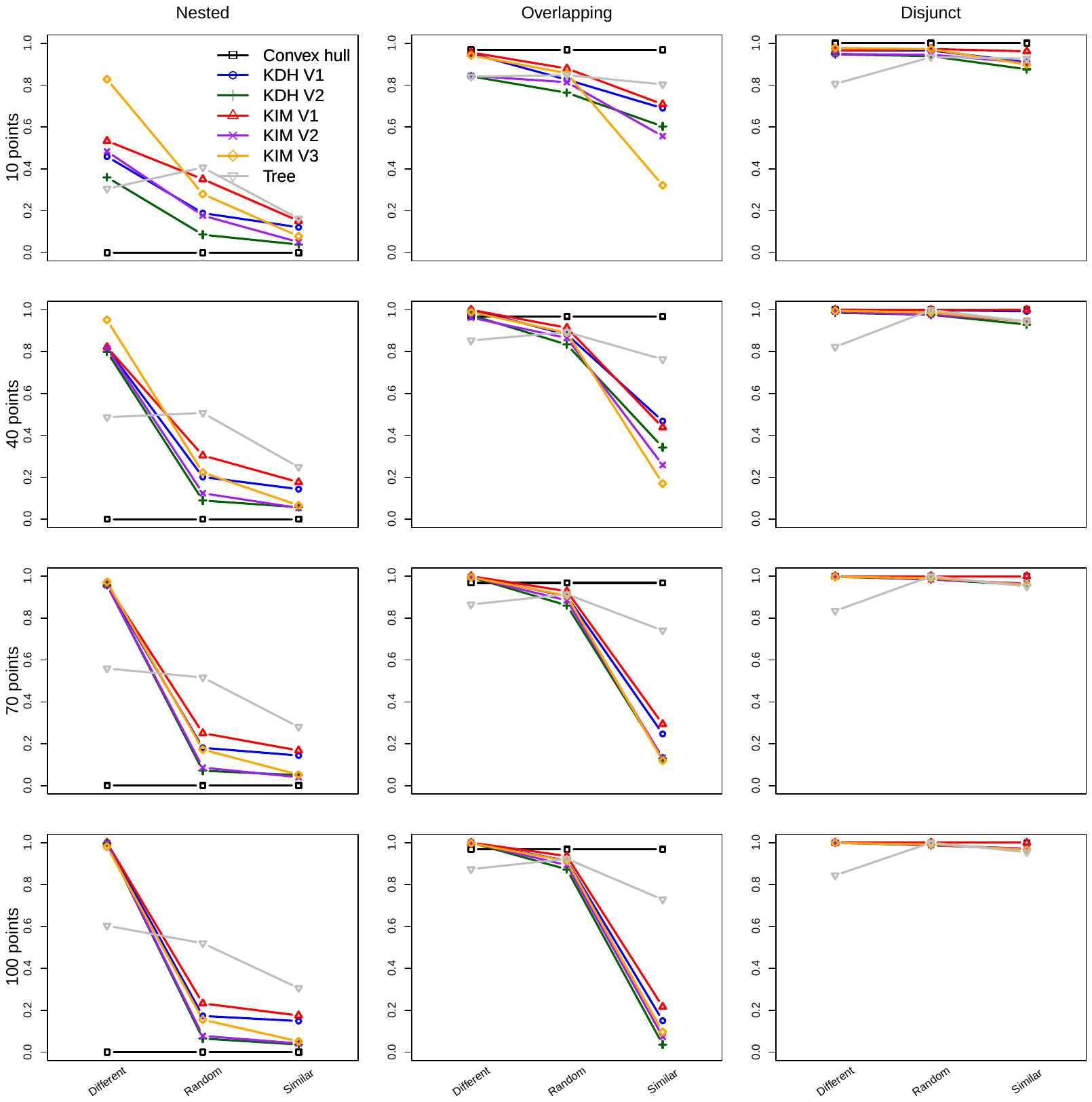

**C4.** Changes in Jaccard dissimilarity for the seven approaches summarised in Table 1, for MCPs of the same size.

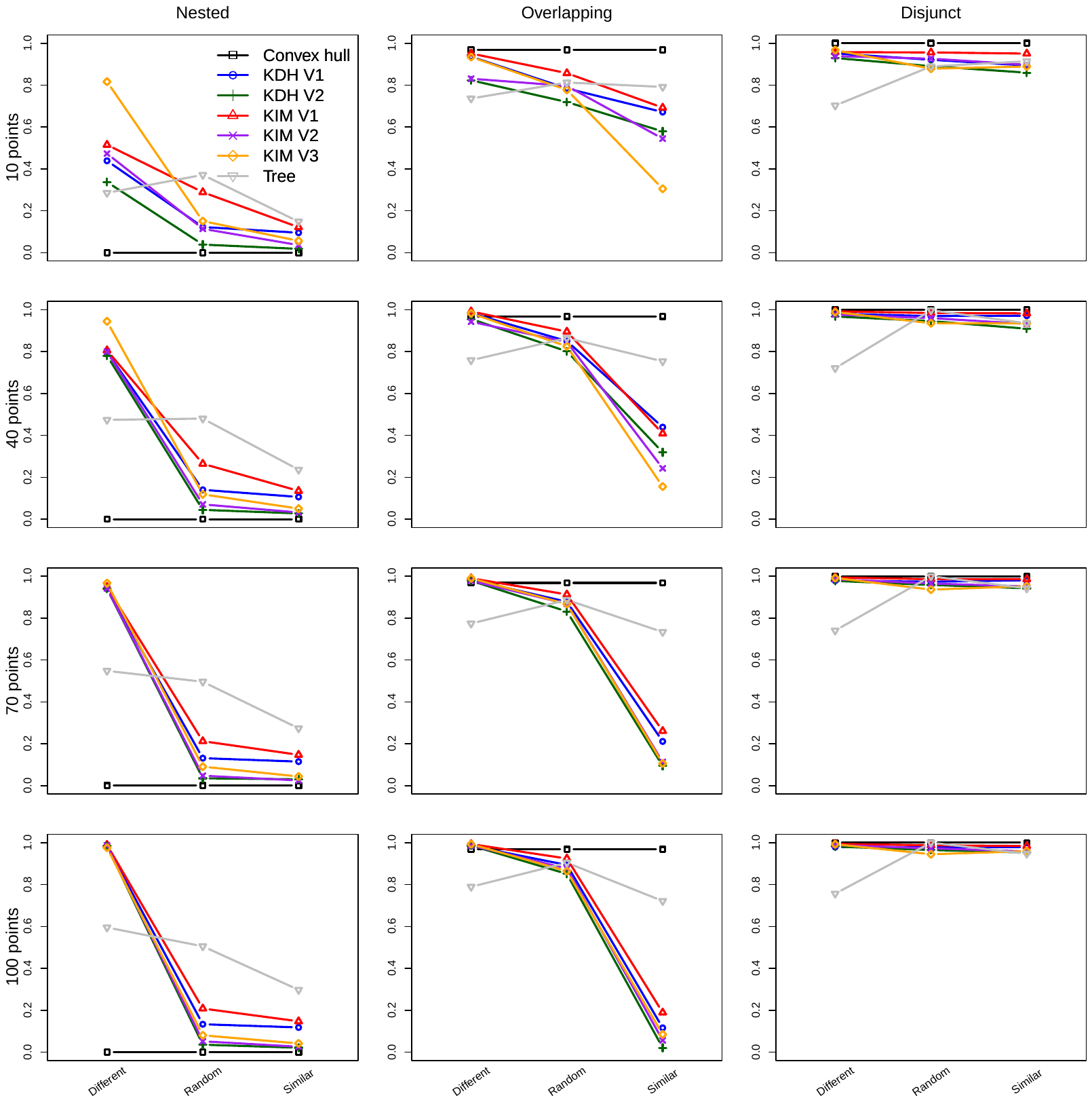

**C5.** Changes in Williams replacement index for the seven approaches summarised in Table 1, for MCPs of the same size.

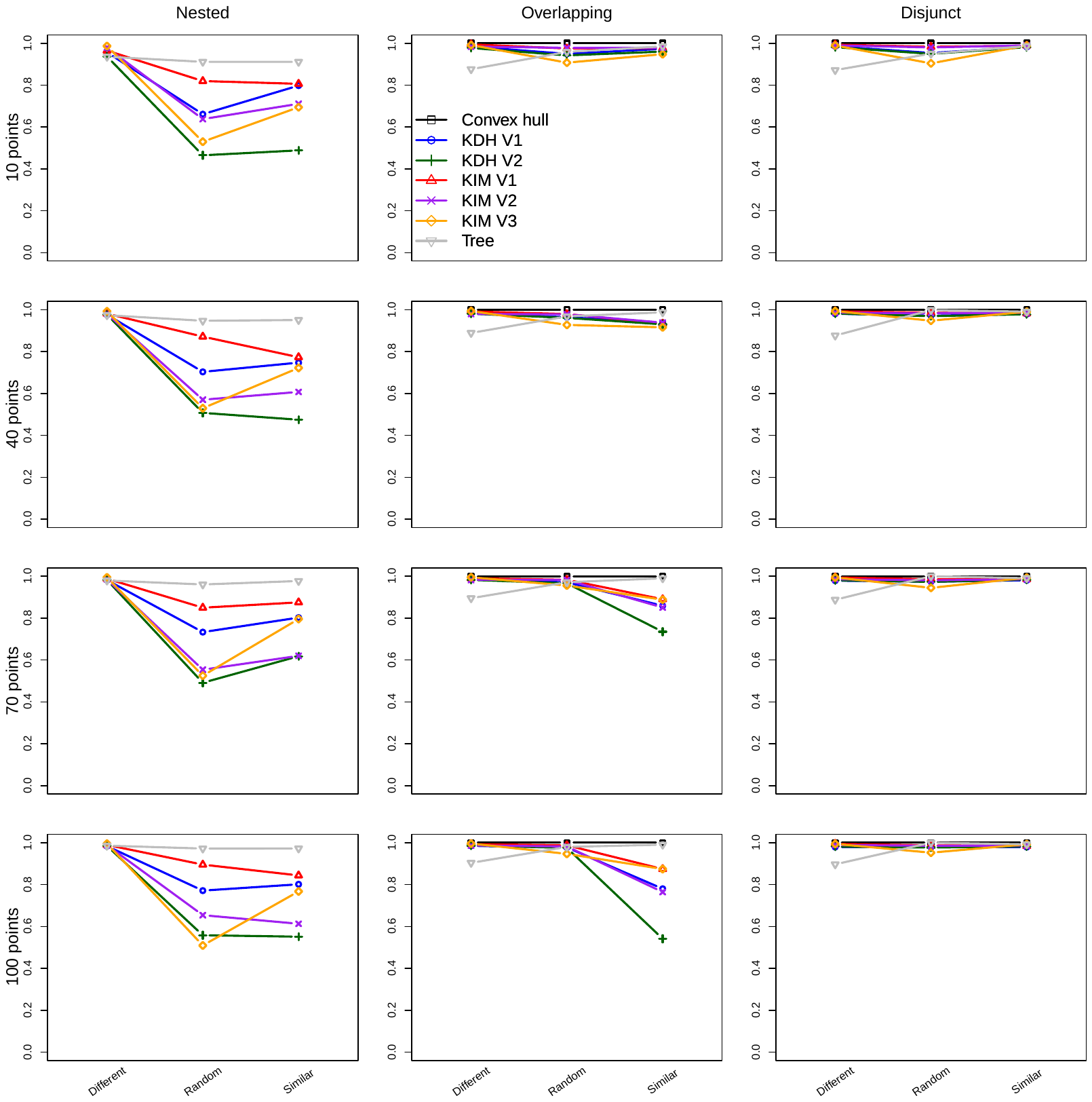

**C6.** Changes in the contribution of replacement to overall turnover, computed as Williams replacement divided by Jaccard dissimilarity, for the seven approaches summarised in Table 1, for MCPs of the same size.

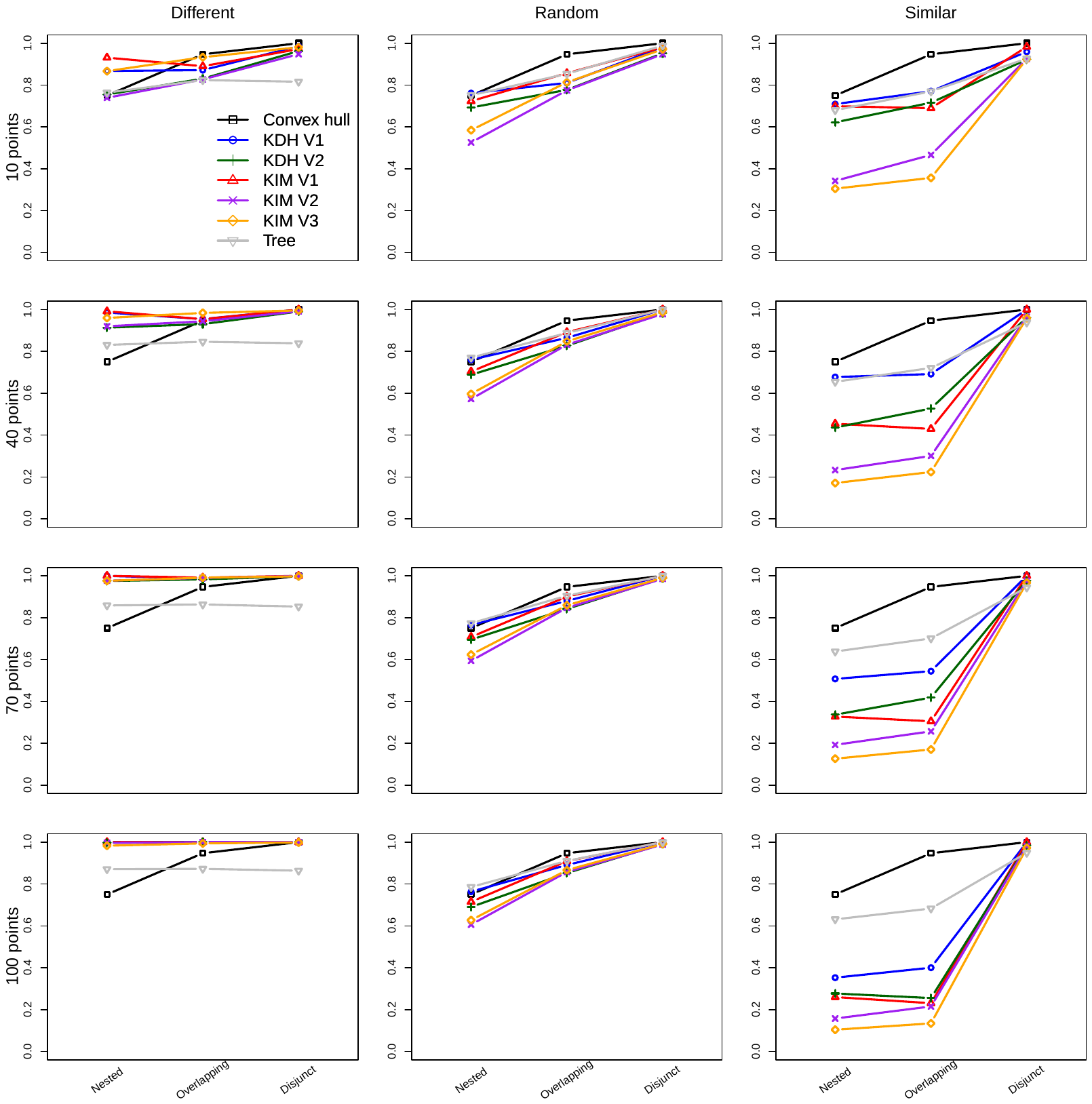

**Figure C7.** Changes in Jaccard dissimilarity for the seven approaches summarised in Table 1, for MCPs of different sizes. These are the same results as Figure C1, ordered differently.

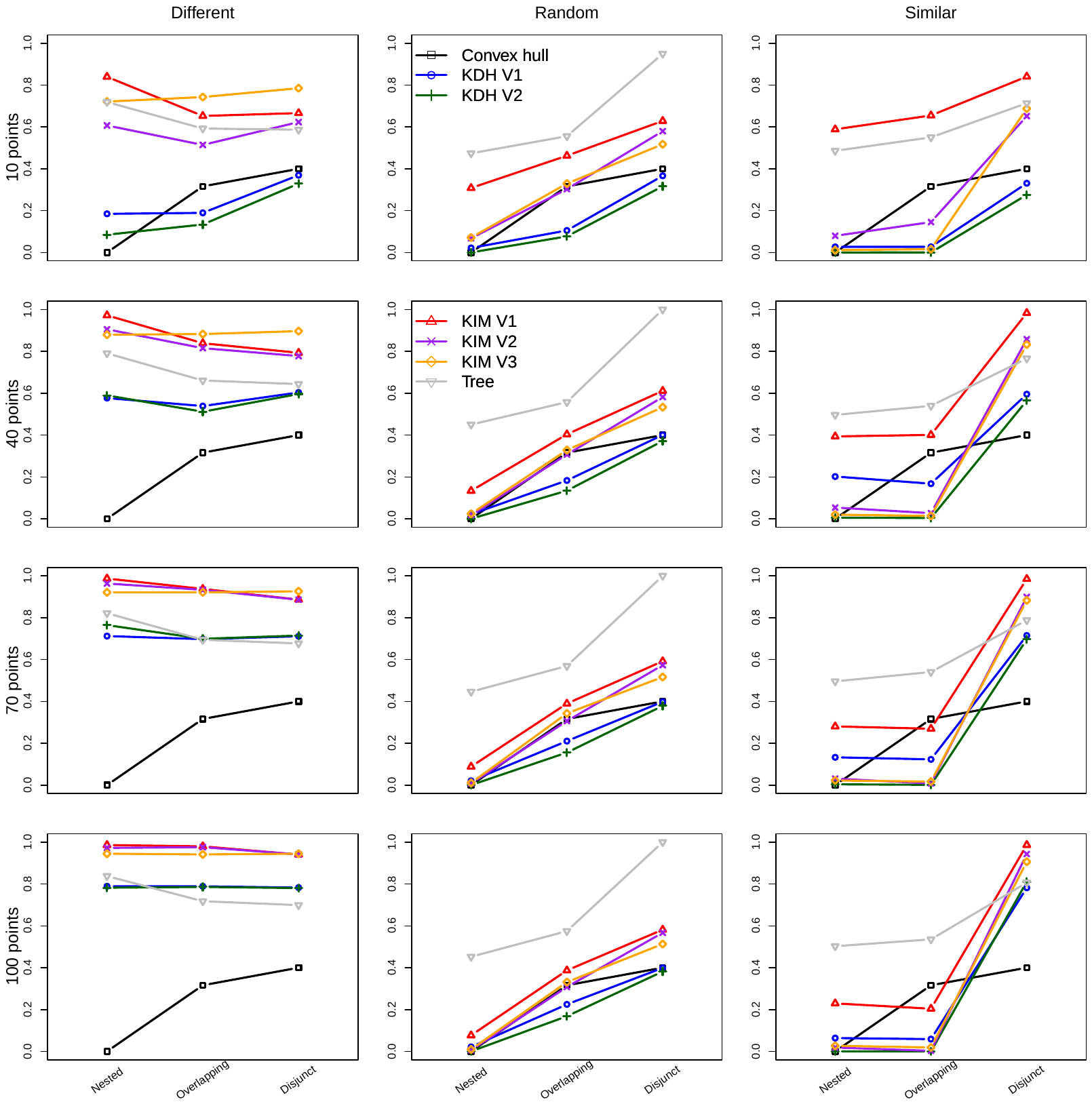

**Figure C8.** Changes in Williams replacement index for the seven approaches summarised in Table 1, for MCPs of different sizes. These are the same results as Figure C2, ordered differently.

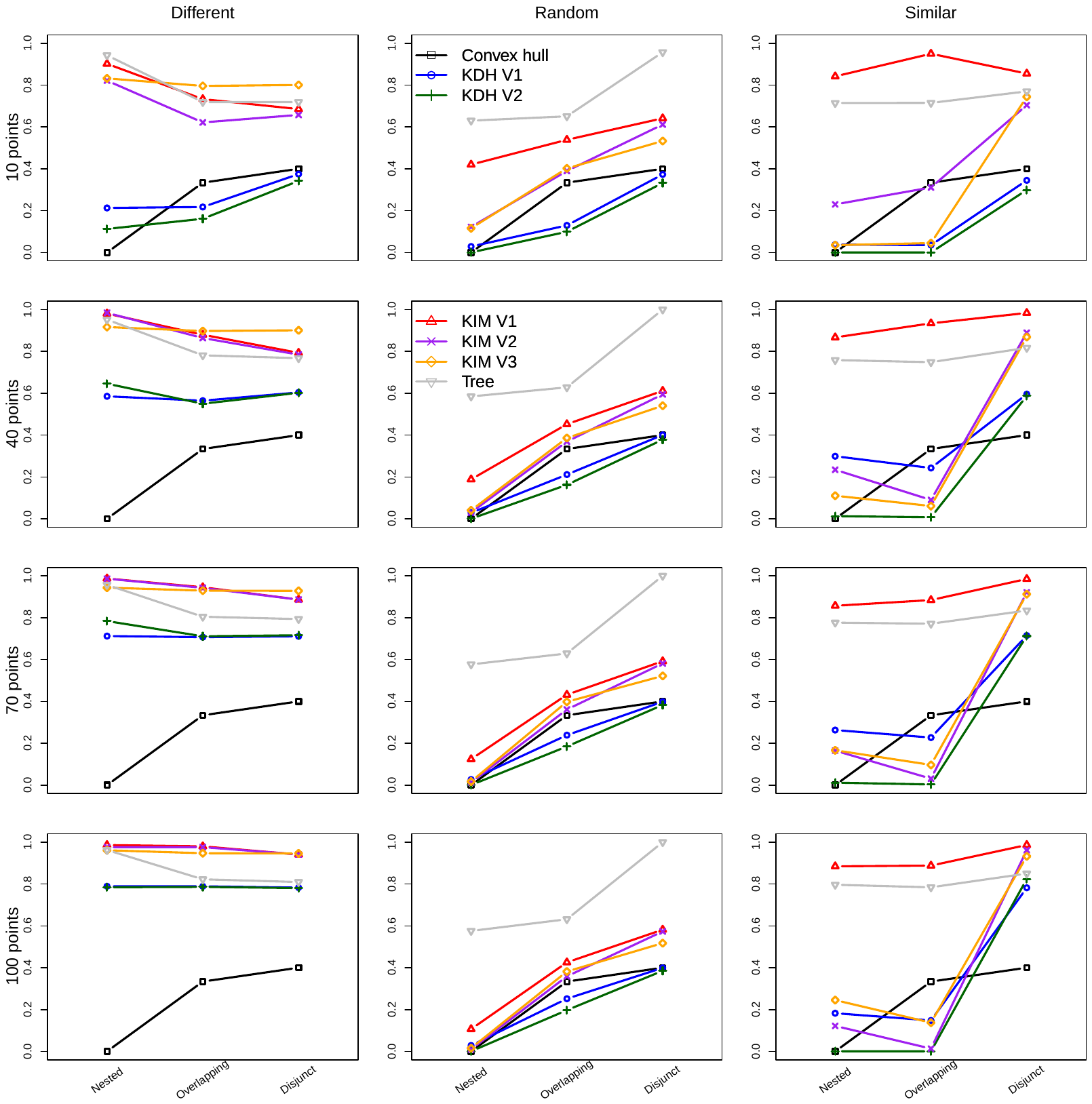

**Figure C9.** Changes in the contribution of replacement to overall turnover, computed as Williams replacement divided by Jaccard dissimilarity, for the seven approaches summarised in Table 1, for MCPs of different sizes. These are the same results as Figure C3, ordered differently.

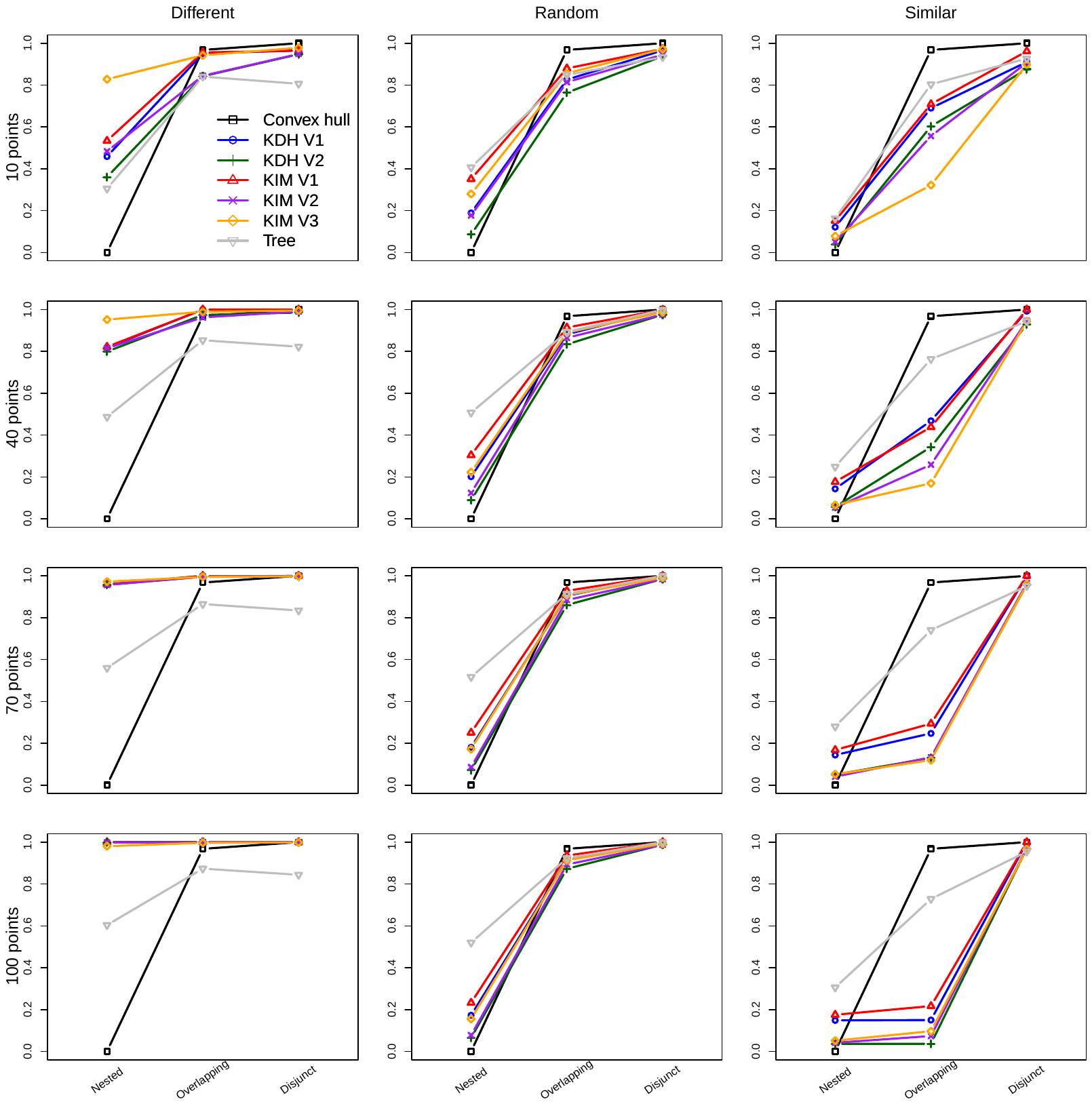

**Figure C10.** Changes in Jaccard dissimilarity for the seven approaches summarised in Table 1, for MCPs of the same size. These are the same results as Figure C4, ordered differently.

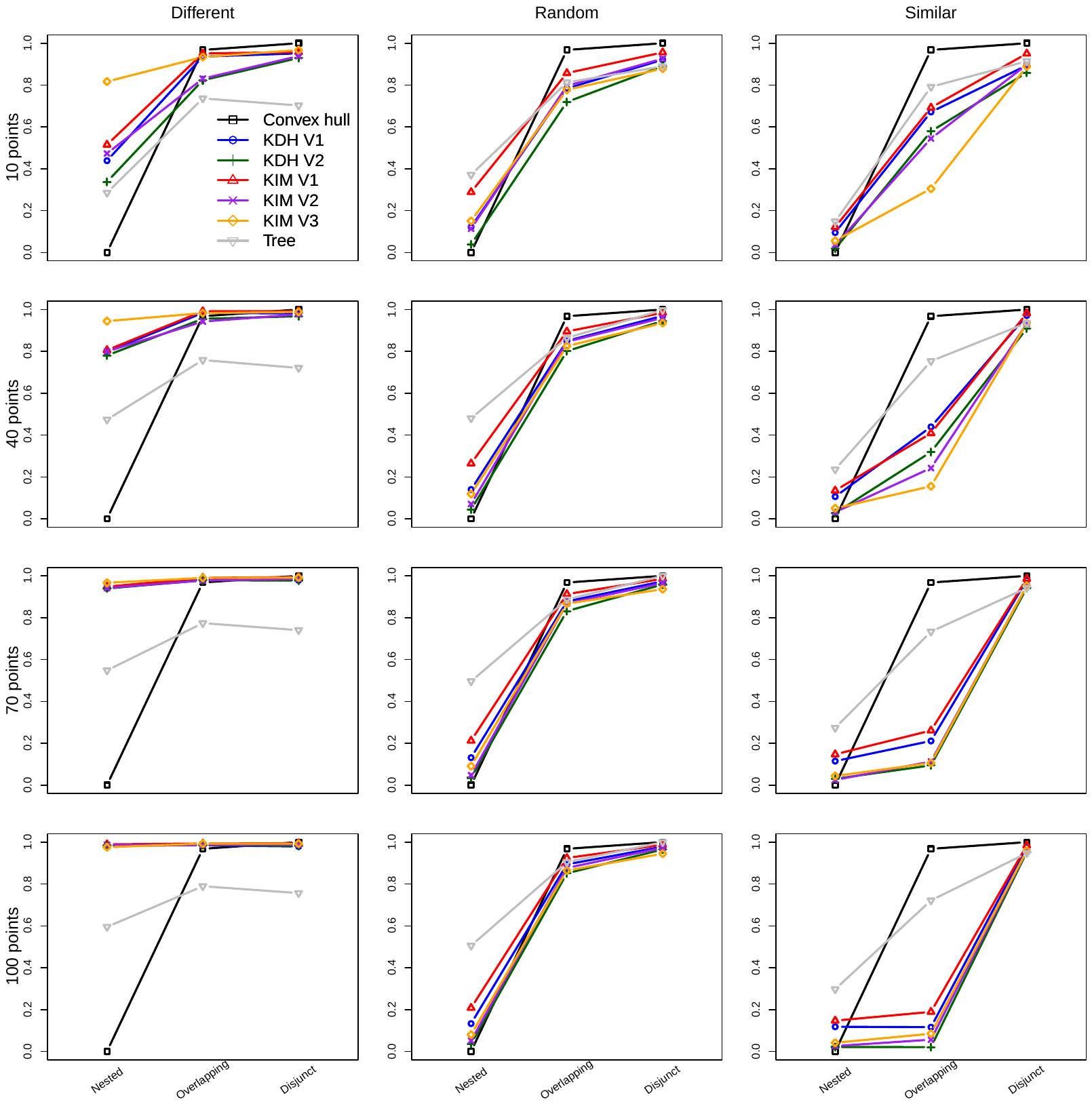

**Figure C11.** Changes in Williams replacement index for the seven approaches summarised in Table 1, for MCPs of the same size. These are the same results as Figure C5, ordered differently.

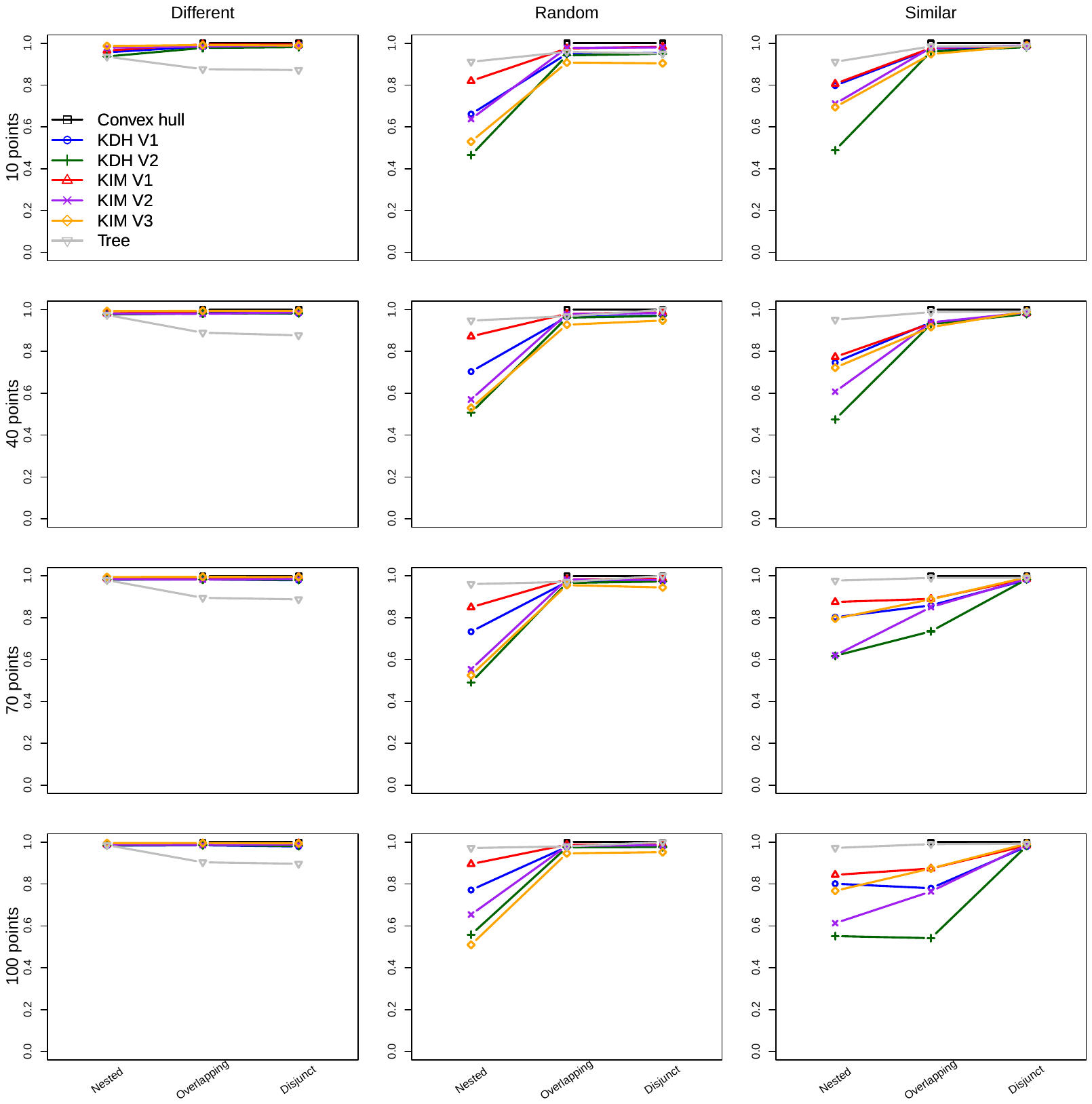

**Figure C12.** Changes in the contribution of replacement to overall turnover, computed as Williams replacement divided by Jaccard dissimilarity, for the seven approaches summarised in Table 1, for MCPs of the same size. These are the same results as Figure C6, ordered differently.

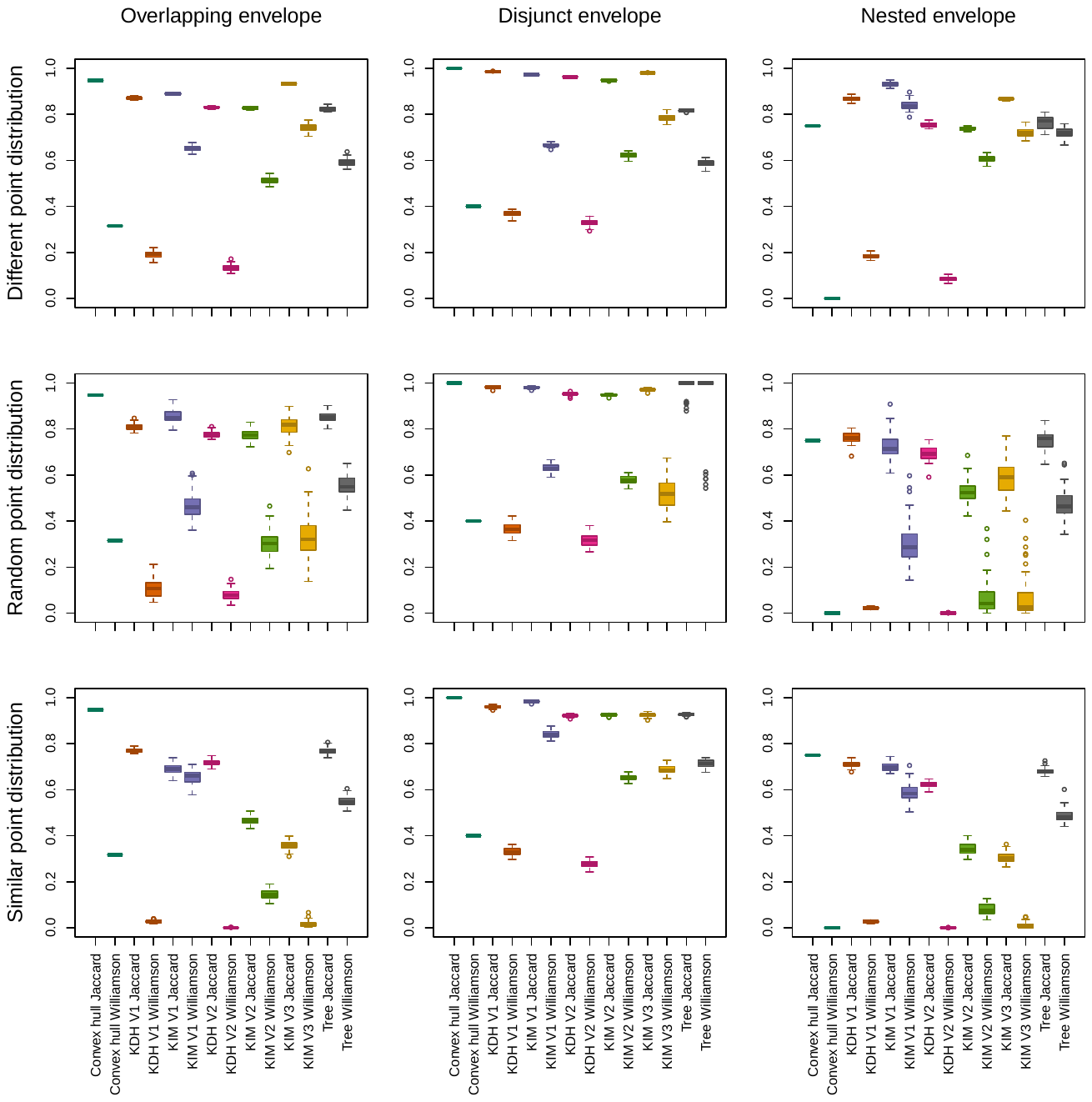

**Figure C13.** Boxplots of the values generated by the different approaches for the Jaccard dissimilarity and the Williams replacement indices, for MCPs of different sizes and 10 points.

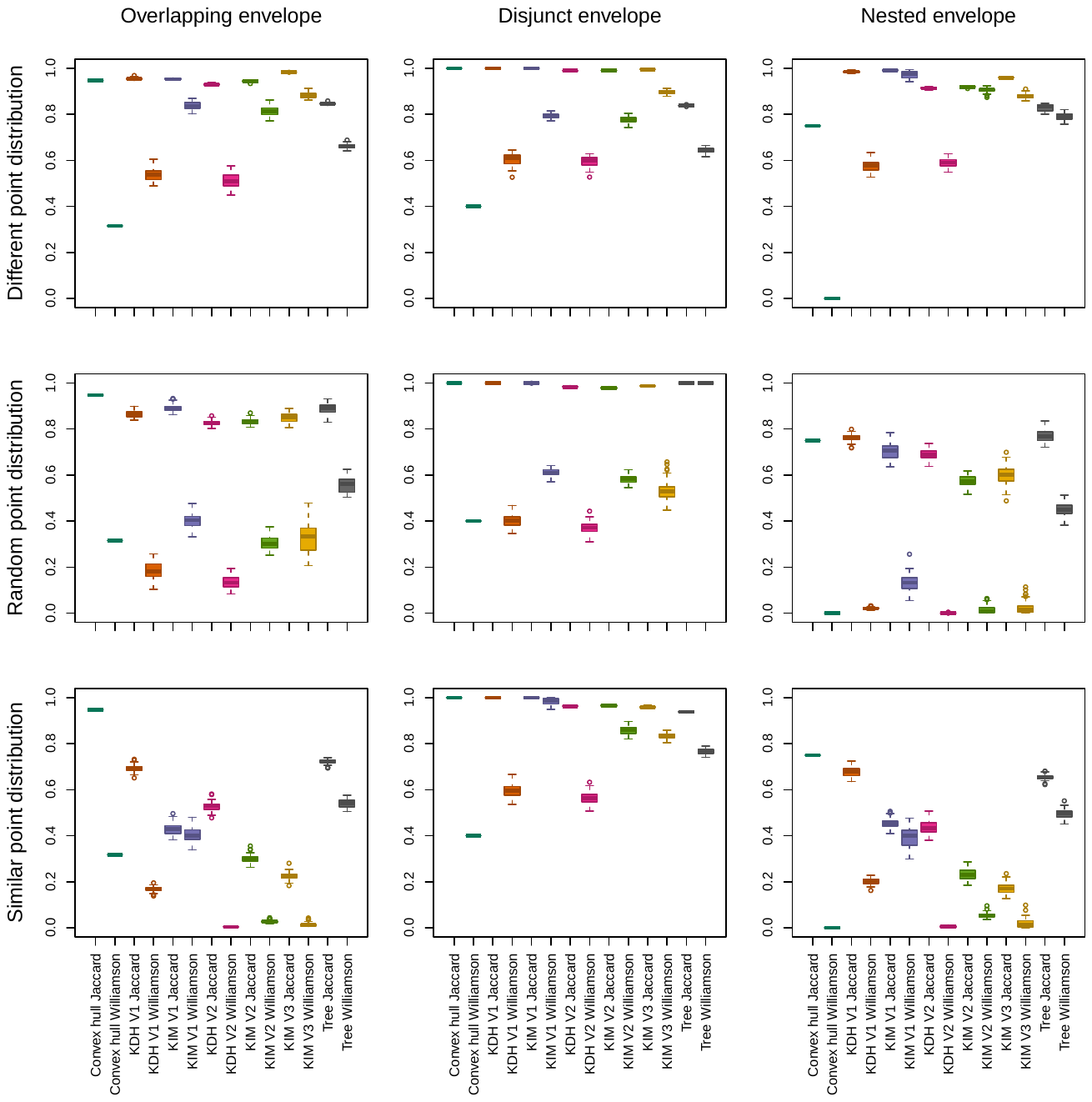

**Figure C14.** Boxplots of the values generated by the different approaches for the Jaccard dissimilarity and the Williams replacement indices, for MCPs of different sizes and 40 points.

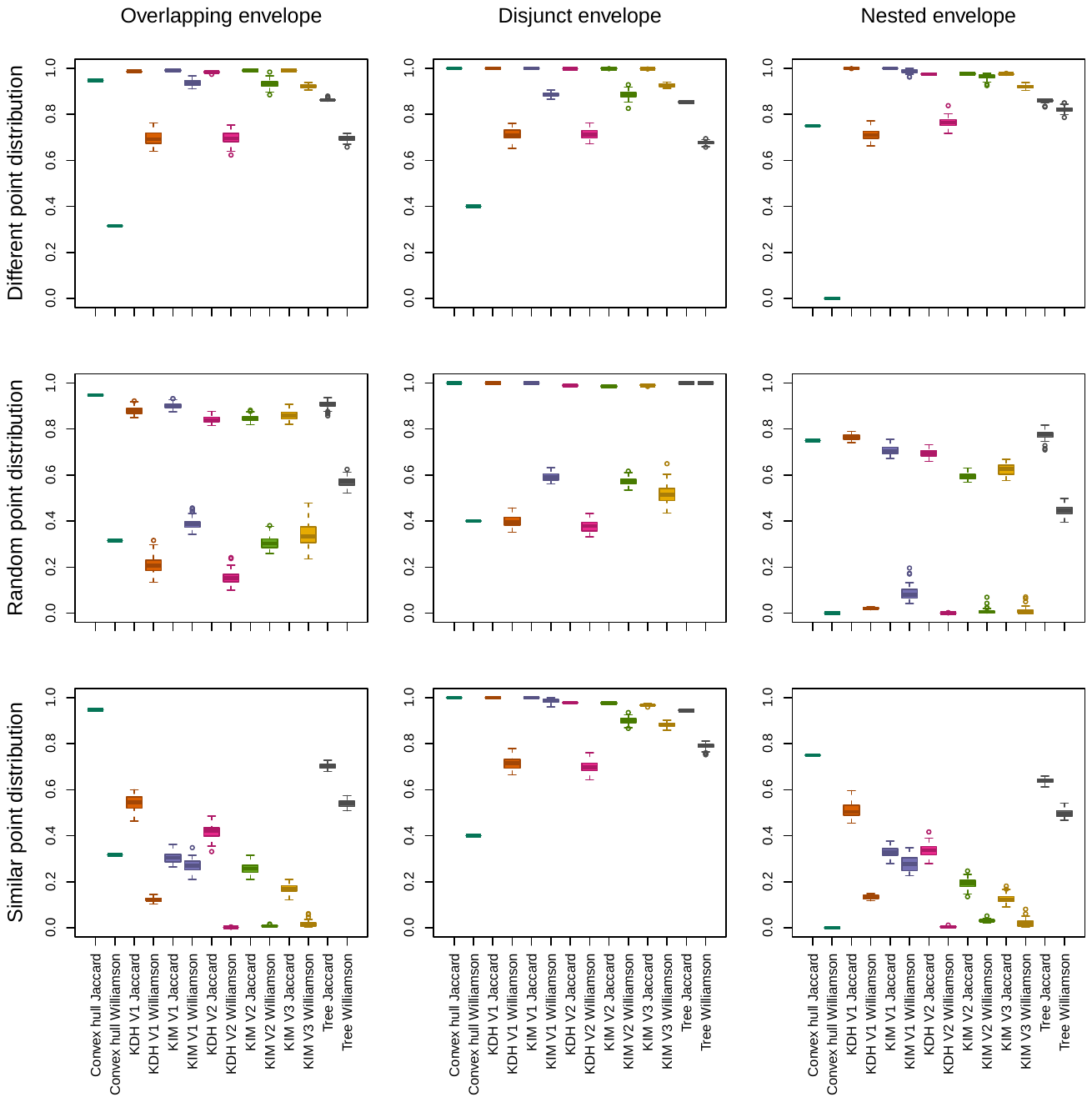

**Figure C15.** Boxplots of the values generated by the different approaches for the Jaccard dissimilarity and the Williams replacement indices, for MCPs of different sizes and 70 points.

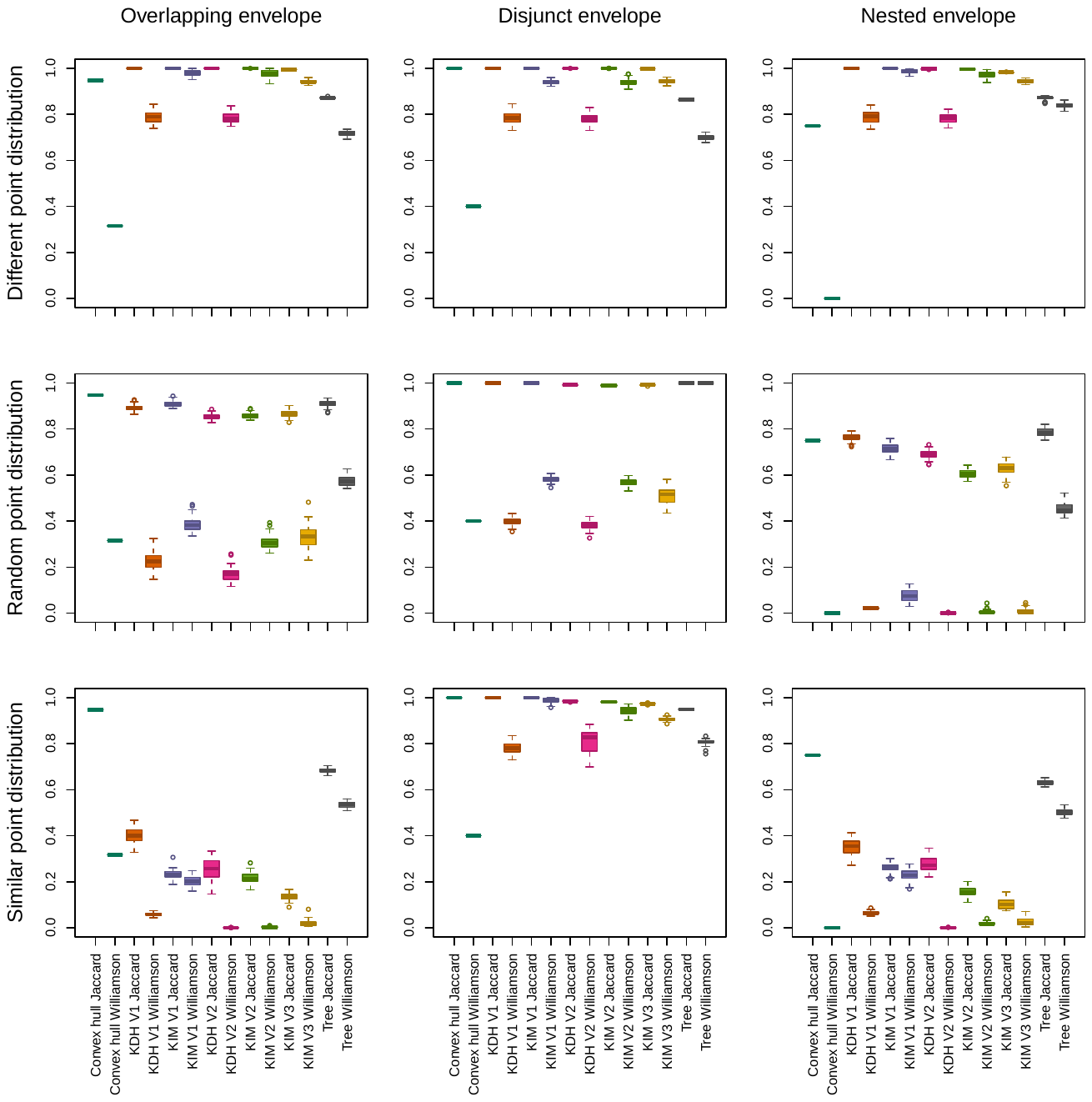

**Figure C16.** Boxplots of the values generated by the different approaches for the Jaccard dissimilarity and the Williams replacement indices, for MCPs of different sizes and 100 points.

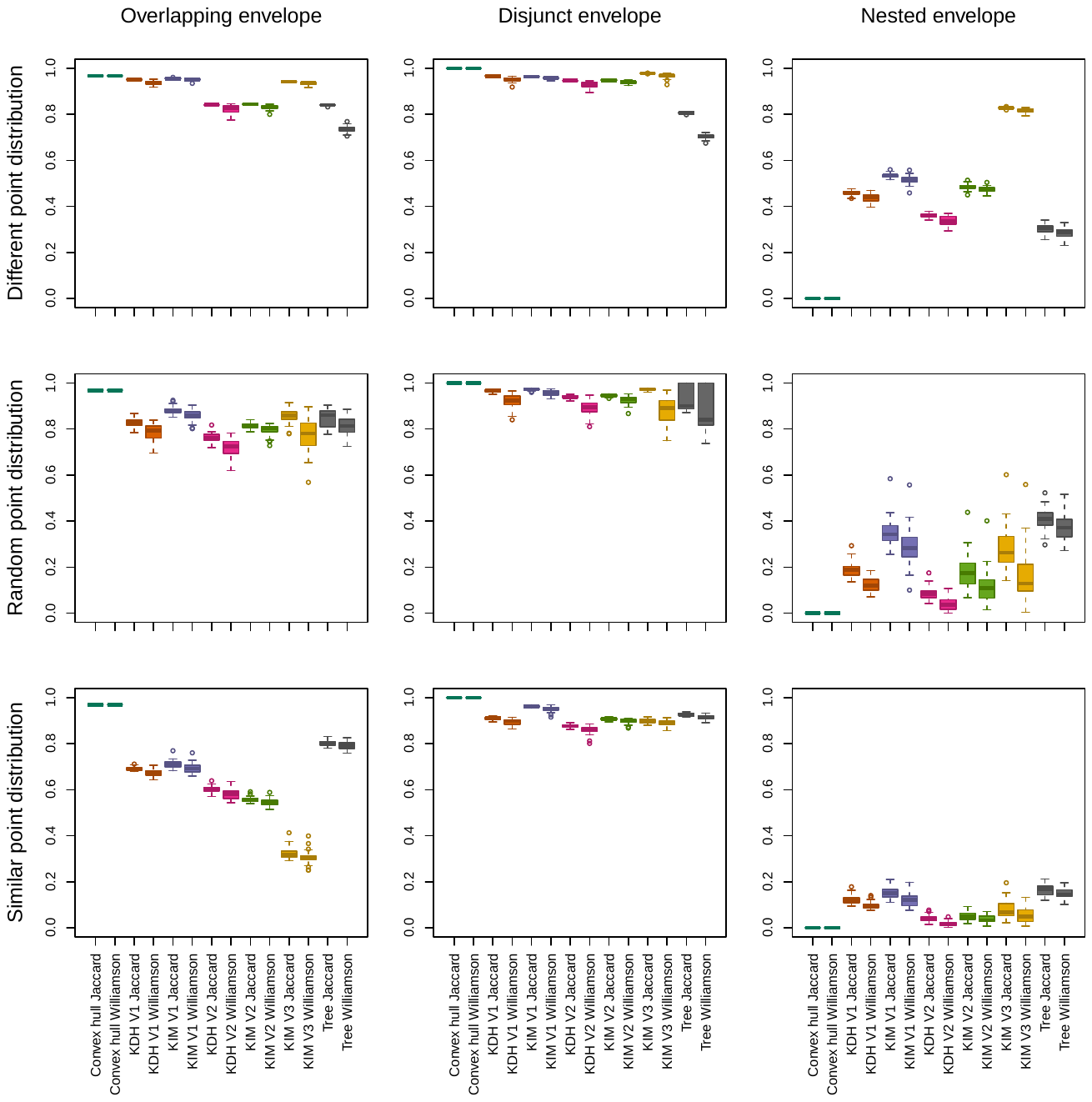

**Figure C17.** Boxplots of the values generated by the different approaches for the Jaccard dissimilarity and the Williams replacement indices, for MCPs of the same size and 10 points.

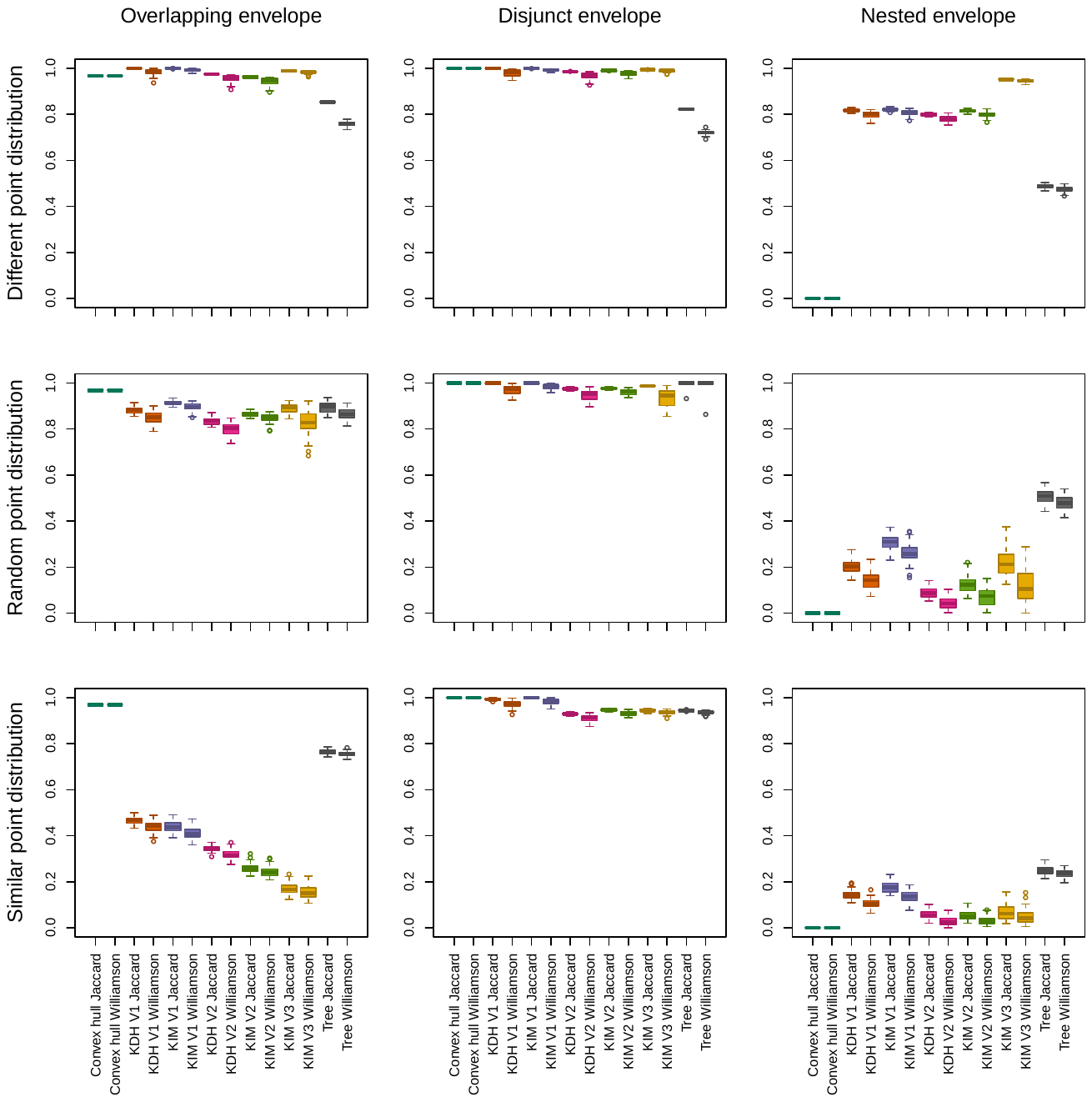
 **Figure C18.** Boxplots of the values generated by the different approaches for the Jaccard dissimilarity and the Williams replacement indices, for MCPs of the same size and 40 points.

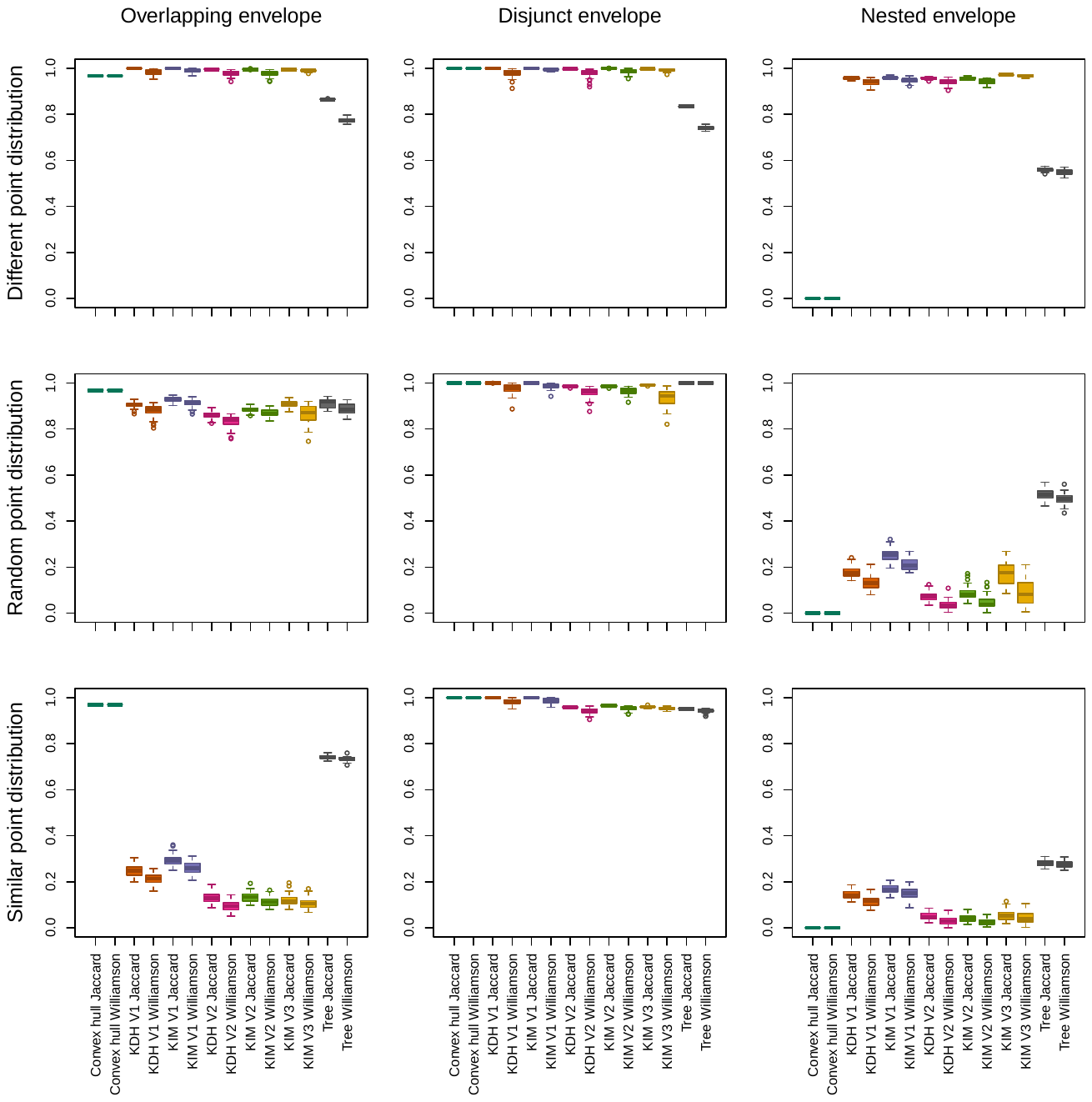

**Figure C19.** Boxplots of the values generated by the different approaches for the Jaccard dissimilarity and the Williams replacement indices, for MCPs of the same size and 70 points.

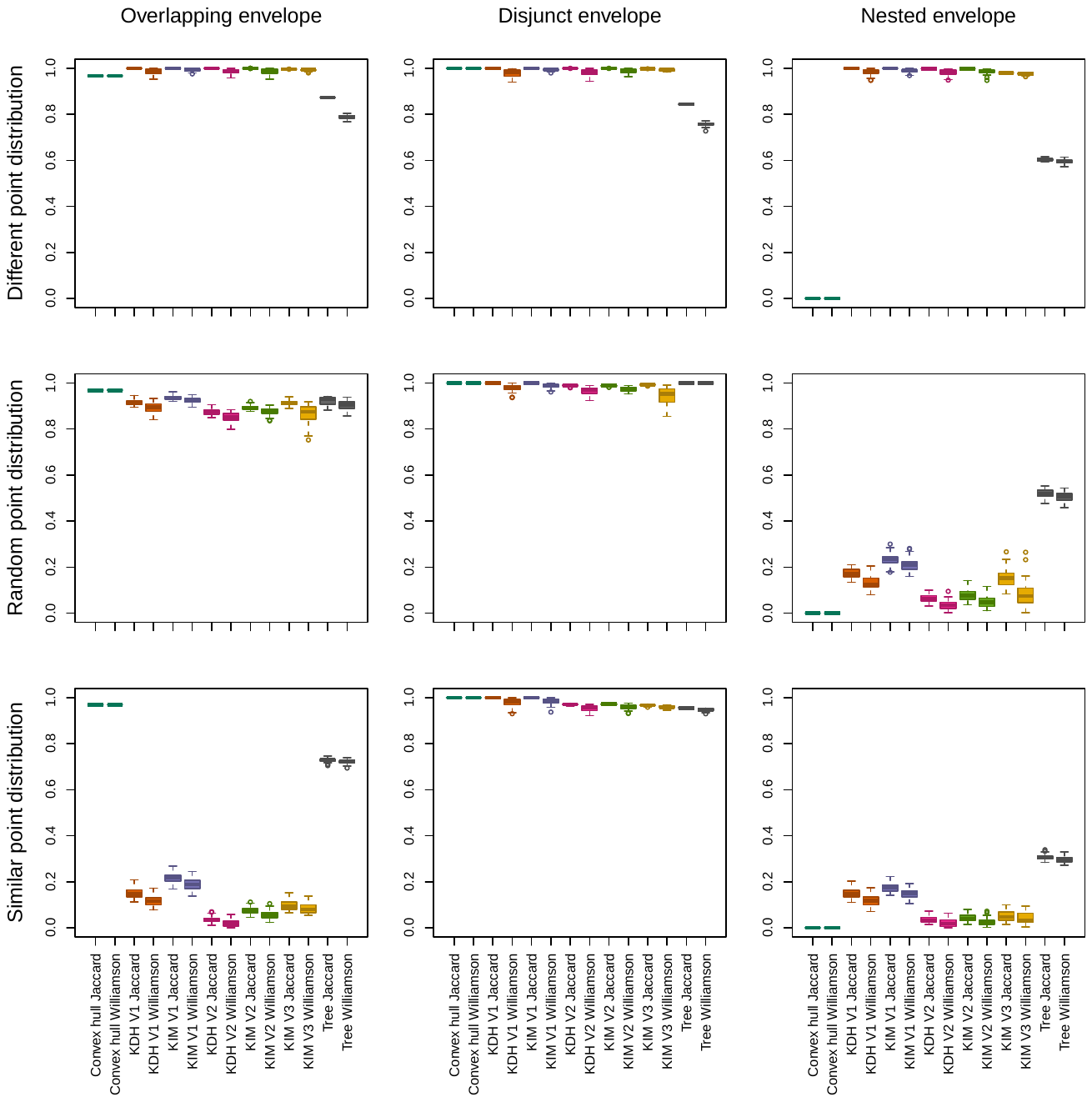

**Figure C20.** Boxplots of the values generated by the different approaches for the Jaccard dissimilarity and the Williams replacement indices, for MCPs of the same size and 100 points.

**Appendix D.** Supplementary results for French Polynesia data

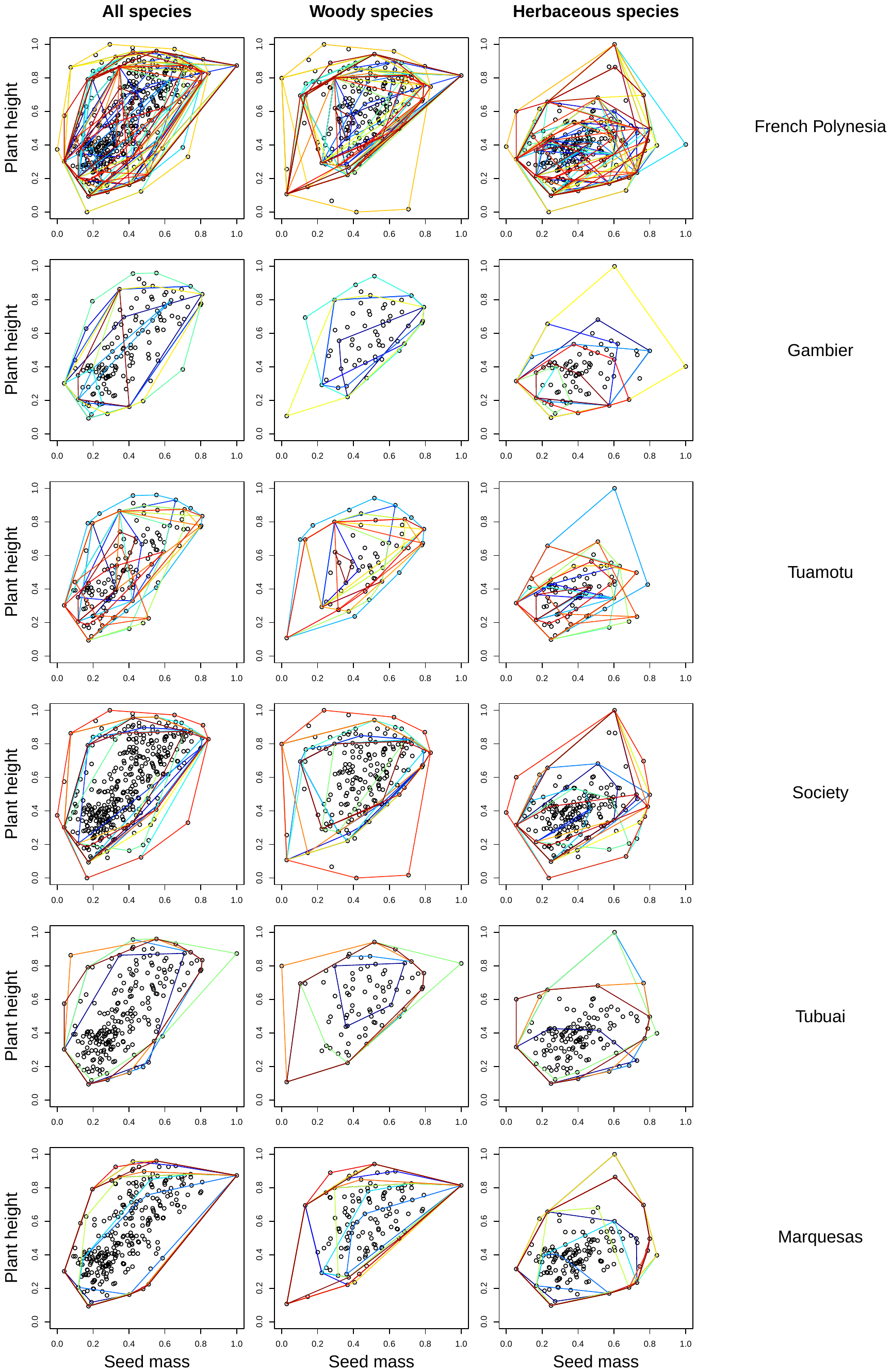

**Figure D1.** Naturalised alien plant species in French Polynesian islands and MCPs of each island in the functional space defined by seed mass and plant height, for each archipelago.

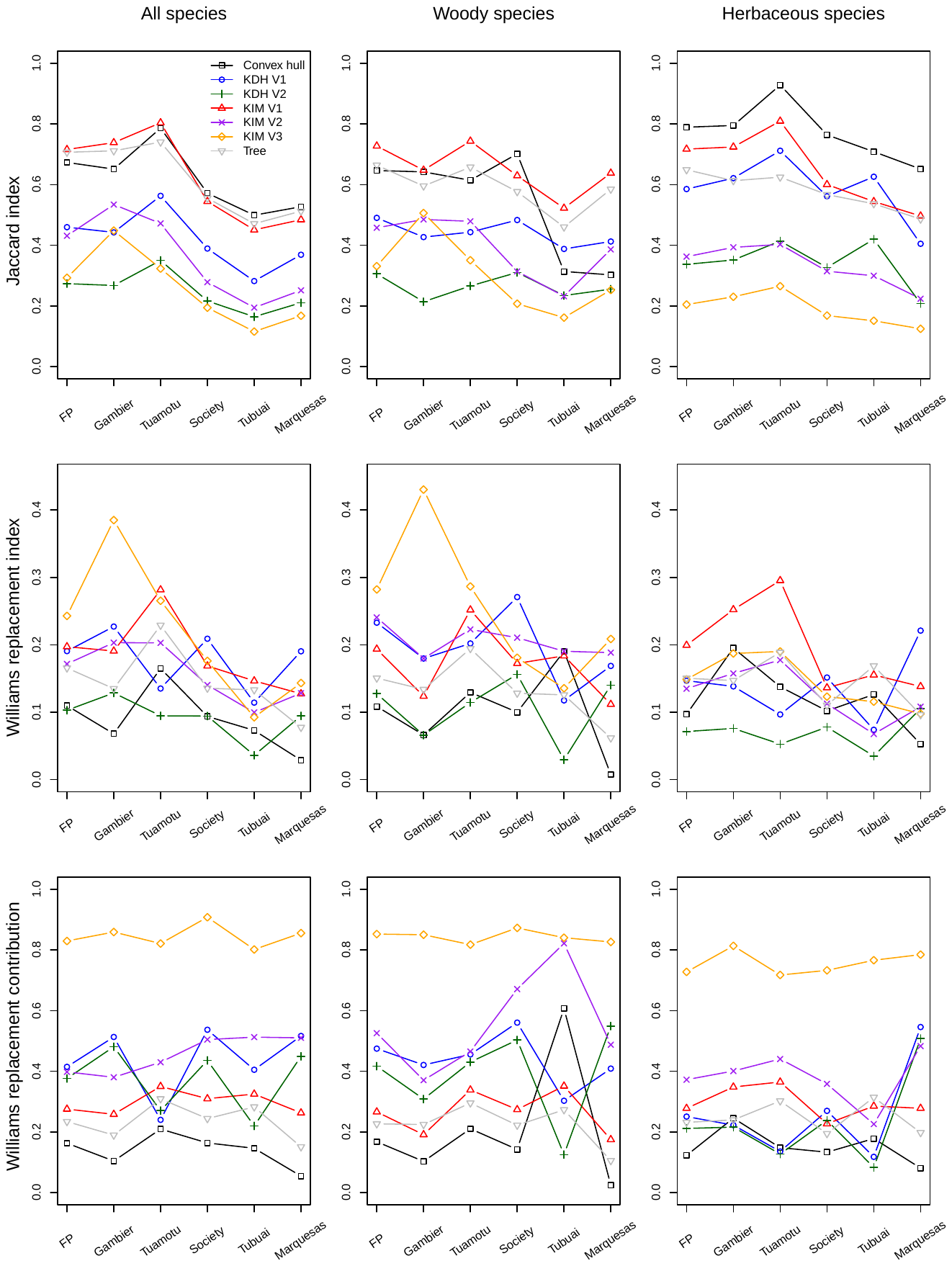

**Figure D2.** Jaccard dissimilarity index, Williams replacement index, and contribution of replacement to overall turnover, computed as Williams replacement divided by Jaccard dissimilarity, for the seven approaches summarised in Table 1, for French Polynesia and its archipelagos, for all species, woody species and herbaceous species, using seed mass, plant height and SLA, for 124 out of 417 species.

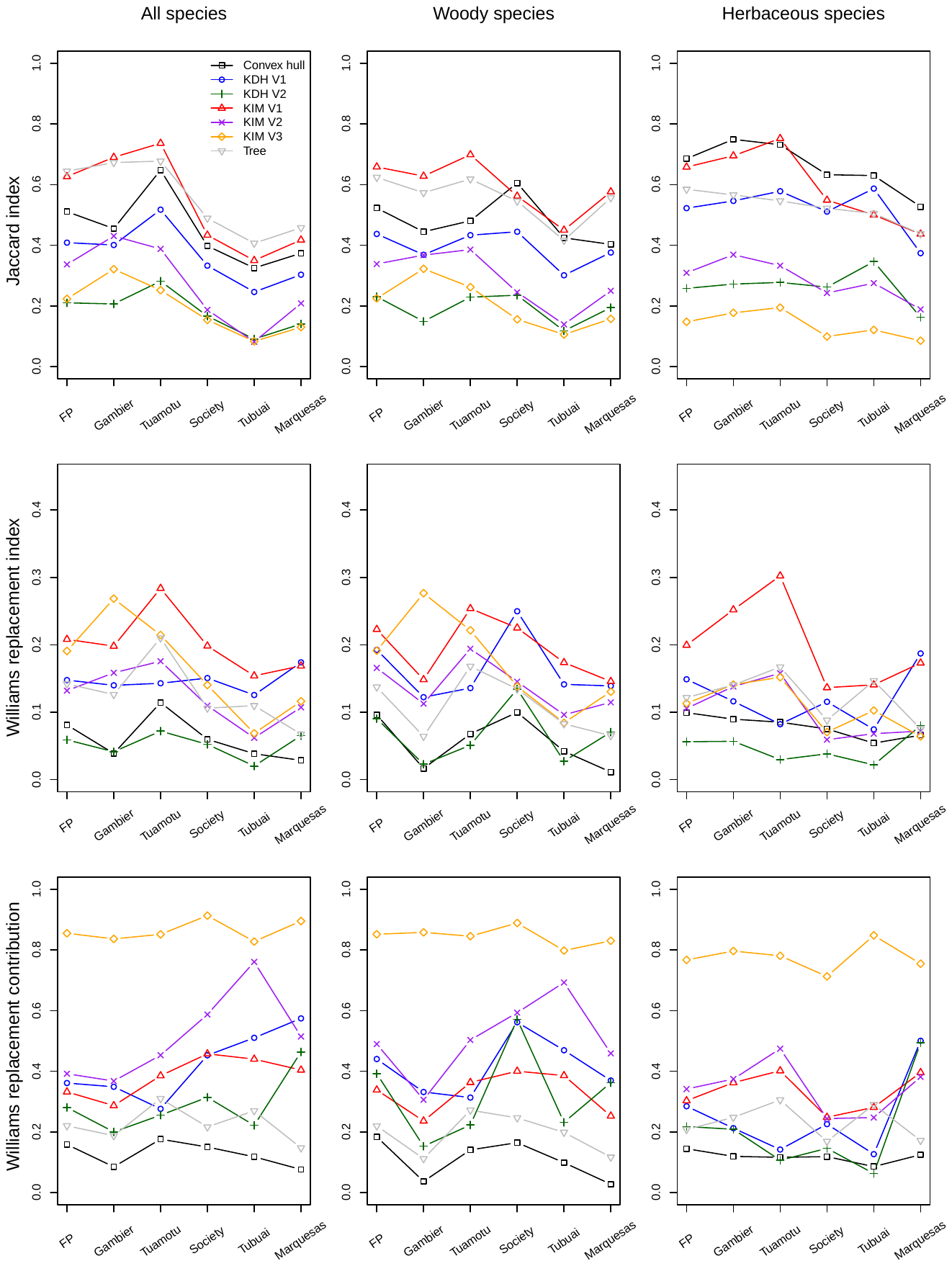

**Figure D3.** Jaccard dissimilarity index, Williams replacement index, and contribution of replacement to overall turnover, computed as Williams replacement divided by Jaccard dissimilarity, for the seven approaches summarised in Table 1, for French Polynesia and its archipelagos, for all species, woody species and herbaceous species, using seed mass and plant height, for 124 out of 417 species.

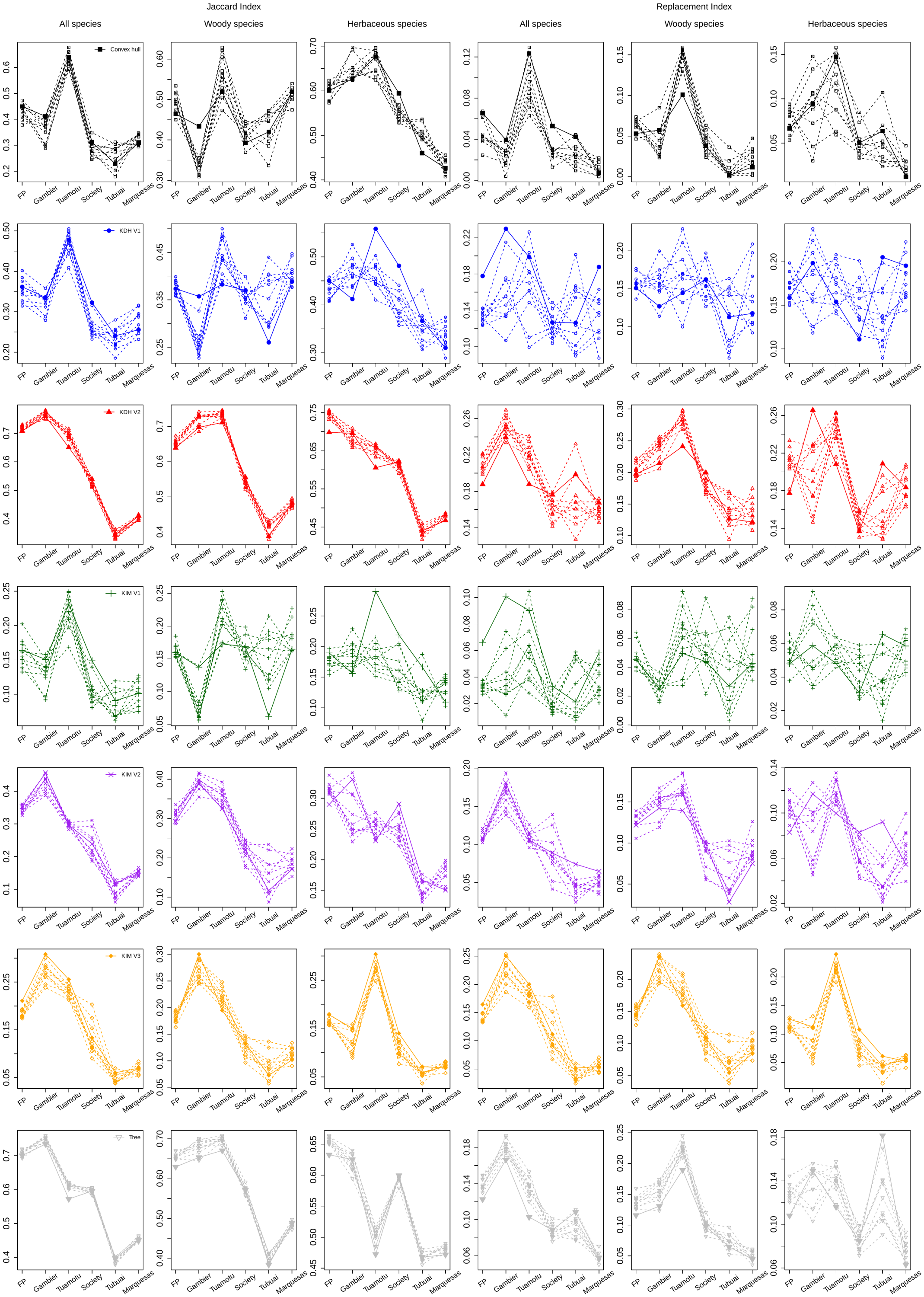

**Figure D4.** Jaccard dissimilarity index and Williams replacement index for the seven approaches summarised in Table 1, for French Polynesia and its archipelagos, for all species, woody species and herbaceous species, using seed mass and plant height, for 425 out of 417 species. Solid lines are observed values, while dashed lines are values from randomised presence-absence matrices.
